# Strong Patterns of Intraspecific Variation and Local Adaptation in Great Basin Plants Revealed Through a Review of 75 Years of Experiments

**DOI:** 10.1101/506089

**Authors:** Owen W. Baughman, Alison C. Agneray, Matthew L. Forister, Francis F. Kilkenny, Erin K. Espeland, Rob Fiegener, Matthew E. Horning, Richard C. Johnson, Thomas N. Kaye, Jeffrey E. Ott, J. Bradley St. Clair, Elizabeth A. Leger

**Author notes:** Corresponding author, current address: The Nature Conservancy, 67826-A Hwy. 205, Burns, OR 97720 USA.

## Abstract

Variation in natural selection across heterogeneous landscapes often produces 1) among-population differences in phenotypic traits, 2) trait-by-environment associations, and 3) higher fitness of local populations. Using a broad literature review of common garden studies published between 1941 and 2017, we documented the commonness of these three signatures in plants native to North America’s Great Basin, an area of extensive restoration and revegetation efforts, and asked which traits and environmental variables were involved. We also asked, independent of geographic distance, whether populations from more similar environments had more similar traits. From 327 experiments testing 121 taxa in 170 studies, we found 95.1% of 305 experiments reported among-population differences, and 81.4% of 161 experiments reported trait-by-environment associations. Locals showed greater survival in 67% of 24 reciprocal experiments that reported survival, and higher fitness in 90% of 10 reciprocal experiments that reported reproductive output. A meta-analysis on a subset of studies found that variation in eight commonly-measured traits was associated with mean annual precipitation and mean annual temperature at the source location, with notably strong relationships for flowering phenology, leaf size, and survival, among others. Although the Great Basin is sometimes perceived as a region of homogeneous ecosystems, our results demonstrate widespread habitat-related population differentiation and local adaptation. Locally-sourced plants likely harbor adaptations at rates and magnitudes that are immediately relevant to restoration success, and our results suggest that certain key traits and environmental variables should be prioritized in future assessments of plants in this region.

## Introduction

All plant species have limits to the range of conditions in which they can live, and all but the narrowest endemics grow across environments that vary in biotic and abiotic conditions. This natural complexity has significant impacts on individual survival and reproduction, and thus plant evolution (Loveless and Hamrick, 1984; Linhart and Grant, 1996; Ackerly *et al.*, 2000; Reich *et al.*, 2003). As plants are subject to different conditions associated with their local environment, populations of the same species will experience differential selection pressures (Turesson, 1922; Clausen, Keck and Hiesey, 1948; Antonovics and Bradshaw, 1968; Langlet, 1971), creating habitat-correlated intraspecific variation. When this intraspecific variation results in populations that are more fit in their home environment than foreign populations, these populations are considered to be locally adapted (Kawecki and Ebert, 2004; Blanquart *et al.*, 2013). The existence of local adaptation is well-established across different organisms and ecosystems, although our synthetic knowledge of this important topic rests on surprisingly few reviews of the subject (e.g. Leimu and Fischer, 2008; Hereford, 2009; Oduor, Leimu and van Kleunen, 2016). Here, we focus on a particular region and ask if plant species share patterns of intraspecific variation and local adaptation, and, across taxa, what functional traits and environmental variables are most important for such patterns in this region. The regional focus provides a strong test of expectations generated from more heterogeneous samples, facilitates comparison of the strength of selection among specific traits, and provides an opportunity to link basic evolutionary patterns with applied concerns.

The detection of local adaptation ideally involves reciprocal transplant experiments designed to test for a local advantage across environments (Blanquart *et al.*, 2013; Bucharova, Durka, *et al.*, 2017). However, patterns associated with local adaptation (hereafter, signatures) can be detected in non-reciprocal comparisons of different populations of the same species (Endler, 1986). When populations are locally adapted to environmental variables, we expect to see three basic signatures from common garden experiments: 1) differences among populations in fitness-related traits, 2) correlations between these trait values and environmental or other habitat-related variables, and, if reciprocal transplants have been conducted, 3) higher fitness of local over nonlocal populations in the local environment. Although population differences (signature 1) are necessary for local adaptation, they alone are not sufficient evidence due to factors such as genetic drift, high gene flow, and rapid environmental change, among other factors (Kawecki and Ebert, 2004; Blows and Hoffmann, 2005). While fitness differences in reciprocal transplant experiments (signature 3) are the “gold standard” for detecting local adaptation, there are experimental trade-offs between the number of populations sampled and the ability to do fully reciprocal transplants (Blanquart *et al.*, 2013). Thus, correlative approaches (signature 2) are popular alternatives that can sample many more populations to infer local adaptation (e.g. St Clair, Mandel and Vance-Borland, 2005), though spurious correlations, low sample sizes, or high variability in trait values could over- or under-predict the degree of local adaptation in wild populations using this approach. Given these considerations, separately reporting all three signatures can give an overall picture of the likelihood of within-species variation and potential local adaptation in a region, and is the first step towards a better understanding of variation in the strength and consistency of natural selection (Siepielski, Dibattista and Carlson, 2009).

The Great Basin Desert of North America is a ∼540,000 km^2^ cold desert landscape characterized by hundreds of internally-draining basin and range formations, which create high spatial and environmental heterogeneity and variability (Tisdale and Hironaka, 1981; Comstock and Ehleringer, 1992). While these are the kinds of conditions that would be expected to result in widespread local adaptation, the flora of the Great Basin is poorly represented in the relatively few reviews on the subject (Leimu and Fischer, 2008; Hereford, 2009; Oduor, Leimu and van Kleunen, 2016), and this has resulted in uncertainty as to the prevalence, magnitude, and importance that local adaptation plays in this large and increasingly imperiled region (United States. House of Representatives. Committee on Appropriations., 2014; Jones, Monaco and Rigby, 2015; Chivers *et al.*, 2016). Gaining a better understanding of local adaptation in the Great Basin is important not only because it is a large, relatively intact floristic region in the Western US, but also because this information has direct impacts on conservation and restoration efforts. Large-scale, seed-based restoration has been very common in the Great Basin for many decades (Pilliod, Welty and Toevs, 2017), and trends in large destructive wildfires (Dennison *et al.*, 2014) and other disturbances (Rowland, Suring and Michael, 2010; Davies *et al.*, 2011) ensure even higher demand for restoration efforts in the future. Guided by the various national policies and strategies dating from the 1960s (Richards, Chambers and Ross, 1998) to the present National Seed Strategy (Plant Conservation Alliance, 2015) and Integrated Rangeland Fire Management Strategy (USDOI, 2015), a growing majority of these efforts are using native plants. However, few of the widely-available sources of commercially-produced seeds of native species originate from populations within the Great Basin (Jones and Larson, 2005) or have been selected based on their success in restoring Great Basin habitats (Leger and Baughman, 2015). Further, demand for native seed has always exceeded supply (McArthur and Young, 1999; Johnson *et al.*, 2010), which has resulted in the prioritization of seed quantity and uniformity over population suitability and local adaptation (Meyer, 1997; Richards, Chambers and Ross, 1998; Leger and Baughman, 2015). Therefore, it is still uncommon for restorationists in this region to prioritize or even have the option to prioritize the use of local populations, despite growing support of the importance of such practices (Basey, Fant and Kramer, 2015; Espeland *et al.*, 2017).

Though our understanding of the prevalence and scale of local adaptation in the Great Basin is far from complete, there is an abundant literature of peer-reviewed studies on the plants native to this region spanning over 75 years that have directly measured trait variation between populations via laboratory, greenhouse, or field common gardens and reciprocal transplants. Many of these studies have also tested for correlations between intraspecific variation and environmental variables, and some were designed to detect local adaptation. This research includes studies of germination patterns (e.g. McArthur, Meyer and Weber, 1987; Meyer *et al.*, 1995), large genecology experiments (e.g. Erickson, Mandel and Sorenson, 2004; Johnson, Leger and Vance-Borland, 2017), and reciprocal transplants (e.g. Evans and Young, 1990; Barnes, 2009), among other types of studies. This rich literature provides an opportunity to summarize local adaptation and its associated patterns, or signatures (defined above), in this region, as well as describe which phenotypic traits have the strongest signatures of local adaptation.

Here, we present results of a broad literature review and subsequent meta-analysis using published studies that compared phenotypic traits of multiple populations of native Great Basin species in one or more common environments. Our first objective was to record published instances of the three expected signatures of local adaptation (population variation, trait-by-environment association, and greater local fitness) within grasses, forbs, shrubs, and deciduous trees native to the Great Basin, asking how common these signatures are, as well as which phenotypic traits and environmental variables were most commonly associated with these signatures. We also present results by taxonomic group, lifeform, lifespan, distribution, and mating system. This first objective encompassed all possible studies, including those that did not provide sufficient details for formal meta-analysis, which allowed us to incorporate the broadest range of studies, including older studies that provided minimal quantitative detail. Our second objective was to examine links between the magnitude of trait and environmental divergence (mean annual precipitation and mean annual temperature) among populations across multiple taxa, for the subset of experiments amenable to this approach, asking whether populations from more similar environments were more similar in phenotypic traits. We also used meta-analysis to ask which traits and environmental variables showed the strongest patterns of association.

We expected to find widespread evidence of local adaptation and its signatures in the plants of the Great Basin, and we hypothesize that phenological and size-based traits, which show phenotypic variation in response to climate variation in both plants and animals (e.g. Sheridan and Bickford, 2011; Anderson *et al.*, 2012) and have been observed to be under selection in the Great Basin (Leger and Baughman, 2015), would be important indicators of adaptation in this region. We discuss our results both as a contribution to our general understanding of natural selection in plants, and as an example of evolutionary theory applied to the management and restoration of a large geographic region, where active and ongoing management can benefit from information on intraspecific variation and local adaptation.

## Methods

### Literature search

We began by using the search engines Google Scholar and Web of Science to search for combinations of key terms (see additional methods in Supporting Information Appendix 1). In order to be included in our review, a study had to meet all these criteria:

a. Examined a species that is native within the floristic Great Basin
b. Examined and compared more than one population of that species
c. Measured at least one phenotypic, physiological, phenological, or other potentially fitness-related trait (e.g. survival; hereafter, trait)
d. Measured the trait(s) of the populations in at least one common environment (including laboratories, growth chambers, greenhouses, or outside gardens; hereafter, garden).

A plant was determined to be native to the Great Basin if the taxa had at least one occurrence with native status within the floristic Great Basin according to occurrence information from the USDA Plants Database (USDA and NRCS, 2018) and/or the U.S Virtual Herbarium Online (Barkworth *et al.*, 2018). A total of 170 studies published between 1941 and July 2017 were encountered that met these criteria.

### Categorization and scoring of literature

All studies meeting our criteria were categorized and scored for each signature. The coordinates of all gardens and populations in each study were recorded or, if possible, generated from localities described in the studies (Supporting Information Appendix 1). For each study, we then noted these 15 characteristics: the year published, year(s) of plant material collection, year(s) of experimentation, number of years reported, taxa (genus, species, subspecies), life history traits (taxonomic status, lifeform, geographic range, life span, breeding system), experiment type (laboratory, greenhouse, common garden, reciprocal transplant), number of gardens, number of populations tested, which generation of material was used, and whether or not experimenters attempted to control for maternal effects prior to testing (Supporting Information Appendix 1). Life history traits were compiled for each taxon from the USDA Plants Database as well as from published literature (Supporting Information Appendix 1). Each taxon (subspecies level, if given) was entered separately for studies addressing multiple taxa. In studies where more than one experiment was performed, and the experiments differed in the experiment type (defined above), the identity of the populations being compared, and/or the generation of material used, they were entered as separate experiments. In cases where the list of tested populations was identical among multiple published studies, and these materials came from the same collections, these experiments were entered separately if the garden type or location(s) differed among the studies or if authors separately published different traits from the same gardens, ensuring that no trait was recorded twice for the same set of populations in the same garden. In cases where the list of tested populations did not completely overlap between studies, even if some from each study arose from the same collections, they were entered separately. These methods carefully emphasized the inclusion of the greatest number of relevant experiments and traits without duplication, but nonetheless resulted in some non-independence between some experiments. A total 327 taxa-specific entries (hereafter, experiments) were generated from the 170 published studies (Supporting Information Appendix 2).

The first two expected signatures of local adaptation were scored using a Yes/No designation for each experiment which considered all measured phenotypic traits. A score of “Yes”, or, in the absence of supporting statistical evidence, “Authors claim Yes”, was given when at least one measured trait significantly demonstrated the signature for at least two populations, and a score of “No” or “Authors claim No” was given when the signature was not detected between any pair of populations (Supporting Information Appendix 1). In addition, each of the measured and reported traits and environmental variables were scored (hereafter, trait scores) in the same way for each signature. Of the 327 experiments, 305 (93.3%) met the criteria to score for among-population variation (signature 1) and 161 (49.5%) met the criteria to score for trait-by-environment association (signature 2). Pearson’s chi-squared tests were used to determine if there were differences in signatures 1 and 2 among plants with different life-history traits, using totals from both “Yes/No” and “Authors Claim Yes/No” results, excluding any life history groups represented by less than 10 experiments.

To score whether there was higher fitness of a local population in a common garden (hereafter, signature 3), only experiments in which outdoor reciprocal transplants or common gardens were performed using a local population in at least one garden were considered (Supporting Information Appendix 1). Additionally, the experiment had to measure a fitness-relevant response: survival, reproductive output (number of seeds or flowers, or other reproductive output), a fitness index (a combination of several size and production traits), or total aboveground biomass. Each experiment was assigned a composite score to fully capture variation in the performance of each garden’s local population, across multiple gardens as well as through multiple sampling dates (Supporting Information Appendix 1). The five possible composite scores were “Yes for all gardens at all times”, “Yes for all gardens at some times”, “Yes for some gardens at all times”, “Yes for some gardens at some times”, and “No for all gardens at all times”. These scores refer only to those gardens within each experiment that included their own local population. Of the 326 experiments, 27 (8.3%) were appropriate for this scoring. This scoring provides an estimate of the commonness of higher local fitness, but it is not a measure of the importance of the difference per se. For example, a fitness difference could occur uncommonly, but have a large impact on population trajectories (i.e. large differences in survival after a rare drought event).

Our dataset, which had uneven numbers of experiments representing each species, contained the possibility of bias associated with highly-studied taxa influencing patterns more than less-studied taxa. To ask how this affected overall results, we compared tallies of all scores without correcting for multiple experiments per species to tallies using an average score for each species for each signature. To generate these average scores for signature 1 and 2, we totaled all “Yes” and “Authors claim Yes” scores for each species and divided by the total number of scores (all Ys plus all Ns) for that species. For signature 3, all forms of “Yes” (all but “No for all gardens at all times”) were totaled into a Y and divided by the total number of scores. Then, we averaged these per-species scores to re-calculate overall effects in which each species was represented only once, and compared the results of the different averaging methods for each signature.

### Quantitative comparison of trait-by-environment associations

As a complement to the survey of author-reported results described above, we conducted a further, quantitative analysis of trait and climate values. Specifically, to examine associations between the differences in trait values and the differences in environmental and geographic distance among population origins, we utilized experiments from which population-specific trait data and geographic coordinates could be extracted or obtained through author contact. Data from laboratory and greenhouse experiments were not considered for this extraction. First, we identified the most commonly measured traits across studies, which were then manually extracted from text, tables, or graphical data (Supporting Information Appendix 1). Next, we extracted trait data from the latest sampling date for which the most populations at the most gardens were represented, and if multiple treatments were used, we only extracted data for the author-defined ‘control’ treatment. However, if no control was defined, we used the treatment that was the most unaltered or representative of the garden environment (e.g. unweeded, or unwatered). For each population/trait combination, we used either author-provided mean values or calculated a mean trait value from available data. Rather than averaging values across gardens, data, data from each garden location within each experiment was extracted separately and considered its own sample. We did this because it is not uncommon for traits to be expressed differently in different common garden locations (e.g. Johnson, Leger and Vance-Borland, 2017). Finally, we generated 30-year annual precipitation and mean annual temperature values for each population’s location of origin using the ClimateNA v5.10 software package based on methodology described by Wang et al. (2016). These 30-year averages are calculated every 10 years (i.e. 1951-1980, 1961-1990, etc.). Because studies took place at many times over the last 75 years, we used the most proximate climate normal for each experiment that did not include or surpass the years during which the experiment’s populations were collected (Supporting Information Appendix 2).

To reduce the likelihood of spurious correlations or false negative results, we limited this dataset to traits measured in at least 5 populations in at least 20 common garden locations (mean locations per trait: 34.4; range: 21-46), resulting in 81 locations (from 56 experiments) that measured at least one of eight frequently-measured phenotypic traits (Table 1). Within each location, we calculated pairwise Euclidean distances for each trait value, climate factor, and geographic distance for every possible pair of populations. Geographic distances were generated using the earth.dist function in fossil package (Vavrek, 2011) in the statistical computing environment R (R Core Team, 2017). Then, partial Mantel tests were used to compare pairwise trait and climate distances for each experiment while controlling for geographic distances, using the vegan package (Oksanen *et al.*, 2018) in R (R Core Team, 2017). We used the metacor.DSL function in the metacor package (Laliberté, 2011) to generate an overall effect size (partial correlation) and upper and lower confidence intervals for each combination of trait and environmental variable. Lastly, to better understand effect sizes for a subset of species, we ran simple linear regression analyses for each location, comparing average trait values and environmental values to generate a slope that estimated trait change per unit change in climate factors. Experiments with R^2^ values of 0.2 or less were excluded from this particular analysis, and the median slope across experiments was retained as an estimate of the trait-by-environment relationship. The arbitrary cutoff (R^2^ = 0.2) for this step was used simply as a way to focus on and report effect sizes from some of the stronger biological relationships that could be of particular interest to managers, restoration practitioners and evolutionary ecologists. Due to limited sample sizes for factors such as lifeform, mating system, geographic distribution, etc., we did not include these factors in any of the quantitative analyses, but present lifeform (shrub, grass, or forb) information for each trait response as additional results in the Supporting Information Appendix 3.

**Table 1.**
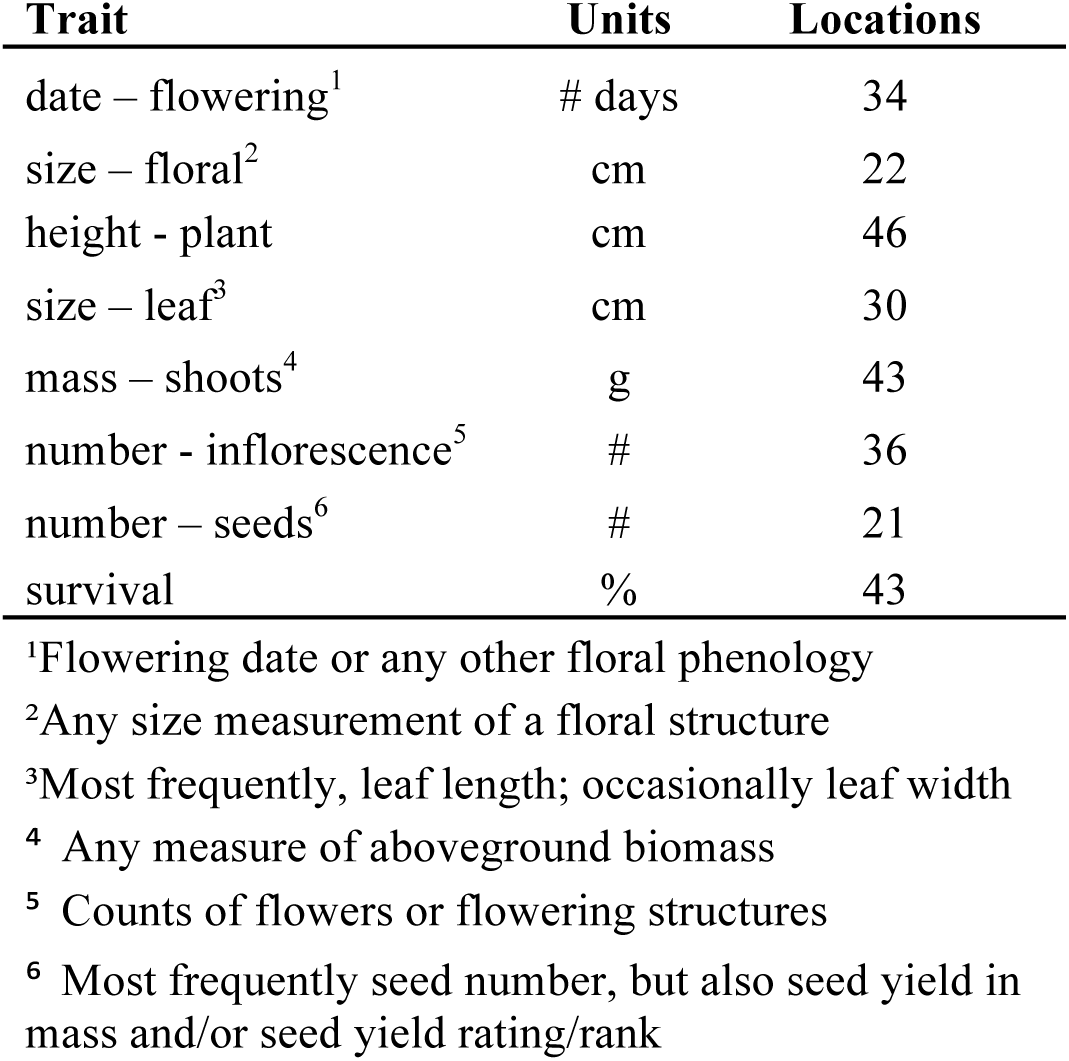
Traits measured in outdoor common gardens or reciprocal transplants for at least 5 populations in at least 20 common garden locations, with data available from text, tables, author contact, or extraction from figures. Note that in some cases, multiple highly similar measures were grouped, as indicated in footnotes.

| **Trait** | **Units** | **Locations** |
| --- | --- | --- |
| date – flowering<sup>1</sup> | # days | 34 |
| size – floral<sup>2</sup> | cm | 22 |
| height - plant | cm | 46 |
| size – leaf<sup>3</sup> | cm | 30 |
| mass – shoots<sup>4</sup> | g | 43 |
| number - inflorescence<sup>5</sup> | # | 36 |
| number – seeds<sup>6</sup> | # | 21 |
| survival | % | 43 |

<sup>1</sup>Flowering date or any other floral phenology
<sup>2</sup>Any size measurement of a floral structure
<sup>3</sup>Most frequently, leaf length; occasionally leaf width
<sup>4</sup> Any measure of aboveground biomass
<sup>5</sup> Counts of flowers or flowering structures
<sup>6</sup> Most frequently seed number, but also seed yield in mass and/or seed yield rating/rank

## Results

### Summary of reviewed literature

Our literature search revealed 170 published studies that measured trait responses from more than one population in at least one common environment, resulting in 327 separate experiments involving 121 taxa of 104 species of grasses, shrubs, forbs, and deciduous trees (Fig. 1). These experiments represent approximately 3,234 unique populations tested in approximately 208 outdoor garden locations (Fig. 2) and 154 indoor lab or greenhouse experiments. Grasses accounted for 21.0% of the taxa and 40.2% of the experiments, forbs composed 50.8% of the taxa and 30.7% of experiments, shrubs 26.6% of the taxa and 28.5% of experiments, and deciduous trees accounted for only 1.6% of taxa and 0.6% of experiments (Fig. 1A). Experiments were most commonly conducted in non-reciprocal outdoor common gardens (47.5%) or in the laboratory (31.9%), with fewer conducted in greenhouses (15.3%) or in reciprocal outdoor gardens (5.2%, Fig. 1B). For experiments in outdoor gardens, the median number of gardens per experiment across lifeform ranged from 1 (grasses, shrubs, and trees) to 2 (forbs) for non-reciprocal gardens, and from 2 (grasses and forbs) to 4 (shrubs) for reciprocal gardens. Overall, the median number of populations tested in each experiment was 5 (range= 2 - 193, IQR = 3 – 11.5, Fig. 1C), and was slightly lower for shrubs (median = 4, range = 2 – 111, IQR = 2 - 8) than grasses (median = 6, range = 2 – 193, IQR = 3 - 12.25), forbs (median = 6, range = 2 – 67, IQR = 3 – 10.25), and trees (median = 7, range = 5 – 9, IQR = 6 – 8).

**Figure 1.**
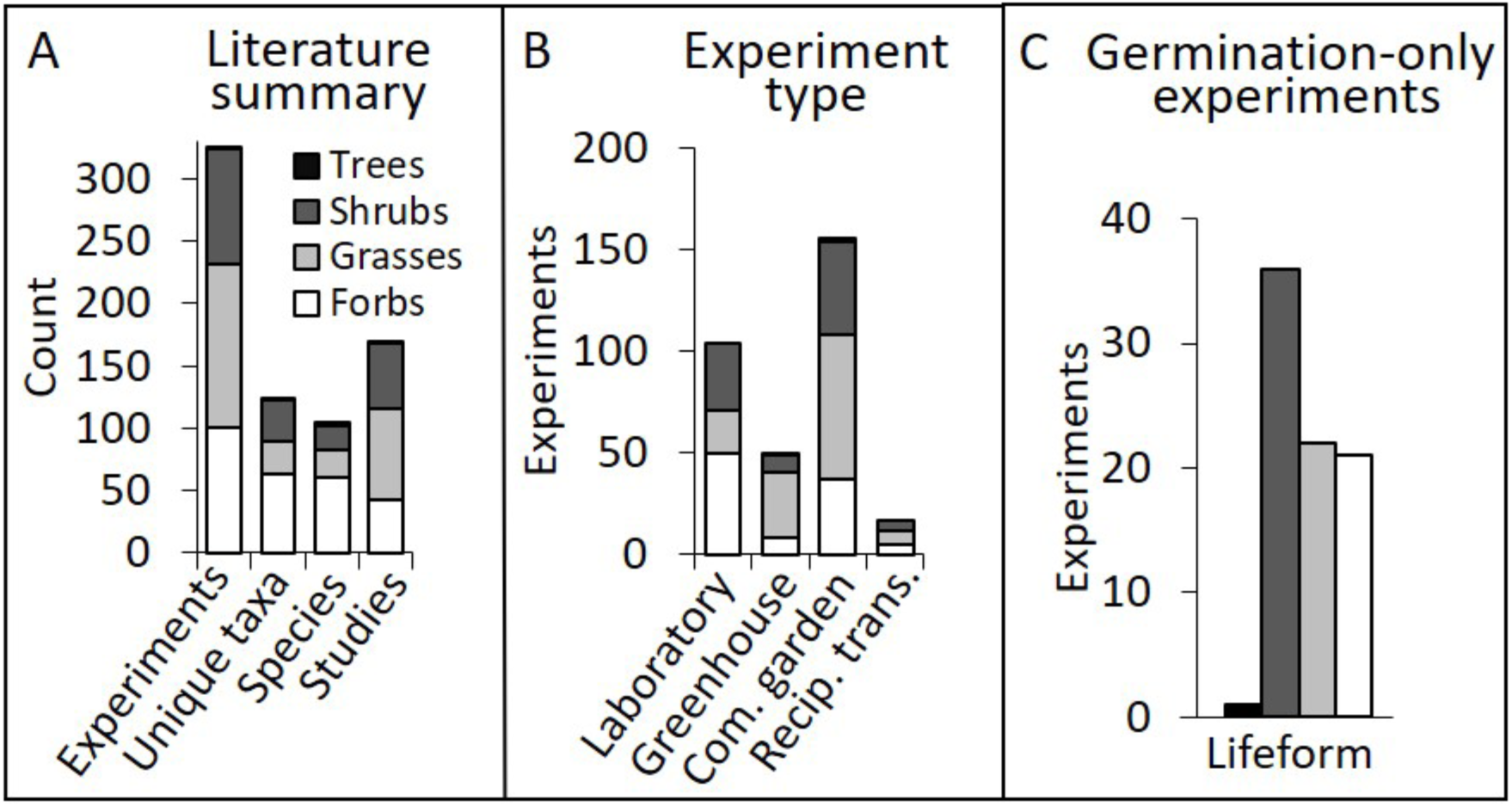
Summary of reviewed literature that compared traits among at least two populations in at least one common environment, by lifeform. Total counts of published studies, species, taxa, and taxa-specific experiments (A); types of experiments (B); means and standard errors of duration of the experiments that measured more than germination traits (C); total counts of experiments that measured only germination traits, (D); means and standard errors of number of populations tested in each experiment (E), and garden sites per experiment for outdoor reciprocal transplant and common garden experiments (F).

**Figure 2.**
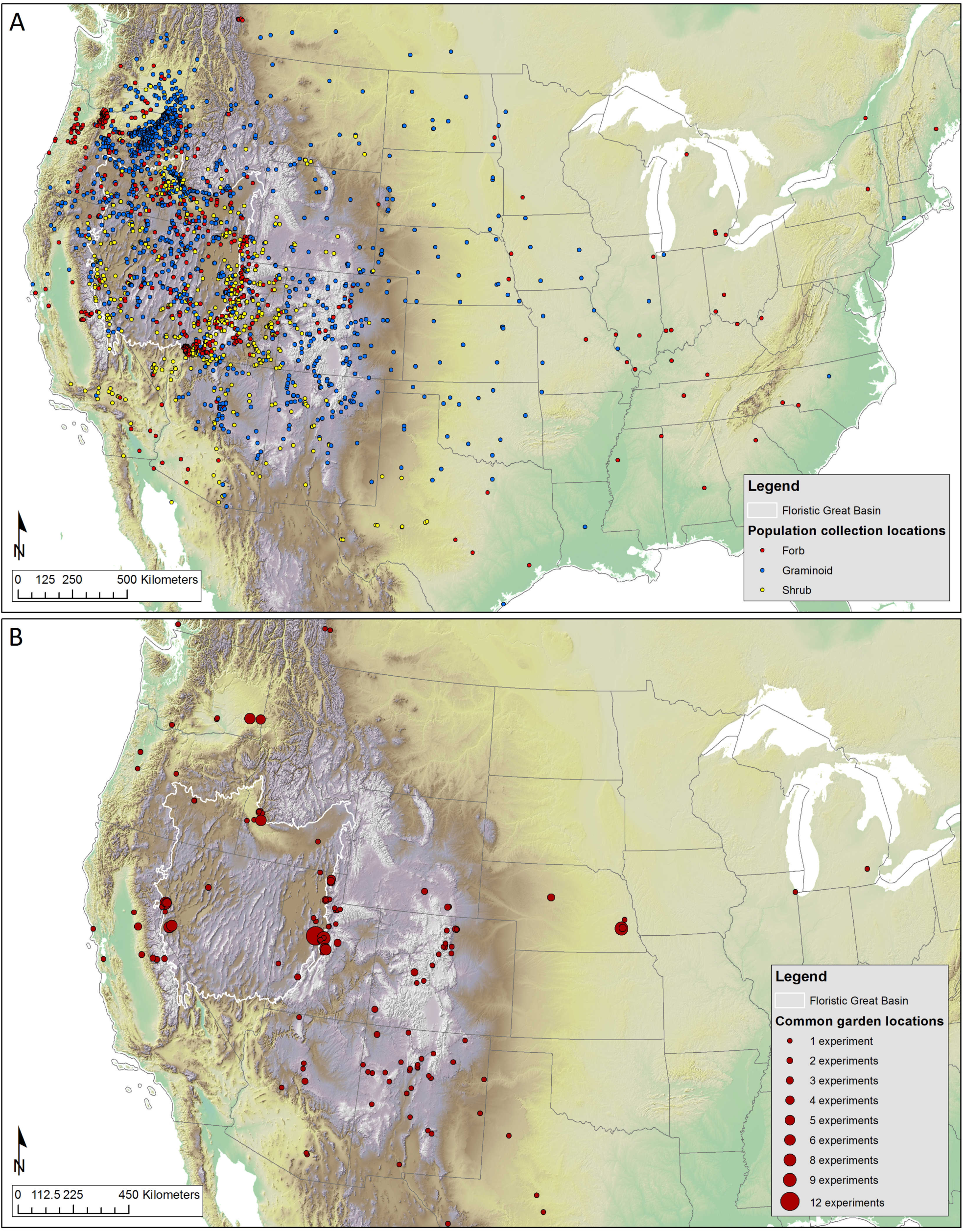
Map of 129 different outdoor common garden locations (A) and 2953 unique population collection sites (B) for the 80% of outdoor gardens and 91% of experiments for which coordinates could be obtained or generated, from 170 studies reviewed. The size of the marker in panel A represents the number of experiments in which each specific garden location was used, with larger symbols indicating garden locations used in more experiments. Although all species represented are native to the floristic Great Basin (white outline), many populations were collected and tested outside this region.

Experiments took place between 1940 and 2015, with collections from native stands occurring between 1938 and 2013 (Fig. 3A). One quarter of the experiments (24.5%) reported only early germination and seedling stages of plants (generally less than 0.5 years), while the remaining experiments (75.5%) reported study periods ranging from 0.5 to 17 years, with an average of 2.1 years (Fig. 3B, C). Average pairwise geographic distance among populations per experiment for the 91% of experiments for which coordinates were available was 351 km±20 SE, with a range from 610 m to 2,551 km. Most experiments were conducted on taxa with regional distributions, perennial species, grasses, and outcrossing species; very few annuals, endemic species, or selfing species were represented (Fig. 4). Over half of experiments (58.6%) tested plants grown directly from wild-collected seeds (or the seed of wild collected adults), 16.9% tested wild-collected adults, 13% tested materials with mixed generations since collection, 6.7% tested 1st or 2nd generation descendants of wild collected seeds, 0.3% tested only cultivars, and 4.3% did not provide enough information to determine.

**Figure 3.**
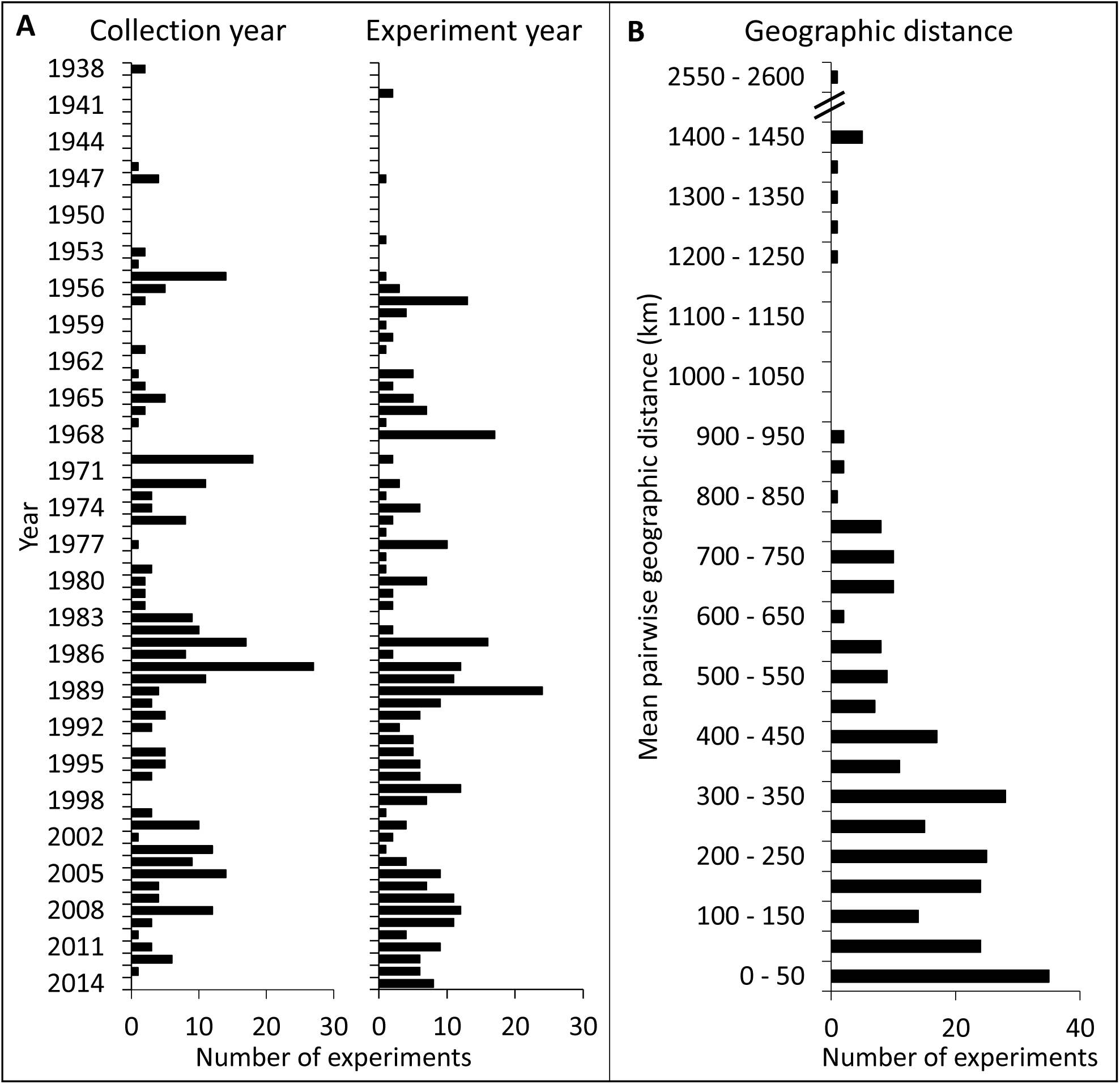
Summary of the years in which the collections of each experiment were made (A, left), the year each experiment was performed (A, right), and the average geographic distance among population collections sites in each experiment. The percent of 327 experiments that reported this information were 99% and 88% (respectively) for panel A, and 80% for panel B. Collection year and experiment year represent the average for each experiment, as it was common for materials to be collected and tested over multiple years for each experiment. Geographic distance is the mean pairwise distance among populations in each experiment; note the noncontinuous vertical axis.

**Figure 4.**
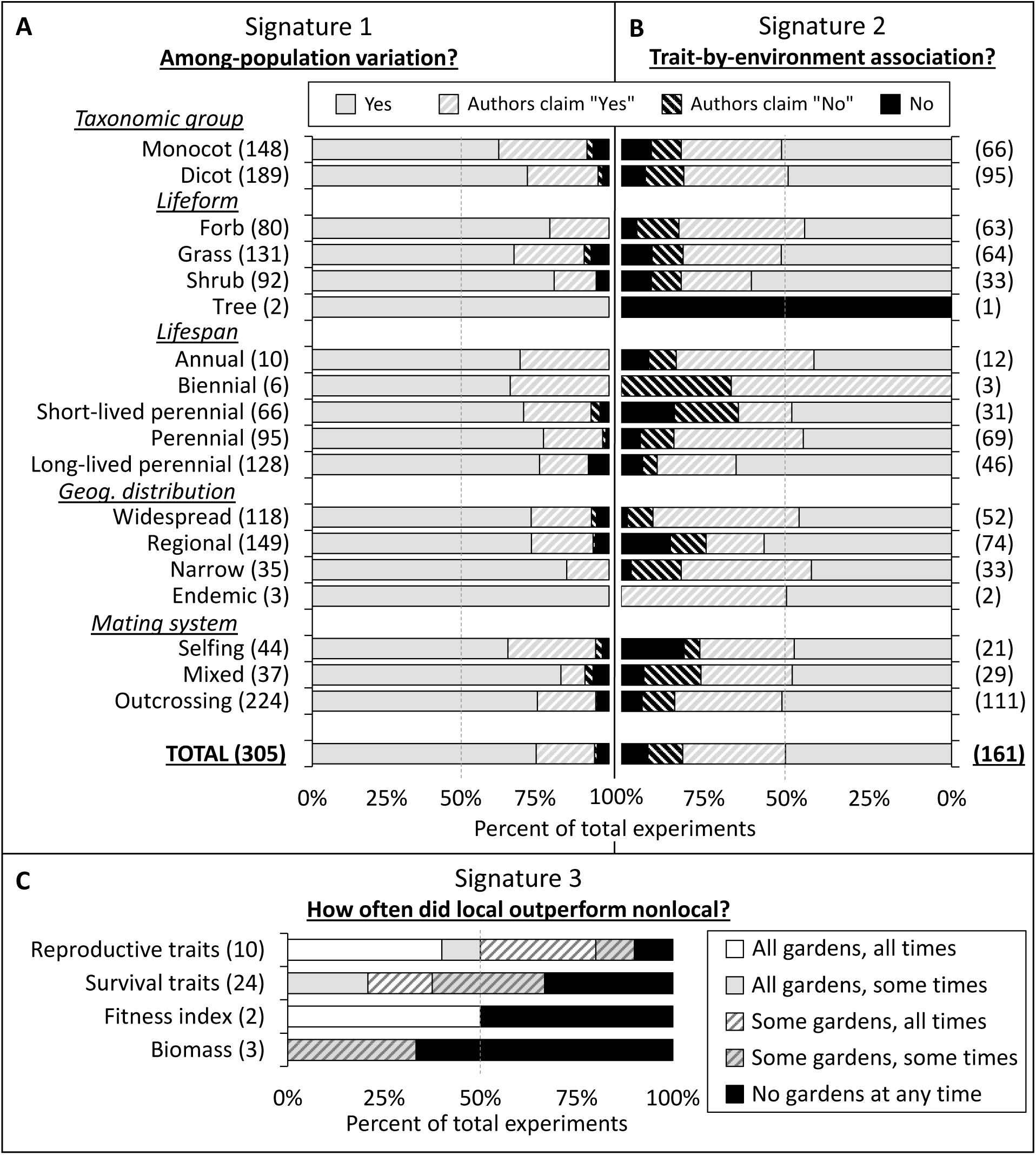
Summary of among-population variation (A, signature 1) and trait-by-environment associations (B, signature 2) for any measured trait, grouped by five life history traits. Summary of local advantage (C, signature 3) for reproductive traits, survival traits, fitness indices, or biomass. Data compiled from 327 experiments from 170 published studies on Great Basin plants (see Supporting Information Appendix 2 and available datasets in electronic supplementary material). For signatures 1 and 2, “Yes” and “No” represent statistical comparisons, while “Authors claim “Yes”” and “Authors claim “No”” represent textual, claim-based results where supporting statistics were not reported (common in older studies). For signature 3, most experiments had multiple gardens, and many evaluated performance at multiple sampling dates, leading to 5 different scores. These scores, from “All gardens, all times” to “No gardens at any time” represent a gradient of incidence and frequency of this signature (see methods). For all panels, numbers in parentheses, (x), indicate the number of experiments scored in a given category, and the dashed gray lines indicate 50%.

### Among-population variation

Of the 305 experiments appropriate for addressing among-population trait variation (signature 1), 290 (95.1%) experiments reported finding variation among populations in at least one phenotypic trait, with 230 (75.4%) of these 290 reporting significant variation, and 60 (19.6%) claiming such variation in the absence of any supporting statistics (Fig. 4A). Only 12 (3.9%) experiments reported no such differentiation in any trait after statistically testing for it, and 3 (1%) claimed no such variation without presenting statistical evidence. When categorized by basic life history traits, several differences appeared among groups. Eudicots exceeded monocots (the majority of which were grasses) in the degree of population differentiation(X_1_^2^ = 7, *P* = 0.0081), and, similarly, forbs and shrubs had more population differentiation than grasses (X_2_^2^ = 8.05, *P* = 0.0143). There were no significant differences in signature 1 among plants with different geographic distributions, life span, or breeding systems.

A total of 1,465 trait scores were recorded from the 305 experiments appropriate for addressing signature 1. Frequently-measured traits (20 or more experiments) that had differences between populations in over 75% of experiments (with or without supporting statistics) were floral structure, vigor, emergence, plant size, number of leaves, plant structure, shoot biomass, leaf structure, and number of inflorescences (Fig. 5).

**Figure 5.**
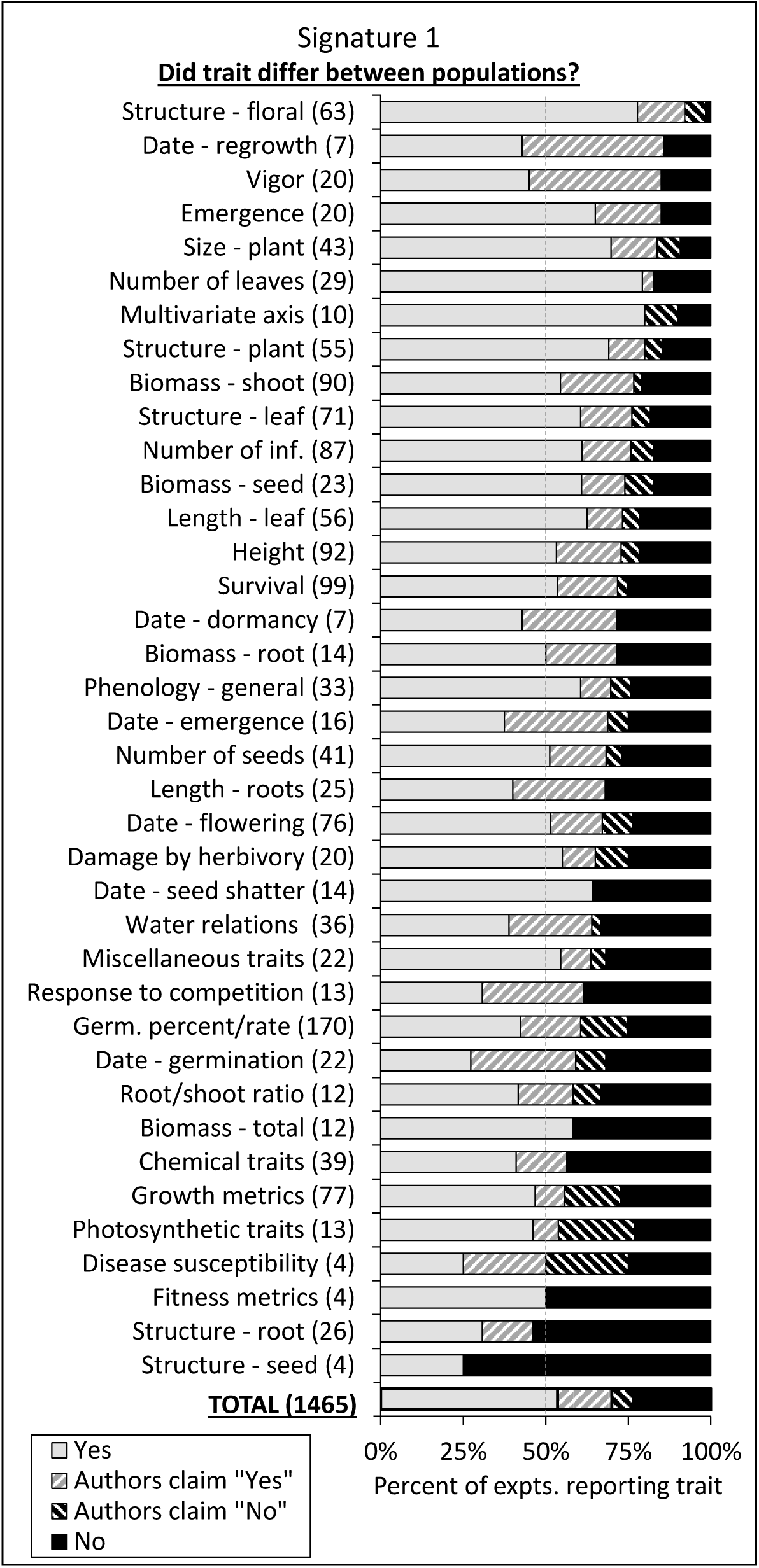
Summary of 1,465 trait scores from the 305 experiments appropriate for detecting signature 1 (differences between populations). Scores of “Yes” and “No” were supported by statistical comparisons, while the “Authors claim…” scores represent textual, claim-based results where supporting statistics were not reported (common in older studies). Numbers in parentheses, (x), indicate the total experiments that measured each trait or reported each factor, and dashed gray line indicates 50%.

### Trait-by-environment associations

Of the 161 experiments appropriate for testing trait-by-environment associations (signature 2), 131 (81.4%) reported associations for at least one comparison, with 81 (50.3%) supported by statistical tests and 50 (31.1%) supported by claims in the absence of statistics (Fig. 4B). Conversely, 13 (8.1%) of experiments reported no such correlations after having statistically tested for it, and 17 (10.6%) reported no such correlations but lacked any supporting statistics. There were no significant differences in the commonness of trait-by-environment associations for taxonomic status, lifeform, geographic distribution, or breeding system, but perennials (both long-lived and short-lived) had more frequent correlations between traits and environment than did annuals or short-lived perennials (X_3_^2^ = 8.08, *P* = 0.0444).

A total of 592 trait scores were recorded from the 161 experiments appropriate for addressing signature 2 (Fig. 6A). Frequently-measured traits (20 or more experiments) that were correlated with environmental variables in over 75% of experiments (with or without supporting statistics) were multivariate trait axes, floral structure, and germination date. Every remaining trait that was measured in >15 experiments was correlated with environmental characteristics in over 50% of experiments, and many, including leaf length, survival, flowering date, and leaf structure, were correlated with environmental variables in ≥70% of experiments.

**Figure 6.**
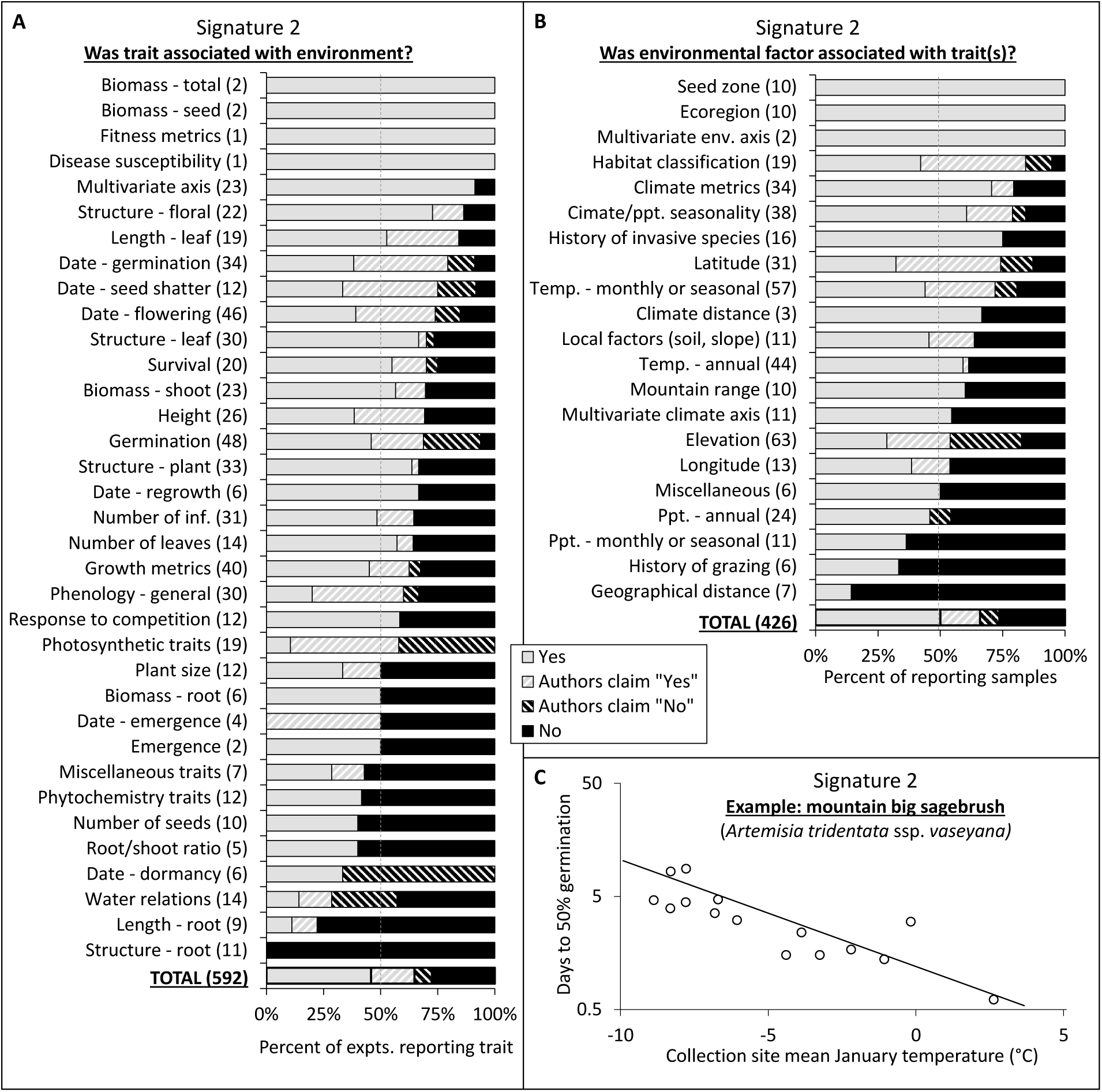
Summary of scores for associations between 592 traits (A) and 426 environmental factors (B) from the 161 experiments appropriate for detecting signature 2 (trait-by-environment association), expressed by trait/factors, and an example from the literature (C, redrawn with permission from (Meyer and Monsen, 1991)) in which date of germination for mountain big sagebrush is correlated with a measure of monthly temperature (treatment: 2-week chill). Scores of “Yes” and “No” were supported by statistical comparisons, while the “Authors claim…” scores represent textual, claim-based results where supporting statistics were not reported (common in older studies). For panels A and B, numbers in parentheses, (x), indicate the total experiments that measured each trait or reported each factor, and the dashed gray lines indicate 50%.

A total of 426 environmental variable scores were recorded from the 161 experiments appropriate for addressing signature 2 (Fig. 6B). Of the variables most frequently reported as correlated with plant traits, many categorical variables or composite metrics made this list, with seed zones, ecoregions, multivariate environmental axes, and habitat classifications topping the list of important environmental variables (important in > 84% of experiments that reported them). Additionally, derived climate metrics (such as climate continentality, heat/moisture index, potential evapotranspiration, etc.), climate seasonality, and history of invasive species presence were correlated with plant traits in over 75% of studies that reported them.

### Higher local performance in a local common garden

The 27 experiments that were suitable for detecting higher fitness of a local population in a local garden (signature 3) generated 39 scores (some experiments measured multiple fitness traits), with 27 scores (69.2%) reporting signature 3 for at least one fitness trait in at least one of the tested gardens during at least one sampling date, and the remaining 12 scores (30.8%) not reporting signature 3 at any point (Fig. 4C). Thirty-two of the 39 scores (82%) were generated from experiments with more than one garden. Survival was the most frequently measured fitness trait in these experiments, reported in 24 of the 27 experiments, followed by reproduction (10), biomass (3), and fitness indices (2). Incidence of the local-does-best pattern was highest in experiments that directly measured reproductive output, with 90% reporting higher values for locals at some point in an experiment, followed by survival (67%), fitness indices that incorporated biomass (50%), and biomass measures (33%). For experiments in which only “some” gardens showed local-does-best patterns (Fig. 4C, hashed bars), the percentage of gardens showing this trend was 40%, 50%, and 40% for reproduction, survival, and biomass traits, respectively (not shown). For experiments in which only “some” sampling dates showed local-does-best patterns (gray bars), the percentage of sampling dates showing this trend was 56%, 47%, and 25% for reproduction, survival, and biomass traits, respectively (not shown).

### Considering possible biases: highly-studied species and maternal effects

The number of experiments per species in our dataset ranged from 1 (52 species) to 25 (*Artemisia tridentata*), with a median of 1 (IQR = 1 – 4). The most highly-represented species were *Artemisia tridentata* (25 experiments), *Elymus elymoides* (24), *Ericameria nauseosa* (17), *Achnatherum hymenoides* (17), *Krascheninnikovia lanata* (13), *Pascopyrum smithii* (11), *Atriplex canescens* (9), *Leymus cinereus* (9), and *Poa secunda,* (8). Results in which scores were averaged for each species (see methods) were similar to uncorrected results: signature 1 was 4% higher when corrected (98% vs. 94%), signature 2 was 1% lower when corrected (79% vs. 80%), and signature 3 was 8% higher when corrected (78% vs. 70%). Thus, uncorrected calculations were used throughout our study.

Only 19 experiments (5.8%) used an experimental design that could control for maternal effects (e.g. growing all populations for a generation in a common environment before initiating an experiment). An additional 30 experiments (9.2%) were unclear on this point, and the remaining 278 (85%) experimented directly on populations differing in maternal environment. The incidence of population differences (signature 1) was 100% in the 16 experiments that moderated maternal effects, 95% for the 259 that did not make an attempt, and 97% for the 30 which were unclear. Too few of the experiments that attempted to control for maternal effects were appropriate for measuring signature 2 (4 experiments) and signature 3 (1 experiment) to compare incidences of these signatures.

### Quantitative comparison of trait-by-environment associations

Overall, we found positive relationships between the magnitude of differences among populations in all eight phenotypic traits and the magnitude of differences between MAT and MAP at the collection locations (Fig. 7). The strongest relationship was observed between differences in flowering time and differences in MAT, and leaf size also showed a strong relationship with MAT. Multiple strong relationships were observed between trait/environment divergence for MAP, with leaf size, survival, shoot mass, inflorescence number, and flowering time all showing strongly positive relationships for grasses, forbs, and shrubs. (Fig. 7, Supporting Information Appendix 3). Regression analyses demonstrated that, for the 15 common garden locations in which strong flowering time and MAT relationships were observed, each degree change in MAT was associated with a median change of 3.5 days (IQR = 1.2 - 5.3) in flowering time. Small sample sizes (few experiments that could be included in the analyses) and challenges with interpreting changes in physical traits across species of various shapes and sizes precluded the presentation of estimates of this nature for the other trait-by-environment relationships.

**Figure 7.**
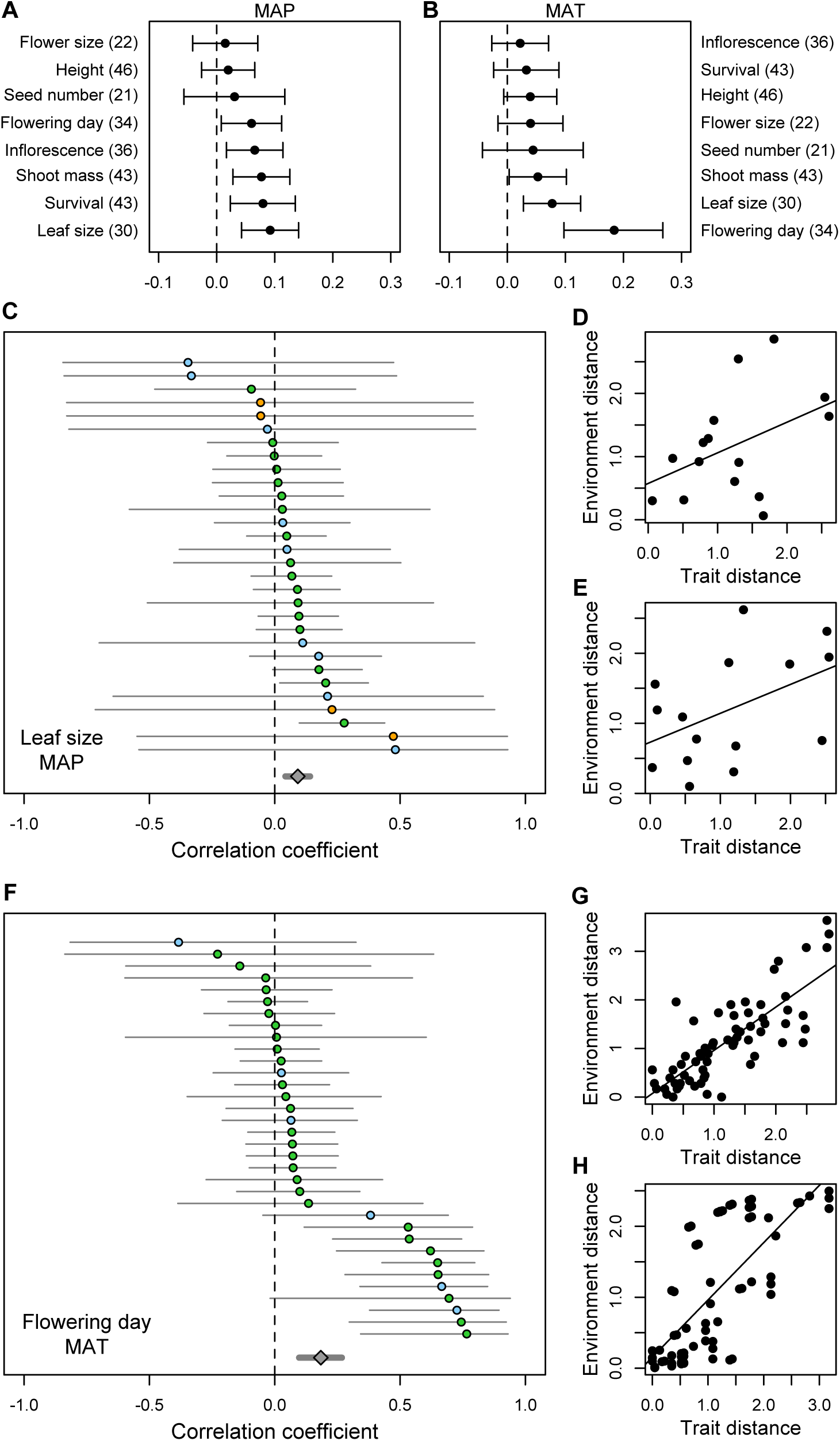
Results of comparisons of pairwise trait and environmental distances for eight frequently measured phenotypic traits and (A) the mean annual precipitation (MAP) or (B) mean annual temperature (MAT) at the original collection location. Values are effect sizes and 95% confidence intervals for each trait, averaged across all experiments for which data were available (number of experiments in parentheses). Examples of the two strongest relationships are shown for leaf size and MAP (C), where each line shows the correlation coefficient and confidence intervals for an individual experiment, for which we calculated the relationship between differences in percent survival and difference MAP at location of origin. Color indicates functional groups: Green = grasses, blue = shrubs, orange = forbs. Examples are shown for the two highest effect sizes: D), experiment 297A, (Kramer, Larkin and Fant, 2015), *Penstemon deustus* and E), experiment 297A, (Kramer, Larkin and Fant, 2015), *Eriogonum microthecum*. Similarly, flowering time and MAT (F) is shown, with examples of G) experiment 271A, (Larsen, 1947), *Schizachyrium scoparium*, and H) experiment 245A, (Ward, 1969), *Deschampsia caespitosa*. Full results for each trait/environment relationship are shown as additional results in Supporting Information Appendix 3.

## Discussion

Our results represent the most extensive review of intraspecific variation and local adaptation for plants native to the floristic Great Basin, a region comprised of largely continuous but increasingly imperiled arid and semi-arid plant communities (Davies *et al.*, 2011; Finch *et al.*, 2016). Additionally, they represent a significant addition to the noteworthy though relatively small number of reviews investigating this topic in a manner that identifies individual traits and environmental factors involved. We found that Great Basin plant species contain large amounts of intraspecific diversity in a wide range of phenotypic traits, that differences in these phenotypic traits are often associated with the heterogeneous environments of origin, and that differences among populations are commonly relevant to outplanting fitness. The cascading importance of intraspecific variation for the structure, functioning, and biodiversity of communities and ecosystems can be considerable (Bolnick *et al.*, 2011; Bucharova *et al.*, 2016), and may equal or exceed the importance of species diversity (Des Roches *et al.*, 2018). Our quantification of local adaptation and trait-environment associations should serve as encouragement to seriously consider intraspecific diversity in native plant materials used in restoration and conservation in this region throughout the selection, evaluation, and development process (Basey, Fant and Kramer, 2015). The results reported here should also serve as a cautionary note to restoration approaches that focus on only a few specific traits or search for general-purpose genotypes. Our results suggest that, in the absence of species-specific information to the contrary, it is reasonable to assume that local adaptation is present in this region, and that locally-sourced populations would outperform non-local populations a majority of the time.

Our investigation encompassed 170 studies published between 1941 and 2017 in which over 3,230 unique populations of 104 native Great Basin plant species were compared in 327 experiments, ranging from laboratory germination trials to multiple-year common gardens and reciprocal transplants. The great majority (95%) found differences between populations (signature 1) in the majority of traits measured in a common environment, which indicates that different traits are variable among populations, at both small and large geographic scales. Additionally, a clear majority (81.4%) of experiments found trait-by-collection environment associations (signature 2), suggesting that intraspecific variation is frequently an adaptive outcome of natural selection in heterogeneous environments (Linhart and Grant, 1996; Reich *et al.*, 2003). In experiments suitable for detecting local performance advantages (signature 3), local populations had higher performance (measured by differences in reproductive output, survival, and biomass) than nonlocal populations more often than not (69.2%), and this was particularly true when researchers reported traits related to reproductive output (90%). We used a vote-counting method to summarize results for our broadest pool of studies, allowing us to incorporate a wealth of older studies for which quantitative details were not available. Results from a vote-counting approach can sometimes differ from results of meta-analysis, as vote-counting does not incorporate the same level of detail about factors such as study size or effect size (Combs *et al.*, 2011). However, in our study, the overall incidence of “local does best” in the Great Basin is similar to other reviews that have found local adaptation to be commonplace, but not ubiquitous. In a review of local adaptation in plants that compared survival, reproduction, biomass and germination traits in reciprocal transplants, Leimu and Fischer (2008) found that local plants outperformed non-local ones in 71% of 35 published experiments. Similarly, Hereford (2009) quantified local adaptation in 70 published studies (50 of them plants), reporting only survival or reproductive traits, and found evidence of local adaptation in 65-71% of experiments. Our results indicated that the strongest indication of local adaptation came from experiments that directly measured reproductive output, and that using biomass as a fitness proxy may not be an effective way to compare relative performance in the Great Basin. This is consistent with a previous study that demonstrated selection for smaller, rather than larger, individuals in disturbed arid systems (Kulpa and Leger, 2013). Literature reviews conducted across biomes may occlude regionally-important trait differentiation and mask patterns of local adaptation, as we might expect, for example, biomass to be more strongly linked to fitness in regions where light is a contested resource (Espeland, Johnson and Horning, 2017).

There are many processes that can reduce or prevent the development of local adaptation, such as the lack of divergent selection between sites, high gene flow, rapid or extreme environmental change, high phenotypic plasticity, and/or low genetic diversity (Sultan and Spencer, 2002; Kawecki and Ebert, 2004; Blows and Hoffmann, 2005). The high incidence of intraspecific variation, much of it habitat-correlated, that we found in the literature confirms that divergent selection by heterogeneous environments is the norm for species native to the Great Basin, presumably outweighing the balancing effects of gene flow and genetic drift. Key environmental factors in the Great Basin such as fire frequency, grazing regimes, resource availability, and climate are certainly being altered to varying degrees by invasive species introductions, changing land uses, and climate change, and it can be argued that such changes could outpace the ability of local populations to remain adapted to their surroundings (Jones and Monaco, 2009; Breed *et al.*, 2013; Havens *et al.*, 2015; Kilkenny, 2015). However, our analysis also demonstrated relatively high instances of trait correlations with relatively recent disturbances such as invasive species introductions. Rapid evolution in response to invasive species (Oduor, 2013) and other anthropogenic changes (Hoffmann and Sgrò, 2011; Franks, Weber and Aitken, 2014) has been documented for many species, indicating that local adaptation can evolve rapidly in some circumstances.

Some traits and environmental characteristics stood out as particularly important indicators of local adaptation and its signatures across the studied taxa. For example, in our quantitative comparison of divergence in traits and environments, flowering phenology was strongly affected by MAT, with a median change of 3.5 days in flowering time per degree change in MAT of collection origin. Flowering phenology, along with germination phenology, were also in the top tier of frequently measured traits that showed significant correlations with environmental variables, consistent with other studies that have shown reproductive (Bucharova, Michalski, *et al.*, 2017) and germination (Donohue *et al.*, 2010) phenology to be an important response to environmental variation. Leaf size is also an important adaptive response to differences in temperature globally (Wright *et al.*, 2017), and in concert with this, we saw overall positive responses to MAP and MAT for leaf size in our analyses as well as frequent trait-by-environment associations in the literature. Floral structure, which has important adaptive significance for angiosperms (Harder and Barrett, 2007; Armbruster, 2014), was among the most frequent traits scored for among-population variation and trait-by-environment interactions. Seasonality of precipitation, which varies in this region depending on summer rainfall (Comstock and Ehleringer, 1992), was more predictive of trait variation overall than was mean annual precipitation (signature 2). In our quantitative comparisons, differences in MAP values were important for multiple phenotypic traits, including leaf size, shoot mass, reproductive output, and flowering phenology, in addition to being important for overall plant survival. Larger scale environmental descriptors, such as ecoregions and seed transfer zones, universally demonstrated signature 2, likely because they were developed based on climate/soil/vegetation associations or, in the case of seed transfer zones, developed based on trait-by-environment correlations. As found in other reviews (Geber and Griffen, 2003), physiological traits, phytochemical traits, and root traits were not measured as frequently as other traits, and though these did not show as frequent associations with environmental characteristics as other traits, they are known to vary across environments in some systems (Reich *et al.*, 2003). Additional studies of these traits in the Great Basin would be informative and could reveal different patterns than those observed here.

As in any review and analysis of published papers, there are elements of our design that were difficult to control. For example, consistent with other reviews (Gibson *et al.*, 2016), the vast majority of studies involved wild-collected plants or seeds, and thus maternal environment effects almost certainly affected some results (e.g. Bischoff and Müller-Schärer, 2010; Espeland *et al.*, 2016). Additionally, though the majority of populations tested in the literature were from western states, some of the populations compared in the literature were collected from well outside of the Great Basin, which increased the likelihood of observing local adaptation in these species. However, understanding patterns of intraspecific variation across the full range of the species native to the Great Basin is pertinent because it has been common (and for some species, ubiquitous) to utilize sources of native species originating from outside the Great Basin to use for restoration within the Great Basin (Jones and Larson, 2005). Finally, the scores and percentages for each of the signatures used throughout this study are uncorrected for phylogeny, as is our pairwise trait/environment analysis, and calculated such that each experiment is weighed equally. This introduces the possibility for phylogenetic biases, in which closely related taxa represented by many experiments affect the results more than less frequently studied taxa or groups of taxa. Though we did not conduct phylogenetic corrections for relatedness among taxa (Harvey and Pagel, 1991; de Bello *et al.*, 2015), our results were essentially identical for signatures 1-3 when we averaged results across species (scores differed by +3%, −1%, and +8%, respectively), suggesting that our lack of phylogenetic corrections are not unduly affecting our results. We present all species-specific information in Supporting Information Appendix 2 and available datasets section of the electronic supplementary material for further review.

Current approaches to seed sourcing in restoration and conservation include genetic (e.g. Williams, Nevill and Krauss, 2014), genecological (e.g. Johnson, Leger and Vance-Borland, 2017), local-only (e.g. Erickson *et al.*, 2017), predictive (e.g. Prober *et al.*, 2015), and agronomic (e.g. United States. House of Representatives. Committee on Appropriations., 2014)) strategies, as well as strategies mixing several of these viewpoints (i.e. Rice and Emery, 2003; Rogers and Montalvo, 2004; Breed *et al.*, 2013; Havens *et al.*, 2015; Bucharova *et al.*, 2018). These approaches vary in the degree to which they meet the needs of seed producers and land managers while balancing population differences that stem from adaptive evolution in different environments. The prevalence of local adaptation and its signatures found in our study justify and support incorporating existing best-practices (e.g. Basey, Fant and Kramer, 2015; Espeland *et al.*, 2017) for capturing and preserving important intraspecific variation into seed sourcing and plant production systems. For example, our results demonstrated a strong relationship between flowering time and MAT, so it would be wise to collect materials for research, evaluation, and testing from populations that vary in MAT, to collect seeds at multiple times to fully capture population variation in flowering time, and ensure that seeds are not transferred during restoration among sites that differ strongly in these characteristics. On the production side, best practices for seed harvesting should include methods that avoid inadvertent selection on flowering time, either for reduced variation or for a directional shift away from the wild condition. Similarly, emergence date was correlated with environmental variation in many plants, so testing in common gardens should involve seeding trials in place of or in addition to using transplants, and evaluation trials should guard against inadvertent selection on emergence timing by randomly, rather than systematically, selecting individuals to use in transplant experiments. These examples are not exhaustive, but demonstrate how evidence revealed by this study regarding which traits and environmental factors are generally involved in adaptation in this region can be used to improve approaches to seed sourcing and restoration. Finally, we acknowledge that ours is not the first review and meta-analysis to affirm an abundance of intraspecific variation and local adaptation in plants. However, our focus on the Great Basin is important, because the large and frequent yet commonly unsuccessful restoration efforts occurring in this region have lagged behind those of other regions with respect to recognizing the importance of intraspecific variation and local adaptation on outplanting success.

## Conclusions

Reestablishing and maintaining native plant communities in arid regions has proven challenging (Svejcar *et al.*, 2017), and the lack of practical knowledge guiding more appropriate selection of seed sources is a major barrier (Friggens *et al.*, 2012; Gibson *et al.*, 2016). The forestry industry has long adopted the principles of local adaptation in their reforesting guidelines with great success (Matyas, 1996; Johnson *et al.*, 2004; Aitken and Bemmels, 2016), and similar approaches to restoration in the rangelands of the Great Basin may also increase success as our data support similarly high levels of population differentiation within grass, forb and shrub life history groups. Our results, including both a qualitative literature survey and a quantitative meta-analysis, could benefit from future work using additional techniques to explore spatial structure (e.g. Griffith and Peres-Neto, 2006) and the relative importance of geographic distance and environmental variation, especially as additional studies become available in the literature. Nevertheless, our results as they currently stand are in agreement with observations of abundant local adaptation in plant populations world-wide, and further, we identified particular phenotypic traits (flowering and germination phenology, floral structures, leaf size, biomass, survival, and reproductive output), environmental characteristics (MAT, MAP, climate metrics, seasonality), and habitat classifications and site history (seed zones, ecoregions, history of invasive species) that were important predictors of local adaptation in plants native to the Great Basin floristic region. Given the speed and severity with which natural communities are being altered by anthropogenic factors, the application of an evolutionary perspective to restoration ecology is more important than ever. Adjusting seed-selection priorities to account for the existence of locally adapted, intraspecific variation in the Great Basin will promote the maintenance and recovery of resilient, self-sustaining vegetation communities in this region (Meyer, 1997; Lesica and Allendorf, 1999; Rogers and Montalvo, 2004; Broadhurst *et al.*, 2008; Vander Mijnsbrugge, Bischoff and Smith, 2010).

## Acknowledgements

We would like to acknowledge Vicki Thill and Sage Ellis for many hours of extracting coordinates and trait data, as well as Susan Meyer, Andrea Kramer, Tom Jones, Vicky Erickson, Allan Stevens, Dan Atwater, Clinton Shock, Jessica Irwin, Huixuan Liao, and David Solance Smith for providing information and data not available in their publications. We also thank several anonymous reviewers for their insightful suggestions that greatly improved this work.

## Dedication

We would like to dedicate this paper to the memory of our co-author Dr. Erin K. Espeland, friend and collaborator to all of us, who worked on this manuscript. Erin’s light and life will never be forgotten by those who knew her, and we want to recognize her creative contributions to the field of plant ecology, including this effort. Erin is dearly missed.

## Data accessibility

Raw datasets and statistical code supporting this study (Baughman *et al.*, In Review) have been deposited at Dryad, [DOI: TBD]

## Authors’ contributions

EAL, OWB, FFK, EKE, RF, TNK, and JBS conceived and designed the study; OWB conducted the literature search; OWB, ACA, FFK, JEO, RCJ, and JBS categorized, compiled and extracted data; OWB, EAL, FFK, ACA and MLF analyzed data; OWB, EAL, and ACA drafted the manuscript; all authors critically revised the manuscript for important intellectual content and approved of the version to be published.

## Appendix 1. Additional Methods

### Literature search

Terms used to search the literature included ‘plant’, ‘Great Basin’, ‘Intermountain West’, ‘western United States’, ‘local adaptation’, ‘ecotypic variation’, ‘phenotypic variation’, ‘genetic variation’, ‘habitat-correlated variation’, ‘genecology’, ‘intraspecific variation’, ‘ecotype’, ‘seed zones’, ‘common garden’, ‘reciprocal garden’, and ‘transplant garden’, as well as combinations of these terms. Literature was obtained primarily using the World Wide Web as well as databases such as Google Scholar, Web of Science, Academic Search Premier, JSTOR, Science Direct, and Wiley Online Library. When digital copies were not available, they were obtained from academic libraries. The citations within the resulting literature were also mined for additional literature that our first search had missed.

### Geographic range categorization

Four categories of geographic range were assigned from distributions in the USDA Plants Database (https://plants.sc.egov.usda.gov), as follows. Widespread: found in majority of United States (e.g. *Elymus elymoides* (Raf.) Swezey); Regional: common in the floristic Great Basin but not found throughout the United States (e.g. *Atriplex confertifolia* (Torr. & Frém.) S. Watson); Narrow: limited to specific, well-defined habitats within the Great Basin (e.g. *Penstemon confusus* M.E. Jones); Endemic: restricted to several counties (e.g. *Allium passeyi* N.H. Holmgren & A.H. Holmgren).

### Geographic coordinate generation

Geographic coordinates and elevations for gardens and populations were recorded verbatim from studies that contained precise coordinates, or were generated manually using Google Earth Pro (Google Inc., 2018) with assistance from the Geographic Names Information System (US Geological Survey, 2018) when vague coordinates or textual localities were given. All coordinates were converted to decimal degrees (WGS 84) and elevations were recorded in meters. Uncertainties in manually generated coordinates were recorded in a measure of accuracy, either ‘high’ (confident to within a ∼2 mile radius), ‘fair’ (confident to within a ∼5 mile radius), or ‘low’ (confident to within a ∼15 mile radius). Numeric coordinates given in the studies were assumed to be accurate to within one mile. If elevations were given for populations or gardens with vague localities, we utilized this information to increase the confidence of our generated location. Coordinates were not generated for localities that were exceptionally vague or studies which did not include localities. If a study utilized a named release or cultivar, the location of origin was determined by locating the original published release notice, if available. Cultivars bred using populations from multiple locations were not assigned origin coordinates.

### Scoring experiments for each signature of local adaptation

For among-population variation (signature 1), a score of ‘Yes’ was given when at least one measured trait was reported to differ significantly between at least two populations, and a score of ‘No’ was given when differences in any phenotypic trait were not detected between any pair of populations. For trait-by-environment association (signature 2), a score of ‘Yes’ was given when authors reported a significant association between at least one trait and one measure of the environment of origin, and a score of ‘No’ was given when the author tested for but found no such relationship. In addition to a score for each experiment, each of the measured and reported traits and environmental variables were scored (hereafter, trait scores) in a manner that indicated which traits did or did not vary between populations, as well as which traits and environmental variables were or were not correlated with each other (see available datasets in electronic supplementary material). Some experiments met the criteria for both signatures while others met only one or the other. In several studies, especially older studies or studies whose analyses did not include among-population comparisons, the significance of variation and/or correlation needed for scoring signatures 1 and 2 could not be determined because the authors provided results without statistical analyses. In these cases, results were scored as ‘Authors Claim Yes’ or ‘Authors Claim No’, and the scoring was done as described above, taking authors at their word in the absence of published statistical evidence.

To score whether there was higher fitness of a local population in a common garden (signature 3), only experiments in which outdoor reciprocal transplants or common gardens were performed using a local population (identified as such by the author, or clearly collected from the common garden site) in at least one garden were considered. Additionally, the experiment had to measure survival, reproductive output (number of seeds or flowers, or other reproductive output), a fitness index (a combination of several size and production traits), or total aboveground biomass. Each experiment was given a composite score to fully capture variation in the performance of the local population across gardens (spatial), as well as through different sampling dates (temporal). For the spatial component, ‘Yes for all gardens’ indicates the highest values in each garden belonged to that garden’s local population, ‘Yes for some gardens’ indicates the highest value in at least one but not all of the gardens belonged to each garden’s local population, and ‘No for all gardens’ if the highest value never belonged to a garden’s local population. For the temporal component, the experiment was scored as ‘Always’ if the local population had the highest value at all sampling dates, or ‘Sometimes’ if the local population had the highest value at one but not all of the sampling dates. For “some” and “sometimes” scores, we calculated the number of observations of higher fitness of local populations per garden and per time measured to understand what proportion of gardens and sampling dates showed higher local fitness. This provides an estimate of the frequency of higher local fitness, but it is not a measure of the importance of the difference per se. For example, a fitness difference could occur at a low frequency, but have a large impact on population trajectories (i.e. large differences in survival after a rare drought event).

### Determining whether maternal effects were controlled

Experiments which tested populations that had all shared one or more generations in the same location prior to testing were considered to have attempted to control for maternal effects. We determined the number of generation in common by carefully reading the methods for mentions of the populations’ lineages prior to testing. Some experiments supplied the original location of material collection but indicated that all materials were collected from areas such as ‘evaluation plots’, ‘seed fields’, ‘uniform gardens’, or ‘increase fields’, indicating that at least one generation was shared among all populations, and therefore and attempt had been made to control maternal effects (intentional or not). Some complex studied had to be split into multiple experiments because they used different generations of the same populations in different tests. For example, a study which collected wild adults from their native habitats and grew them in a common garden for the duration of the experiment before measuring traits of the plants as well as traits of the seeds they produced were split into two experiments, one containing the traits of the adult plants (which did not attempt to control for maternal effects, because the progenitors of the measured material did not share a common location), and one for the seed traits (which did attempt to control for maternal effects, because the progenitors of the measured material did share a common location).

### Extraction for quantitative comparison of trait-by-environment association

To examine links between the variation in trait values and the variation in environmental and geographic distance among the population’s origins, we utilized experiments from which population-specific trait data as well as geographic coordinates for at least one garden and at least two populations could be extracted or obtained through author contact. Data from laboratory and greenhouse experiments were not considered for this extraction, because the great majority of these experiments were not designed to completely simulate natural growing conditions. Excluding these experiments reduced our pool from 325 to 161. Next, a list of priority fitness traits were developed (Table S1-1) based on traits that were most commonly measured and potentially associated with plant fitness in the Great Basin (Bower, Clair, and Erickson, 2014; Leger and Baughman, 2015). Any experiment that did not measure at least one priority trait was omitted from next steps, and this further reduced our pool from 161 to 153.

**Table S1-1.** Priority traits targeted in the extraction for the dataset used in the quantitative comparison, and the preferred units. Note that for several traits, several highly similar measures were included, as indicated in footnotes.

| Trait | units | Trait | units |
| --- | --- | --- | --- |
| survival | % | number - inflorescence<sup>4</sup> | # |
| emergence | % | number – seeds<sup>5</sup> | # |
| germination | % | number – leaves | # |
| height - plant | cm | date – germination | # days |
| length – root | cm | date – regrowth/greenup | # days |
| length – leaf<sup>1</sup> | cm | date – emergence | # days |
| dimensions – floral<sup>2</sup> | cm | date – flowering<sup>6</sup> | # days |
| mass – roots | g | date – seed shatter | # days |
| mass – shoots<sup>3</sup> | g | date – senescence | # days |
| mass – seed | g/seed |  |  |

<sup>1</sup>If unmeasured, then leaf width was recorded, if available
<sup>2</sup>Any measure of a floral structure
<sup>3</sup>Any measure of aboveground biomass
<sup>4</sup>Any kind of count of flowers or flowering structures was recorded
<sup>5</sup>If no direct count was available, any measure of seed yield was recorded, including total seed yield in weight and/or seed yield rating/rank
<sup>6</sup>If unmeasured, any other floral phenology was considered

The remaining studies were then examined for textual, tabular, or visual data that could be extracted as mean values of priority traits for each population in each garden. Extracted values for were recorded verbatim from tables and throughout the text where possible, and from figures using WebPlotDigitizer (Rohatgi, 2017) when needed. Means for at least two populations in at least one garden were required for extraction. If exact matches to certain priority traits were not reported in the studies, similar measures that were likely to be strongly correlated to the given trait could be recorded as surrogates if available, and a note was made (footnotes, Table S1-1). We extracted the latest date for which the most populations at the most gardens were represented if studies presented data for multiple dates throughout the experiment. In some cases, experiments were conducted with multiple treatments in which growing conditions were altered to address study questions. In these cases, we only extracted data for the author-defined ‘control’ treatment. However, if no control was defined, we used the treatment that was the most unaltered or representative of the garden environment (e.g. unweeded, or unwatered).

## Appendix 2. Summary of literature and available datasets

The data collected and generated by this study (Baughman *et al.*, 2019), as well as the list of publications that were involved in each part of this study, are provided so that additional questions may be addressed and for other applications. We encourage such additional analyses.

### Summary of literature

**Appendix 2 Table 1.**
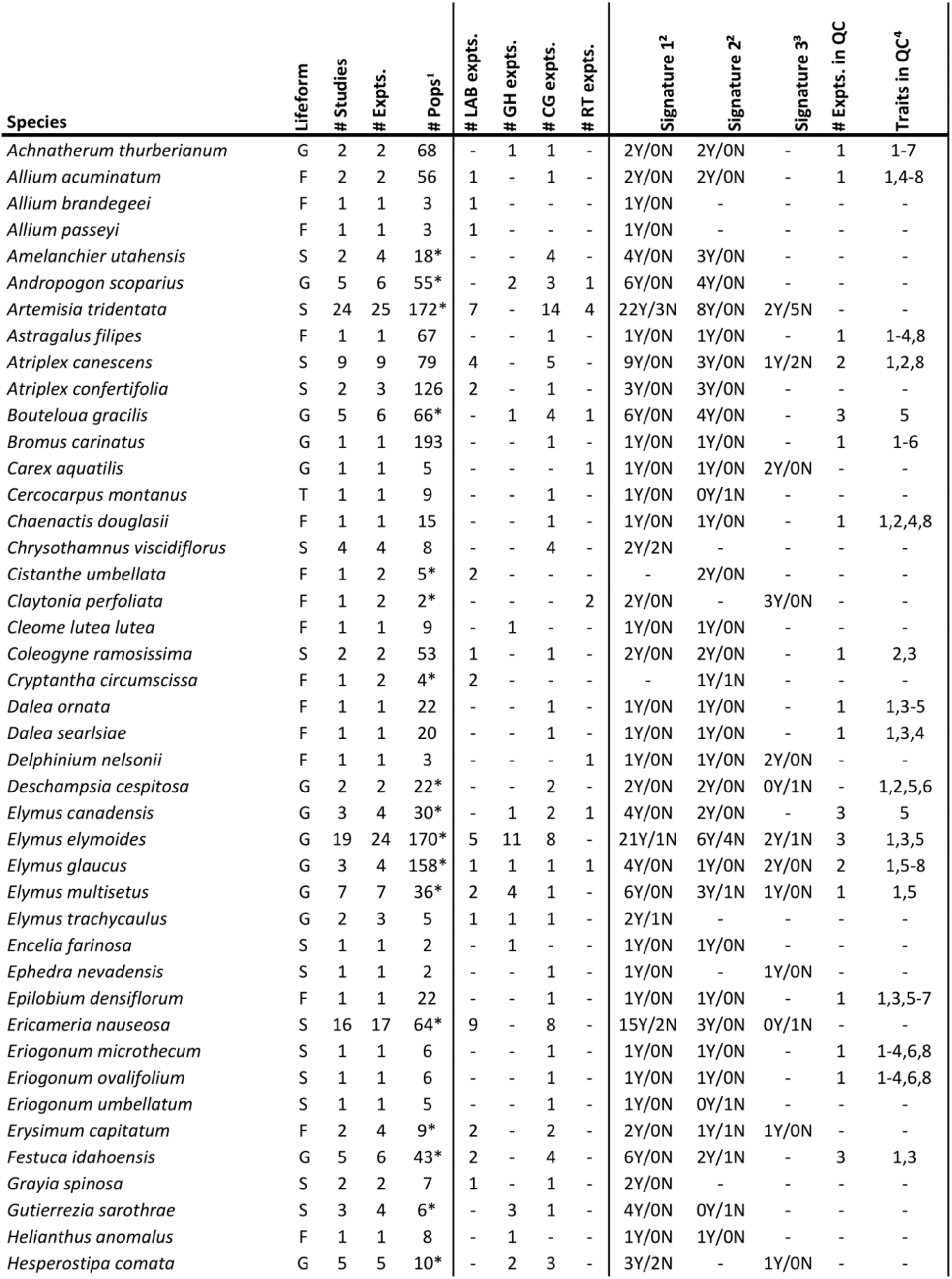
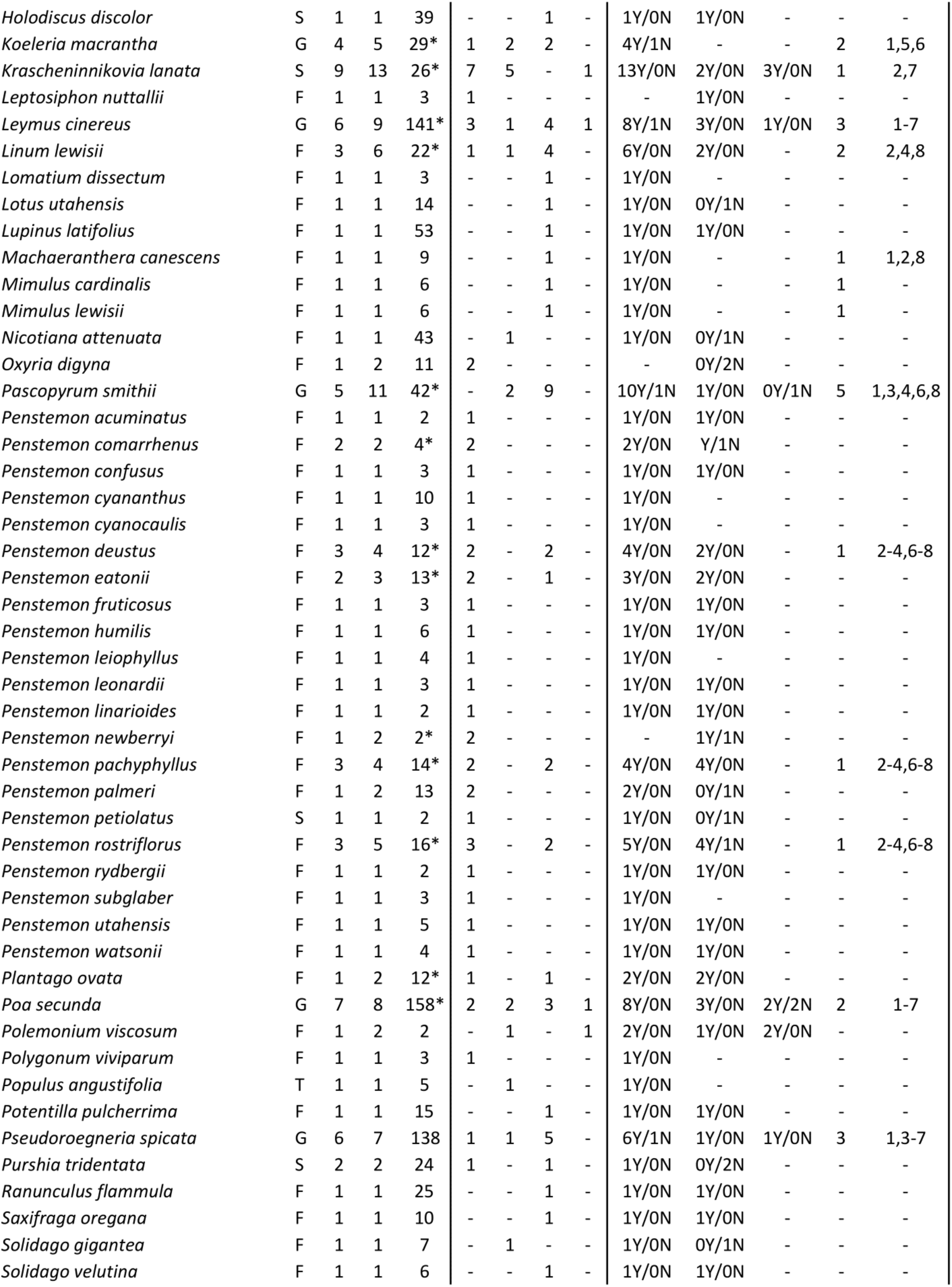
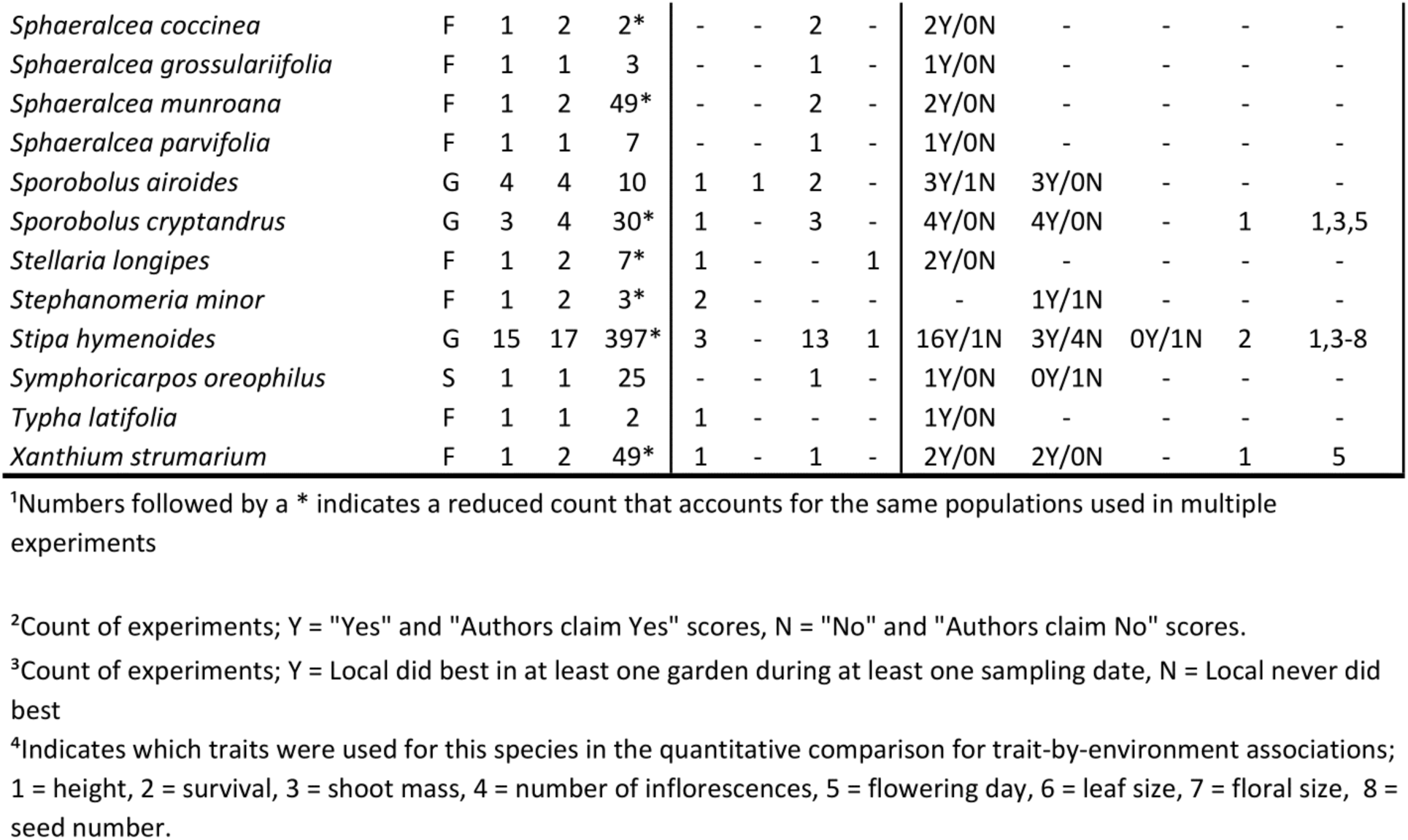
Summary, by species, of the literature included in this study, including lifeform (F = forb, G = grass, S = shrub, T = tree), counts of studies, experiments, unique populations, and experiments by type (LAB = laboratory, GH = greenhouse, CG = outdoor common garden, RT = outdoor reciprocal transplant), the incidence of each signature of local adaptation (1 = differences among populations, 2 = trait/environment correlations, 3 = higher performance of local than nonlocal population in local’s environment), counts of experiments used in the quantitative comparison of trait-by-environment associations (QC), and a list of traits used in the QC. See footnotes for additional information.

### Available datasets

Data have been uploaded to Dryad at DOI: TBD (Baughman *et al.*, In Review). Several datasets are available. The “Summary and signature scores” dataset includes all of the studies and experiments and summarizes literature categorization as well as scores and associated information for each of the signatures of local adaptation. The “Trait scores” dataset includes basic study categorization as well as information that indicated which phenotypic traits (for signatures 1 and 2) and environmental variables (for signature 2) were involved in each of the signatures of local adaptation. The “Quantitative comparison” dataset includes all of the data used to conduct the quantitative comparison of trait-by-environment associations, and lists population-specific mean values for our priority traits for all studies for which such data were available, the latitude and longitude of each population origin, and extensive climate information for each origin generated with the ClimateNA v5.10 software package based on methodology described by Wang et al. (2016). The “Location data” dataset lists all outdoor gardens and population origin coordinates and elevations for which authors gave this information, as well as those for which we could confidently generate it. For descriptions of each column in each of these datasets, refer to the “Data Dictionary” file.

### Bibliography of reviewed literature

A list of all the literature used in any of the datasets is provided below. Following each citation is a set of codes in brackets indicating which parts of our study the publication was used in. Codes S1, S2, and S3 indicate that at least one of the “experiments” in the given publication was used to generate a score for signatures 1, 2, and 3, and code QC indicates the publication (or the data summarized in it, even if not available from the publication itself) was used in analyses for the quantitative comparison of trait-by-environment associations. Note that some published studies were scored as multiple experiments for multiple species.

## Appendix 3. Additional Results

### Additional results of literature summary

Dicots accounted for 23.6% of the taxa and 42.8% of the experiments in the final pool of reviewed literature. Regional taxa accounted for 47.2% of the taxa and 48.9% of experiments, widespread taxa accounted for 26.0% of taxa and 36.7% of experiments, narrow taxa accounted for 24.4% of taxa and 13.5% of experiments, and endemic taxa accounted for 2.4% of taxa and 0.9% of experiments. Perennials accounted for 46.3% of taxa and 32.1% of experiments, long-lived perennials accounted for 32.5% of taxa and 39.4% of experiments, short-lived perennials accounted for 14.6% of taxa and 25.1% of experiments, annuals accounted for 5.7% of taxa and 3.1% of experiments, and biennials accounted for 0.8% of taxa and 0.3% of experiments. Primarily outcrossing plants accounted for 71.4% of taxa and 72.2% of experiments, primarily selfing plants accounted for 11.1% of taxa and 14.1% of experiments, and plants with mixed mating accounted for 17.5% of taxa and 13.8% of experiments.

### Additional results for quantitative comparison of trait-by-environment associations

**Appendix 3 Figures 1-16.**
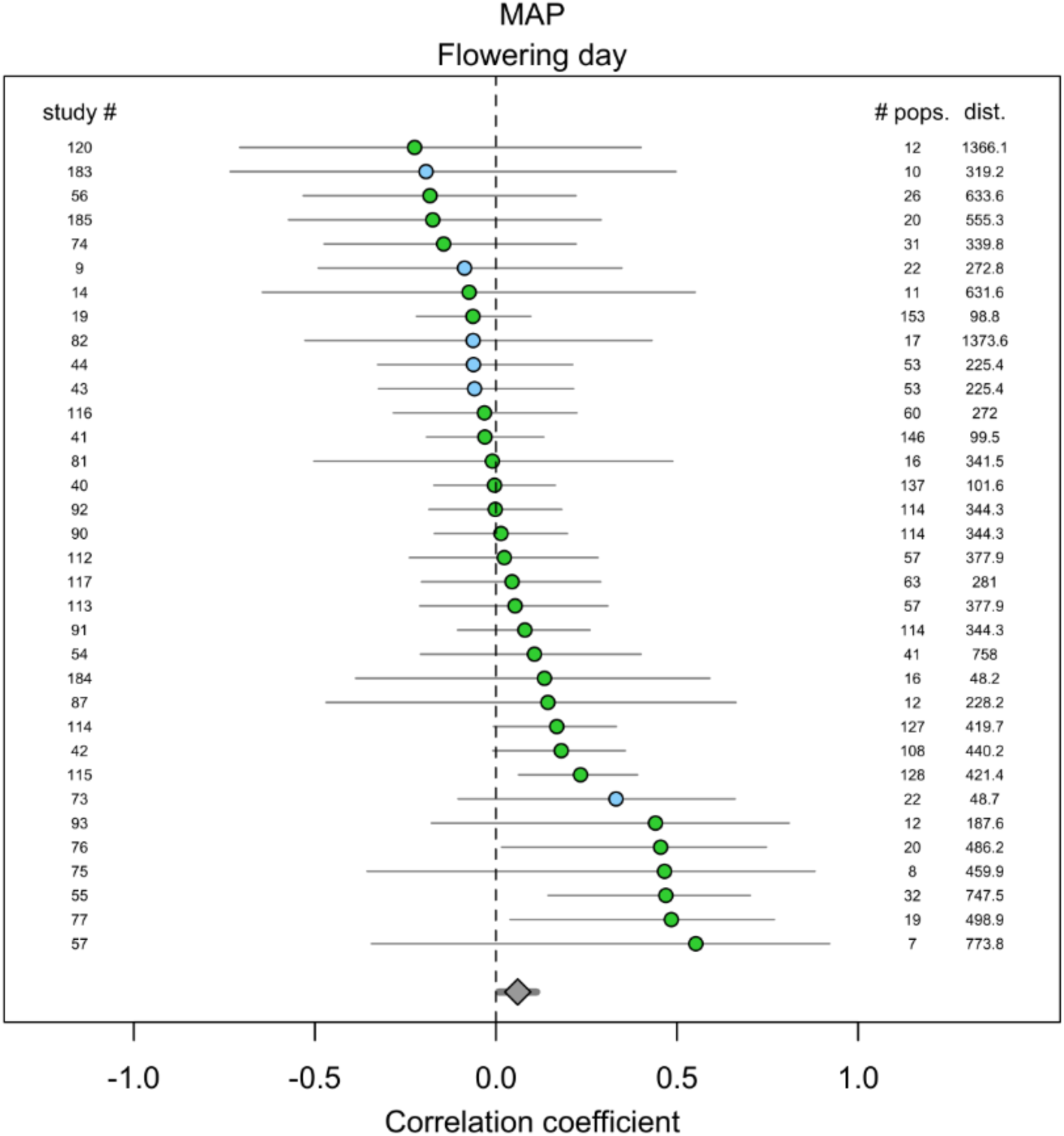
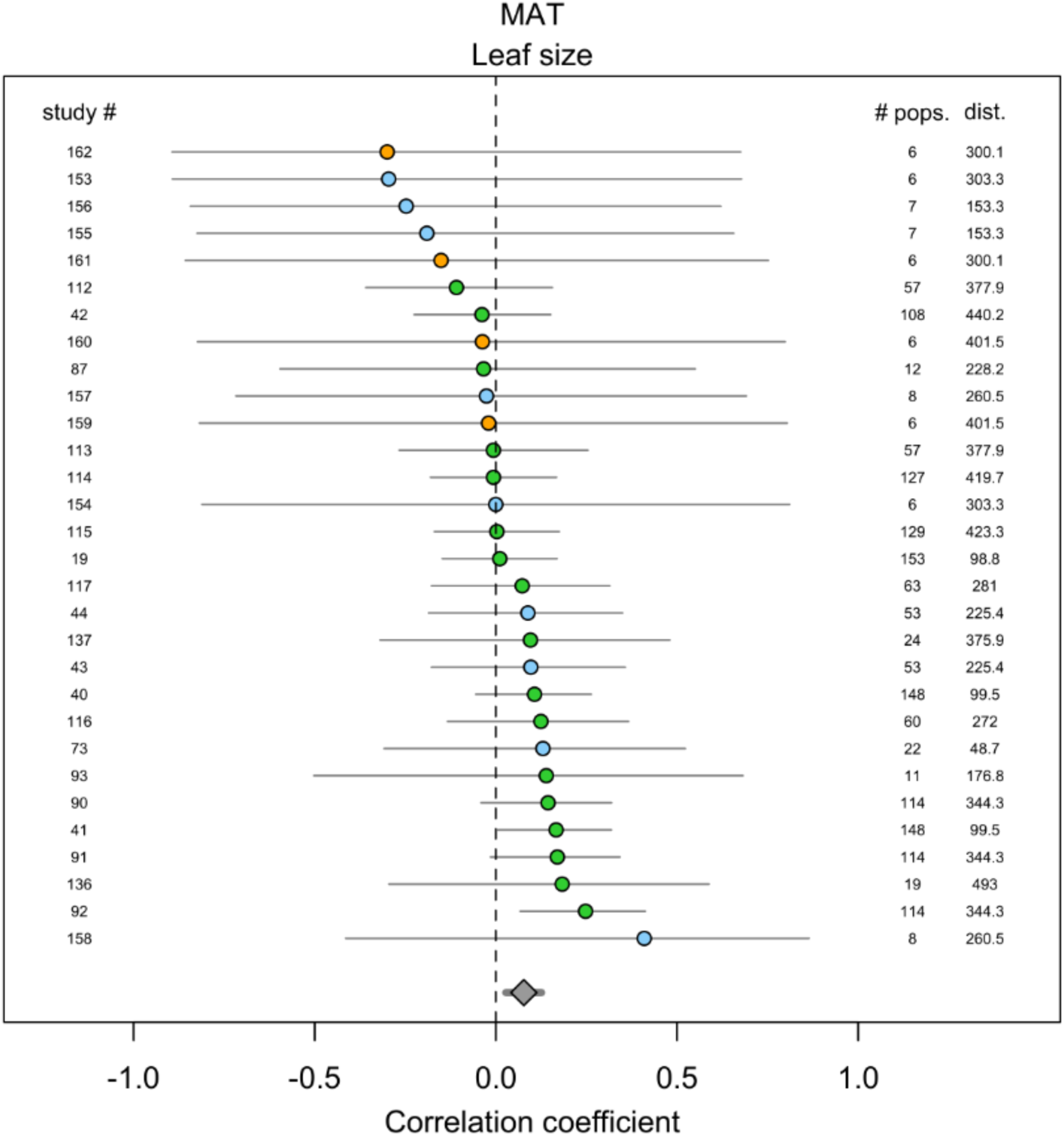
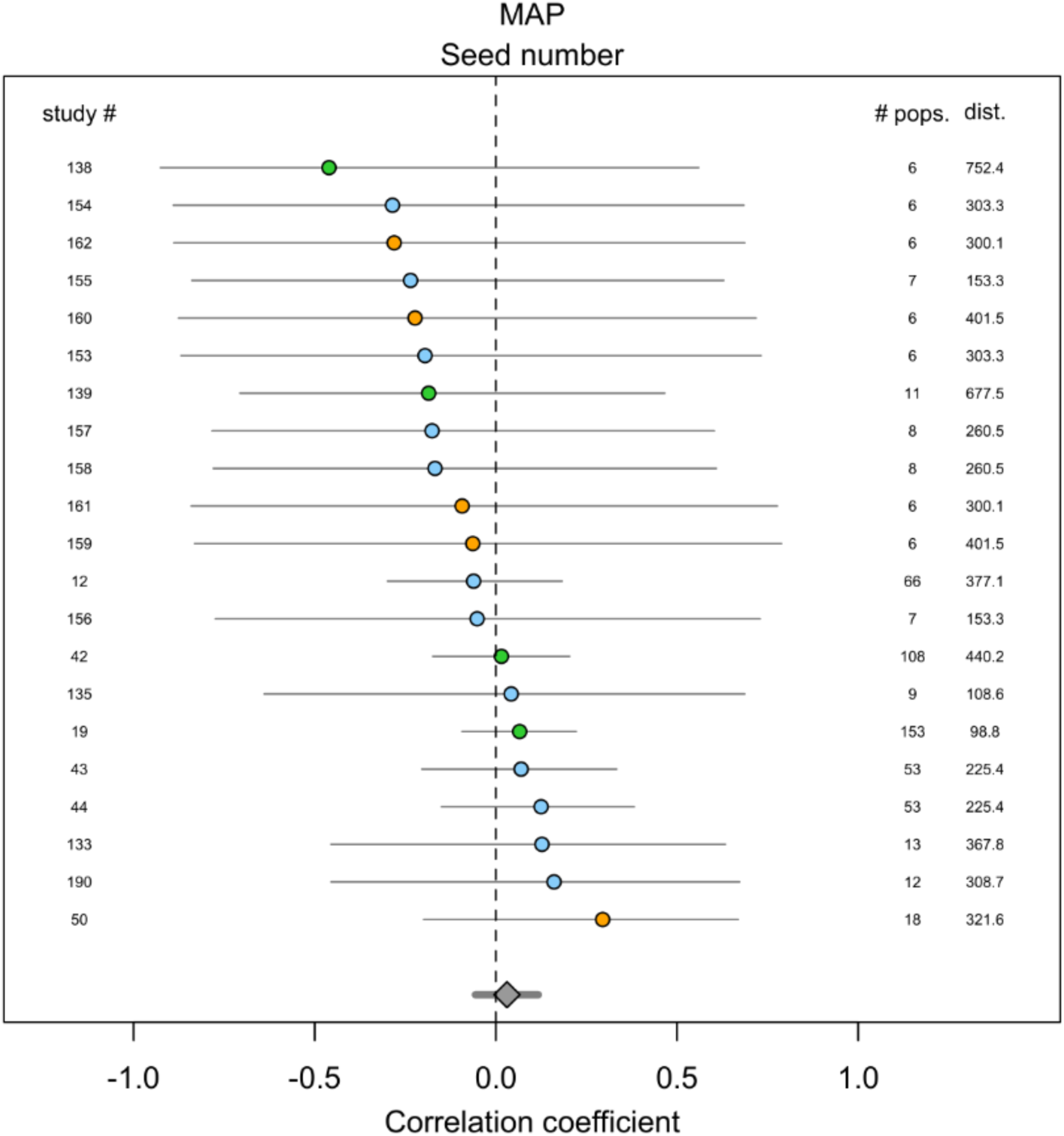
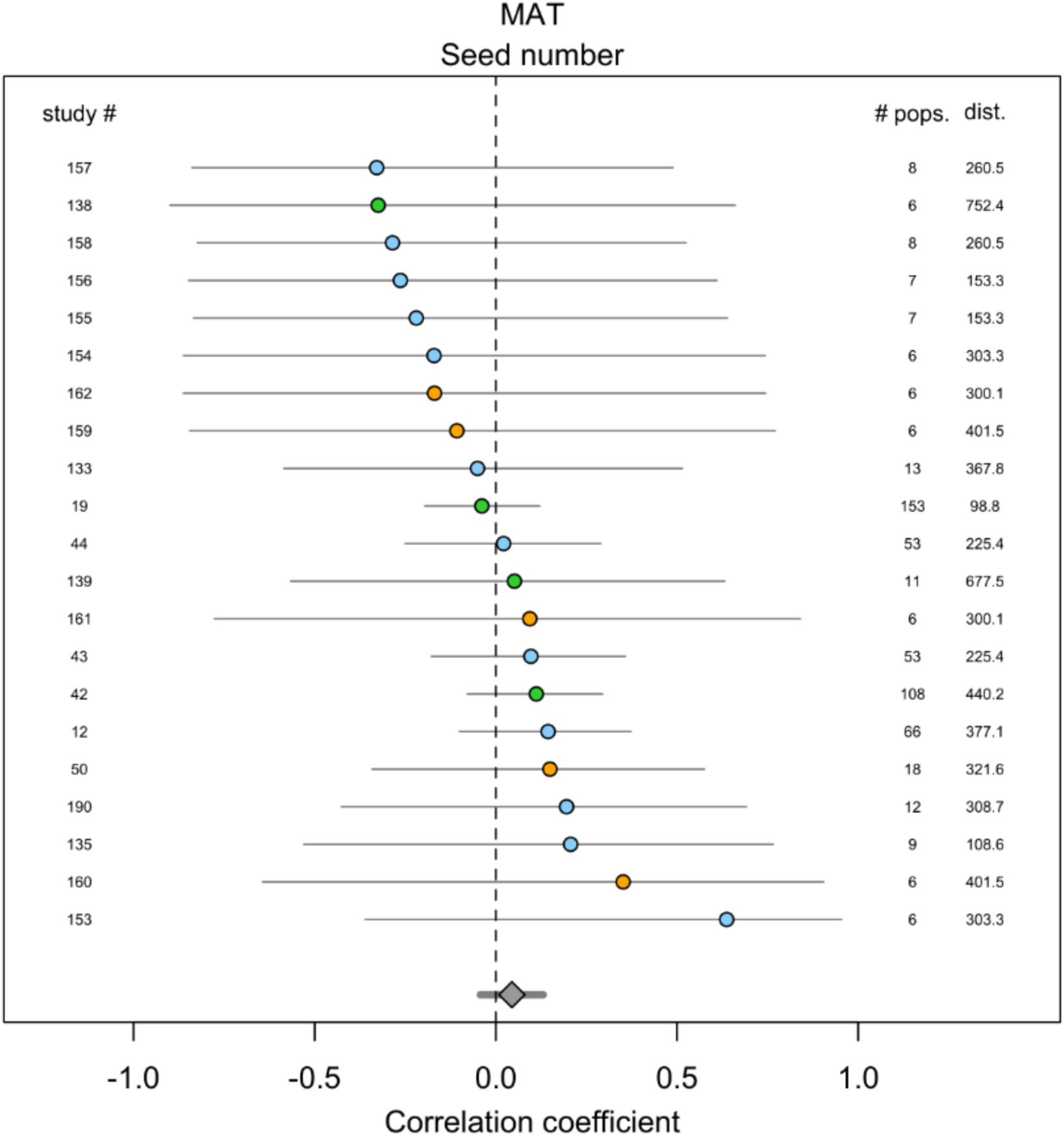
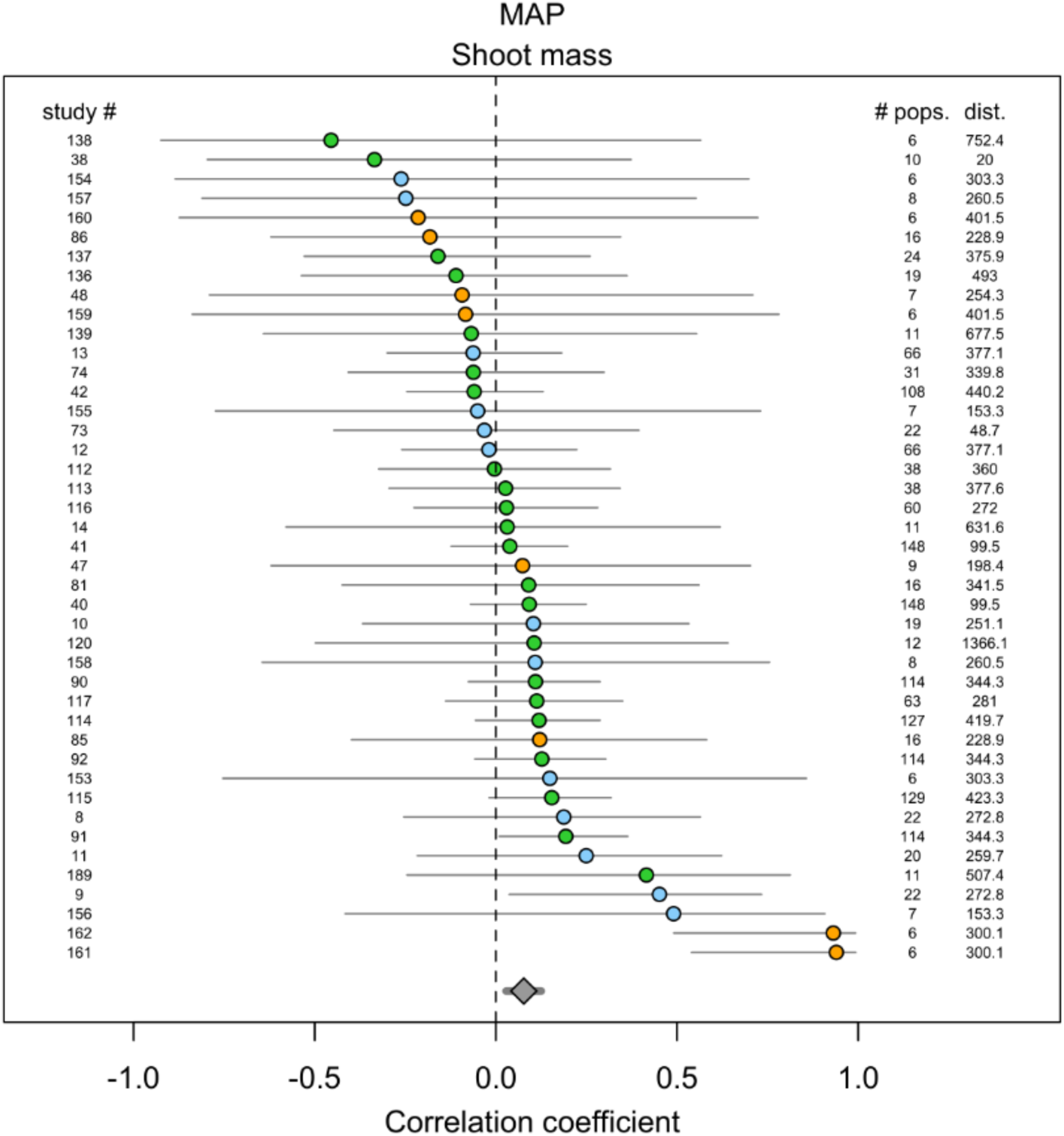
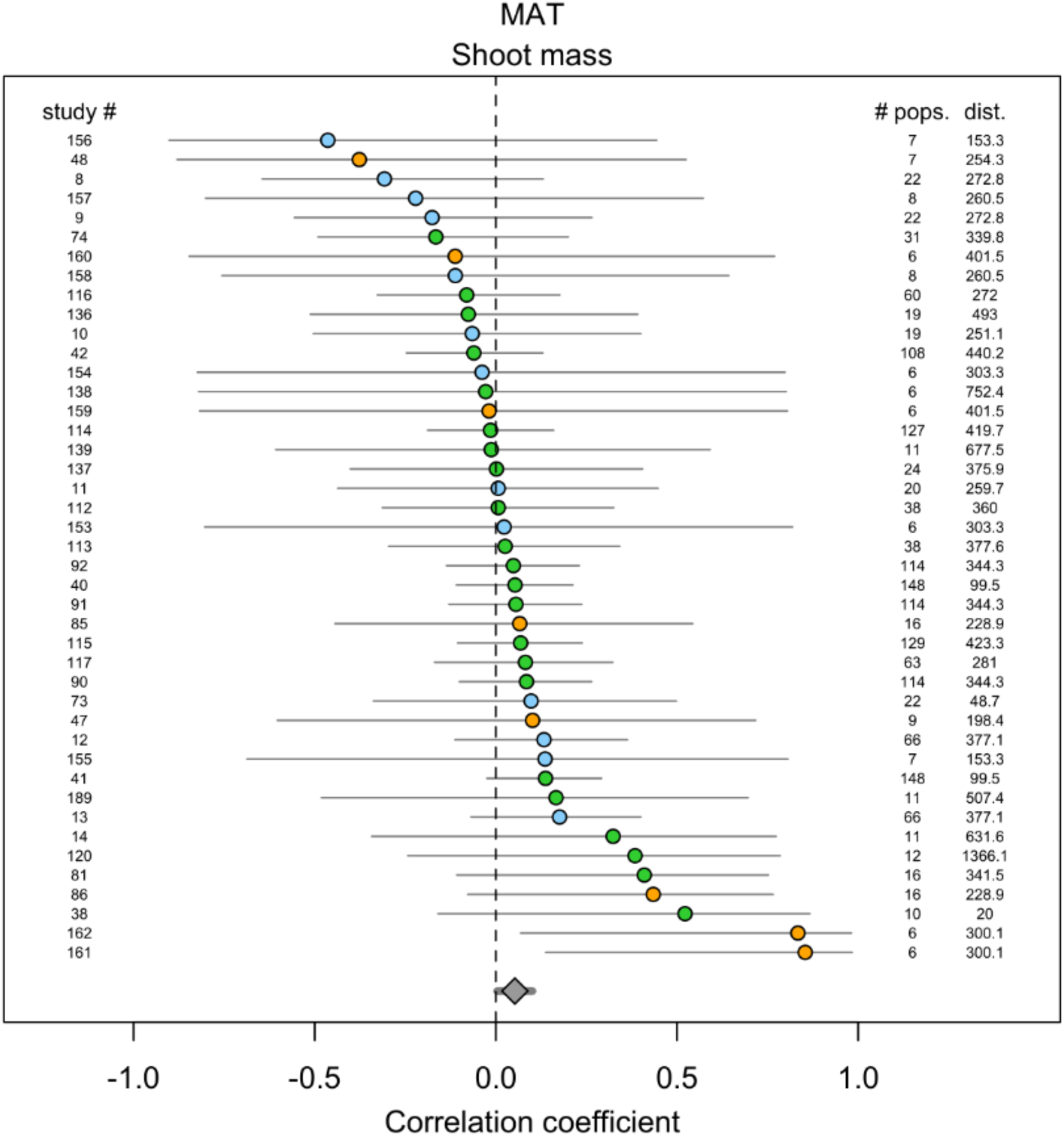
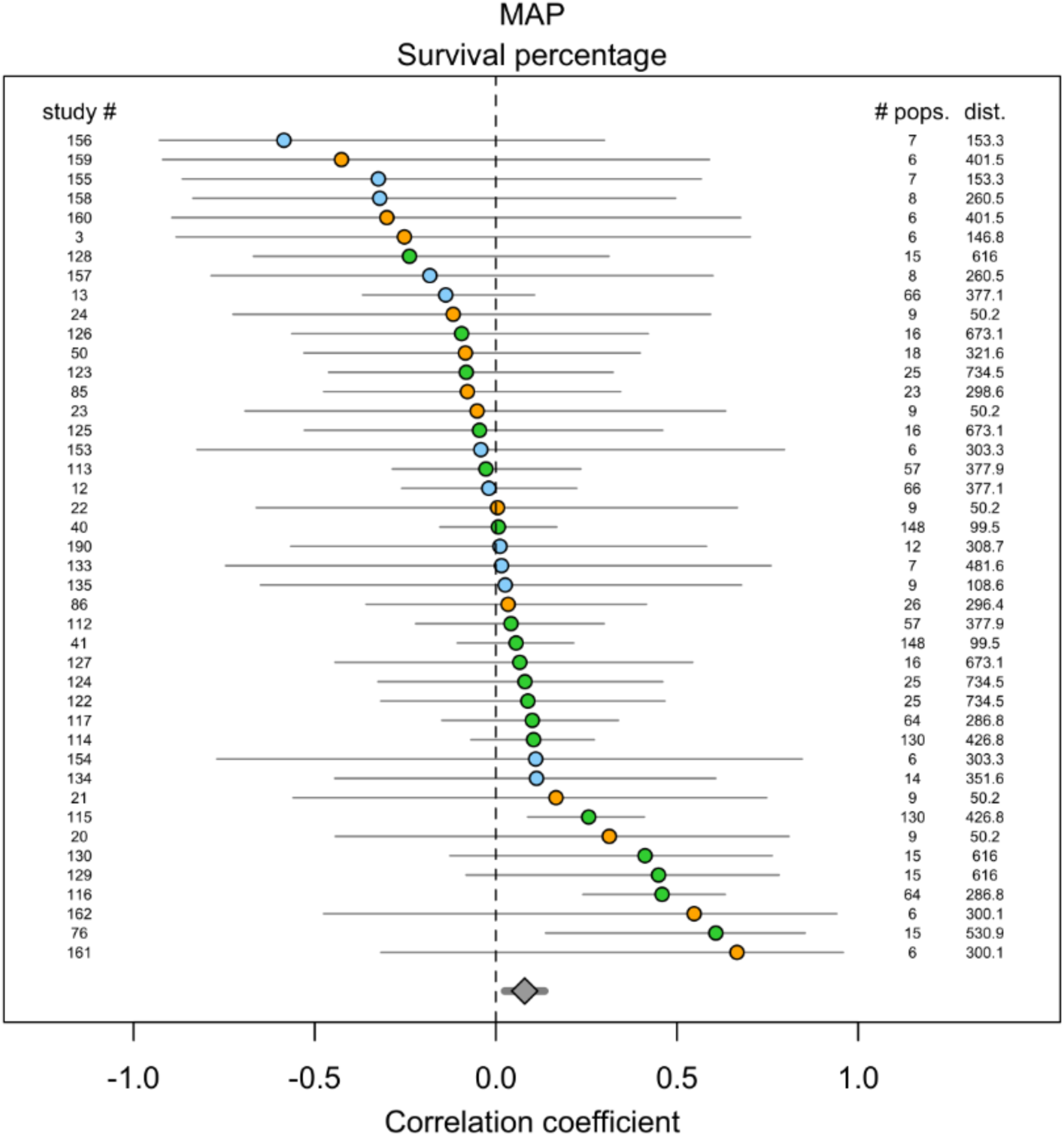
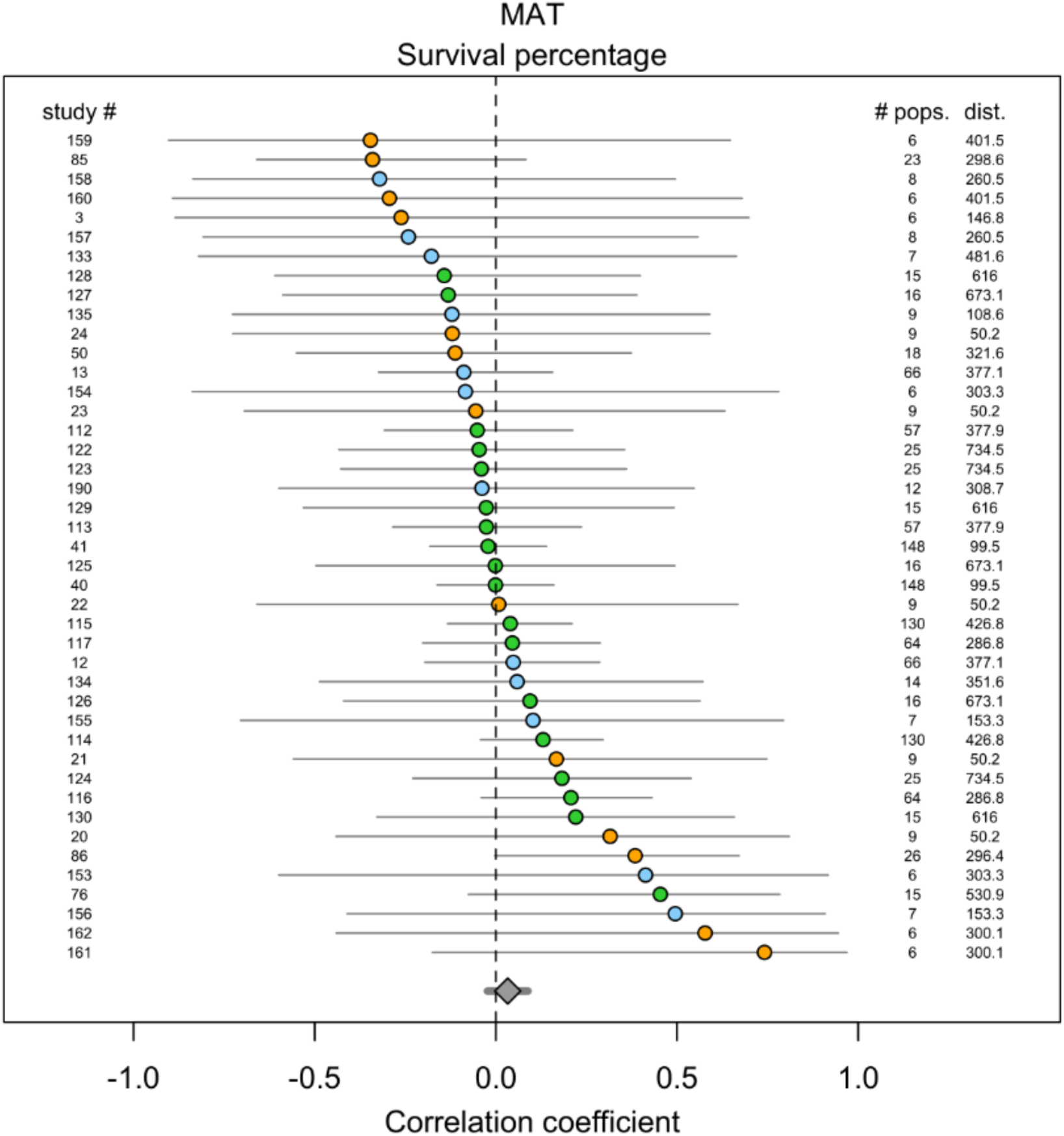
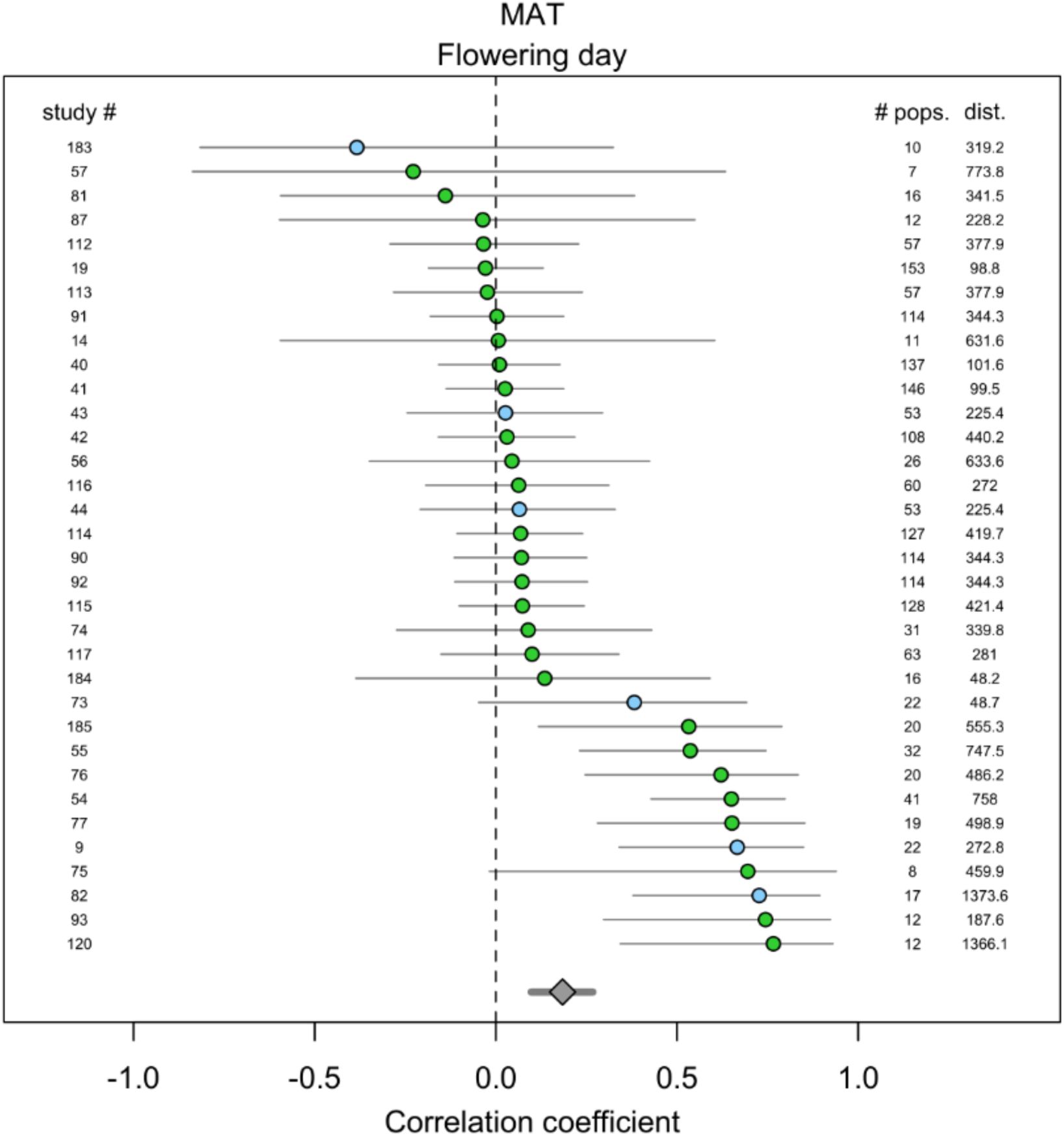
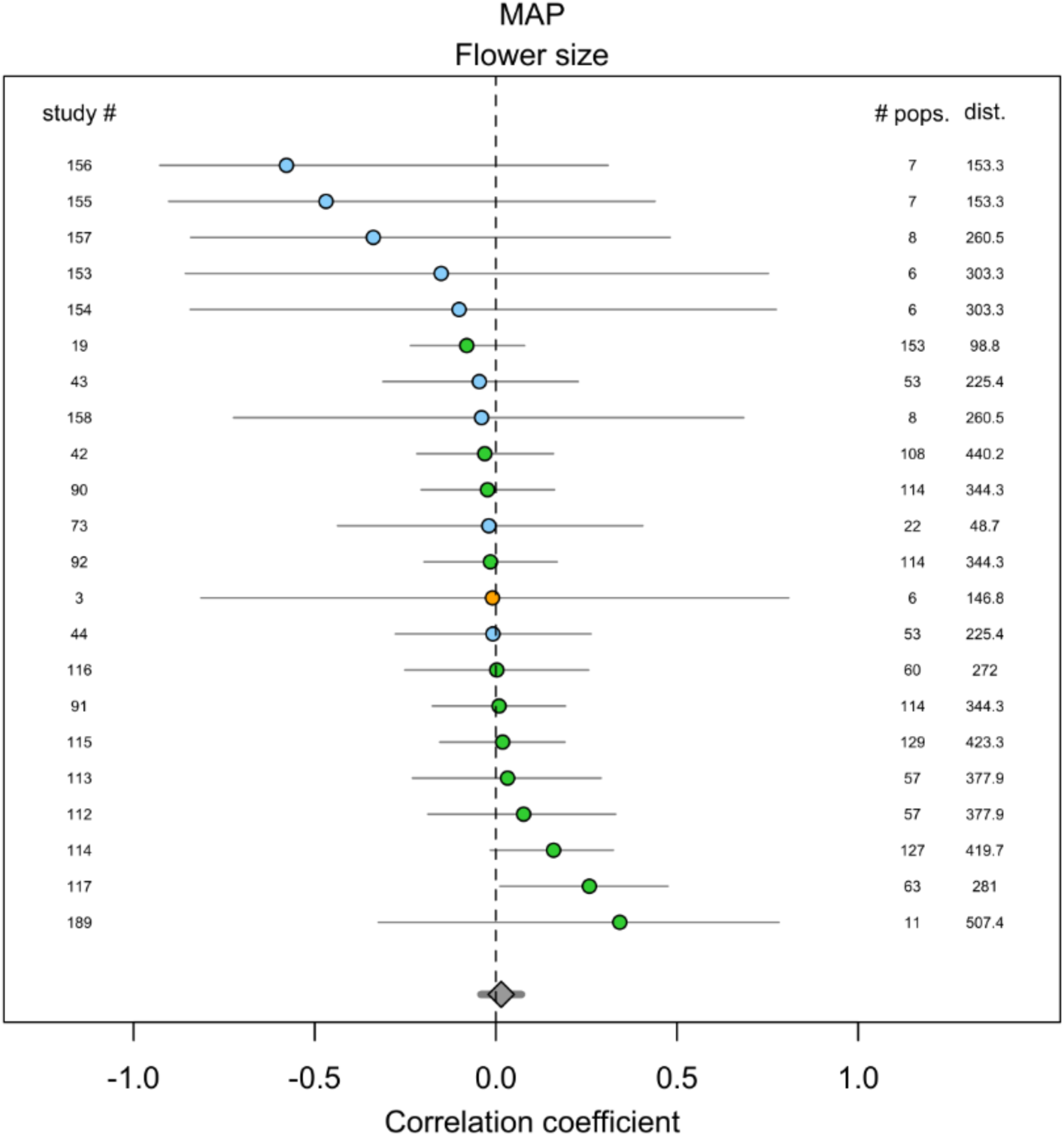
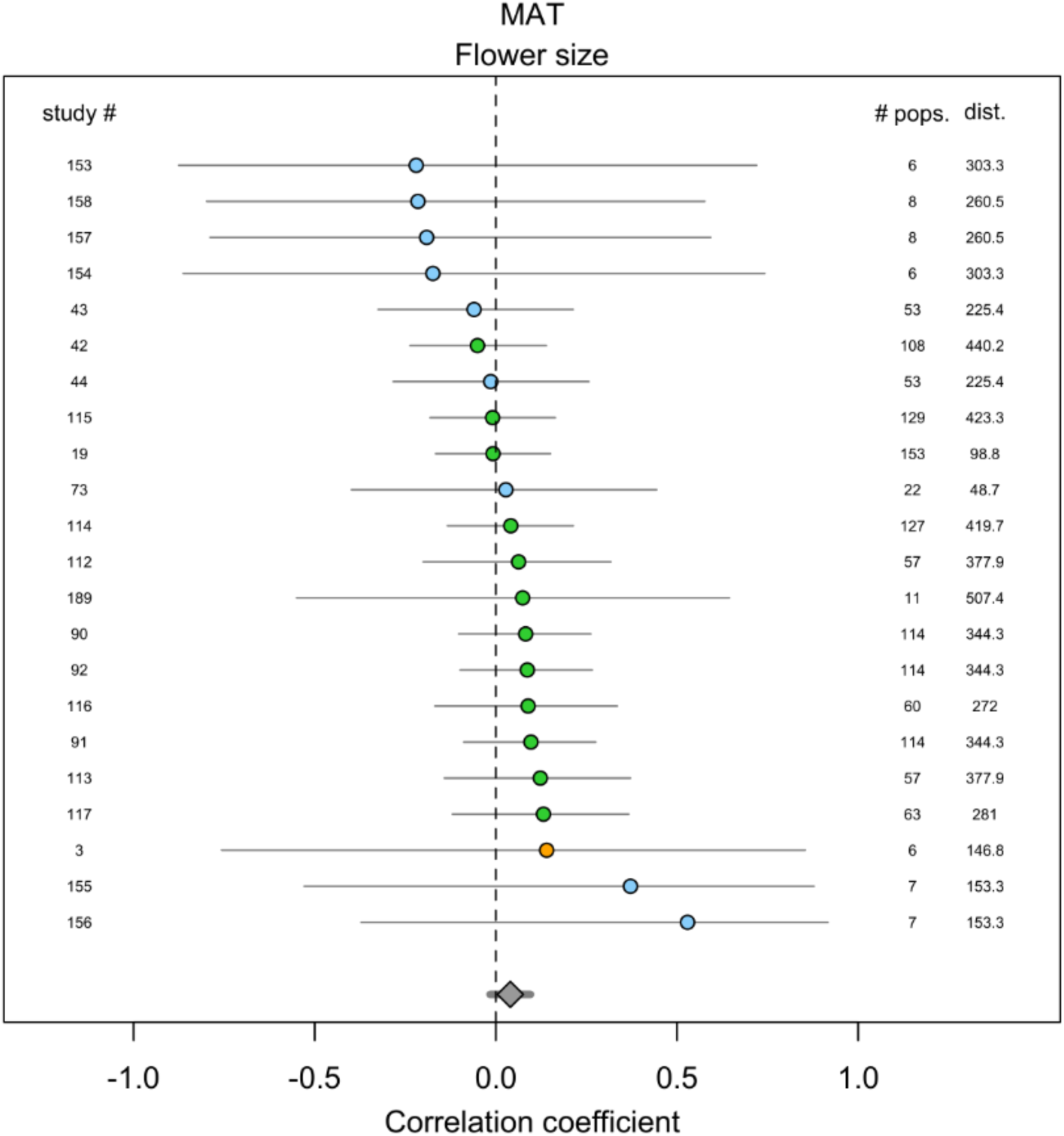
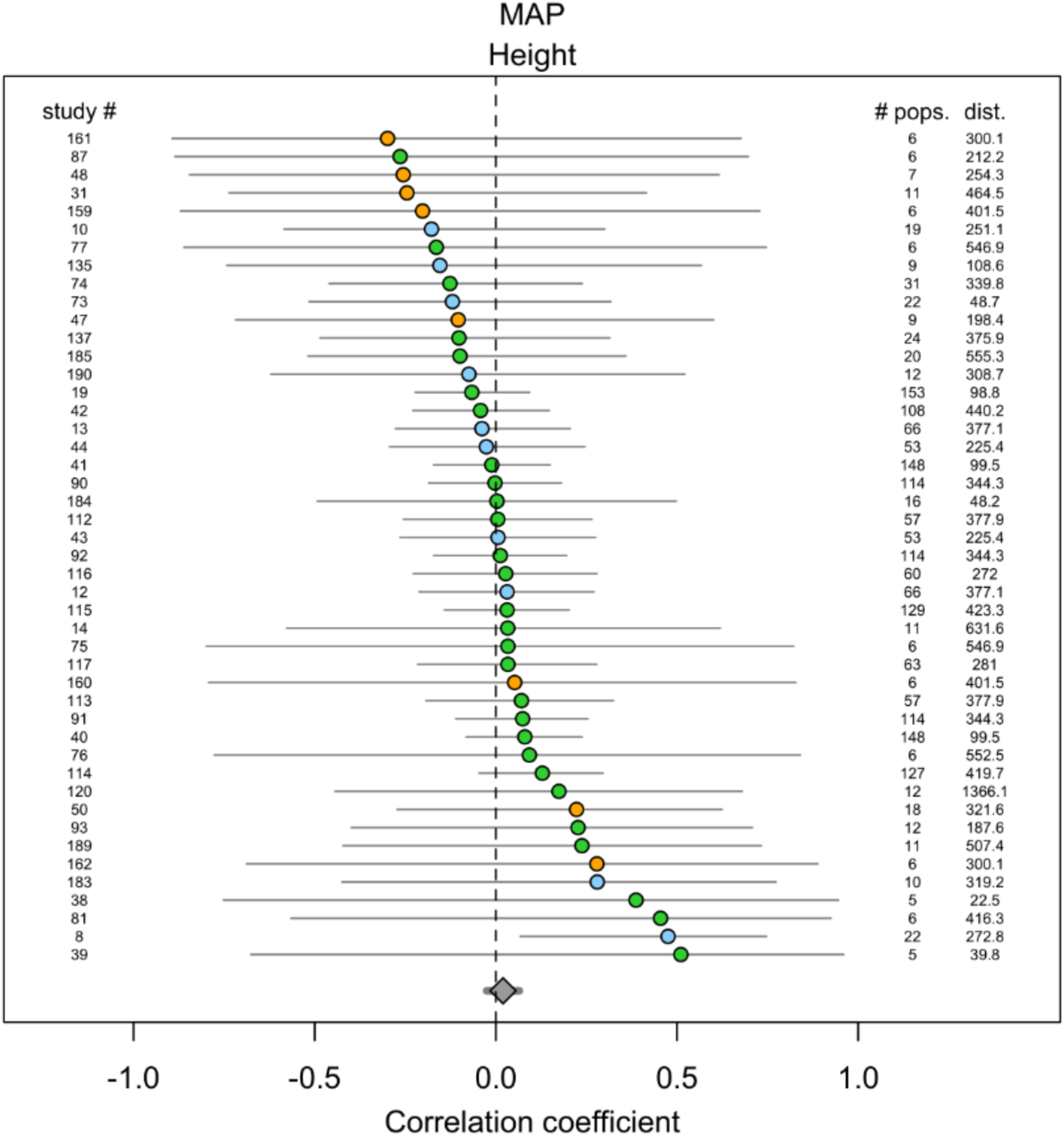
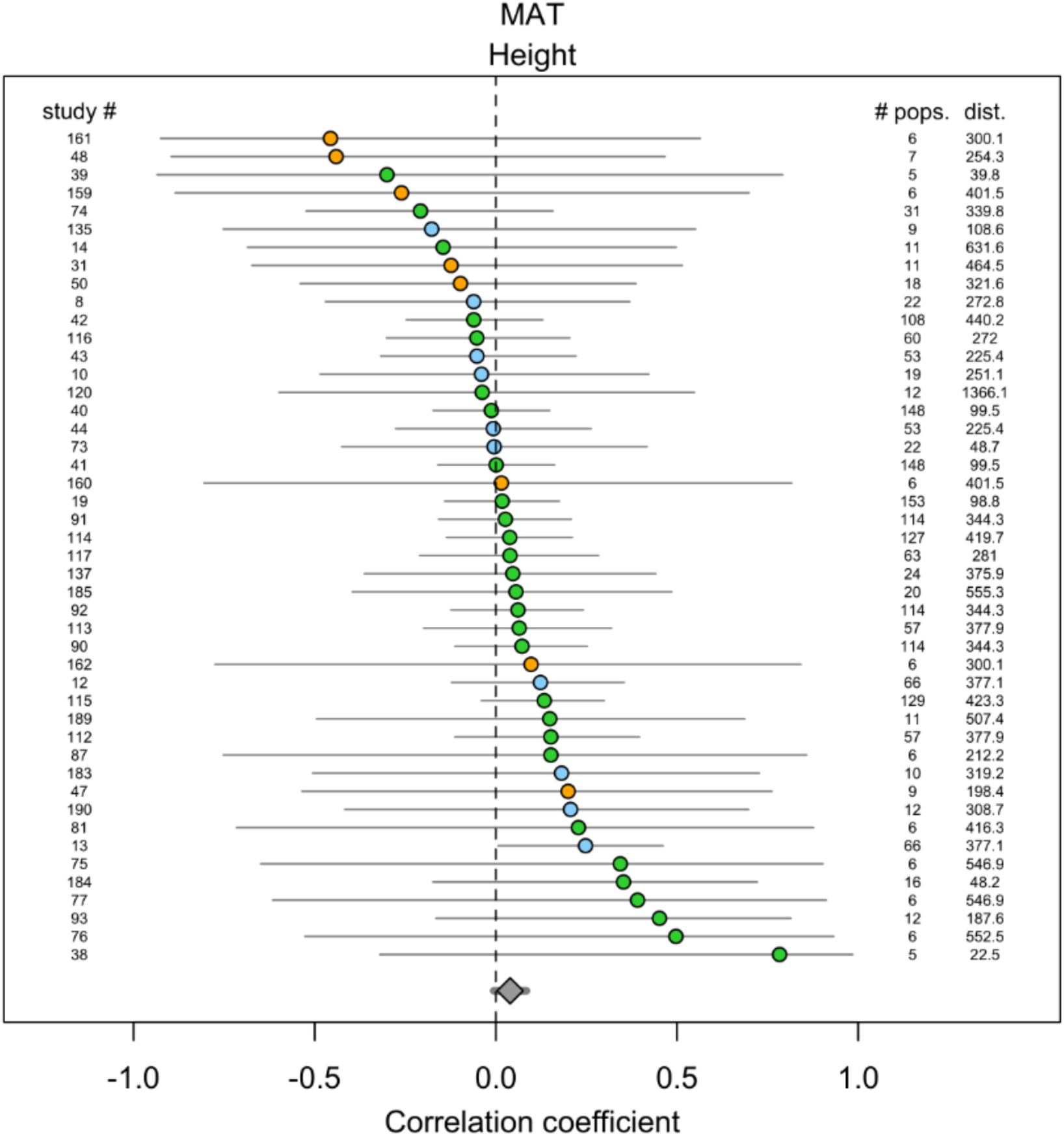
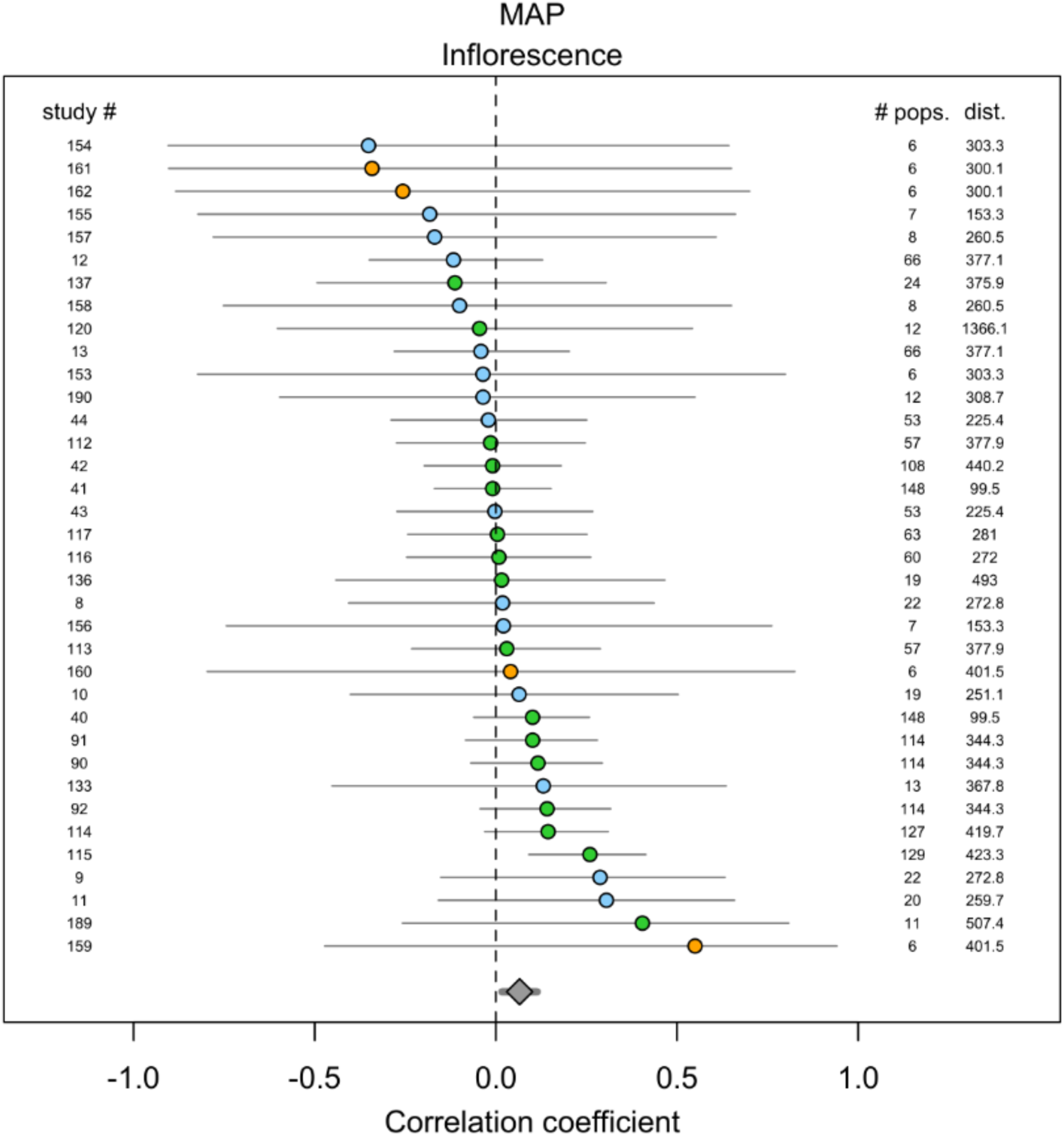
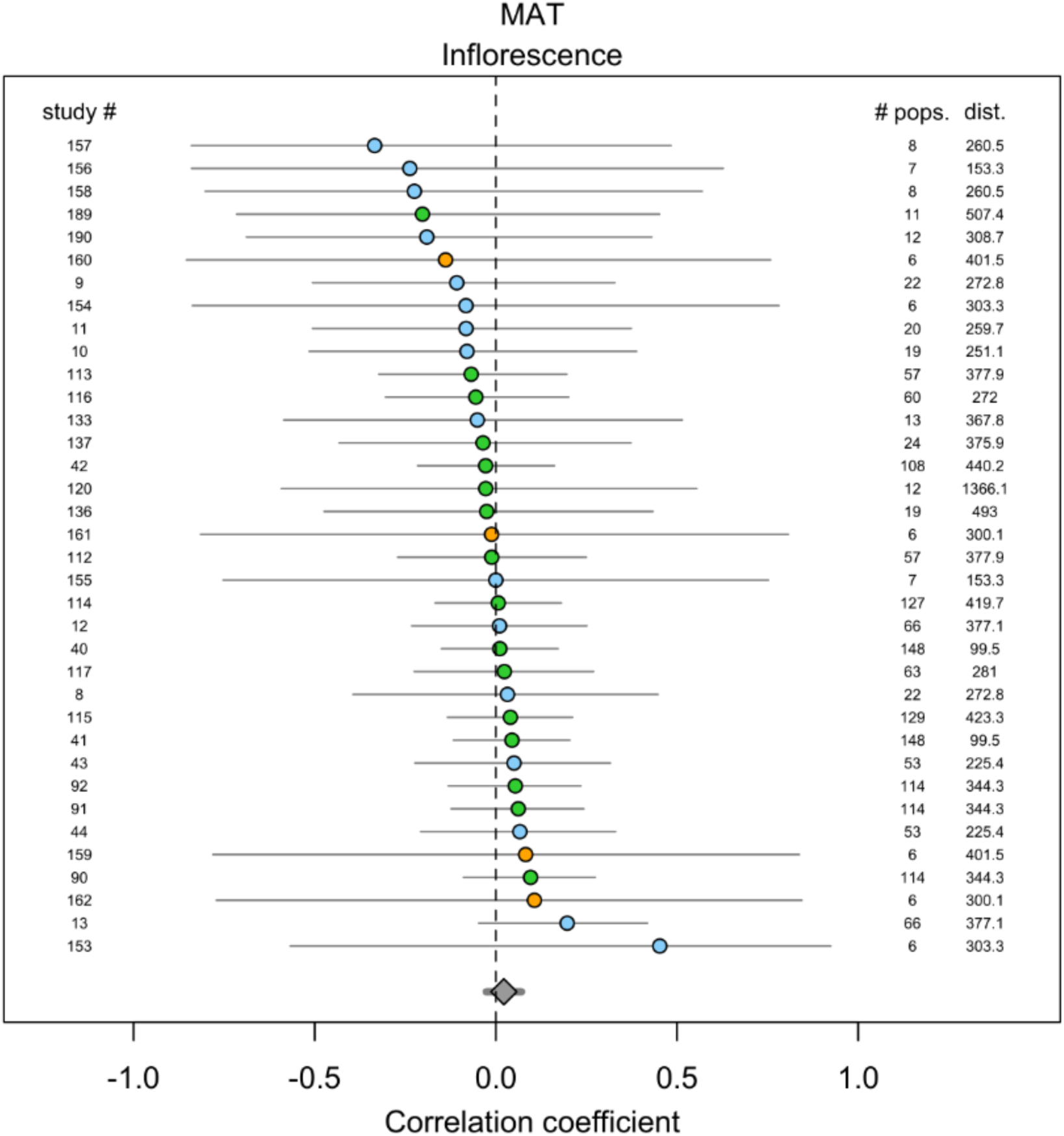
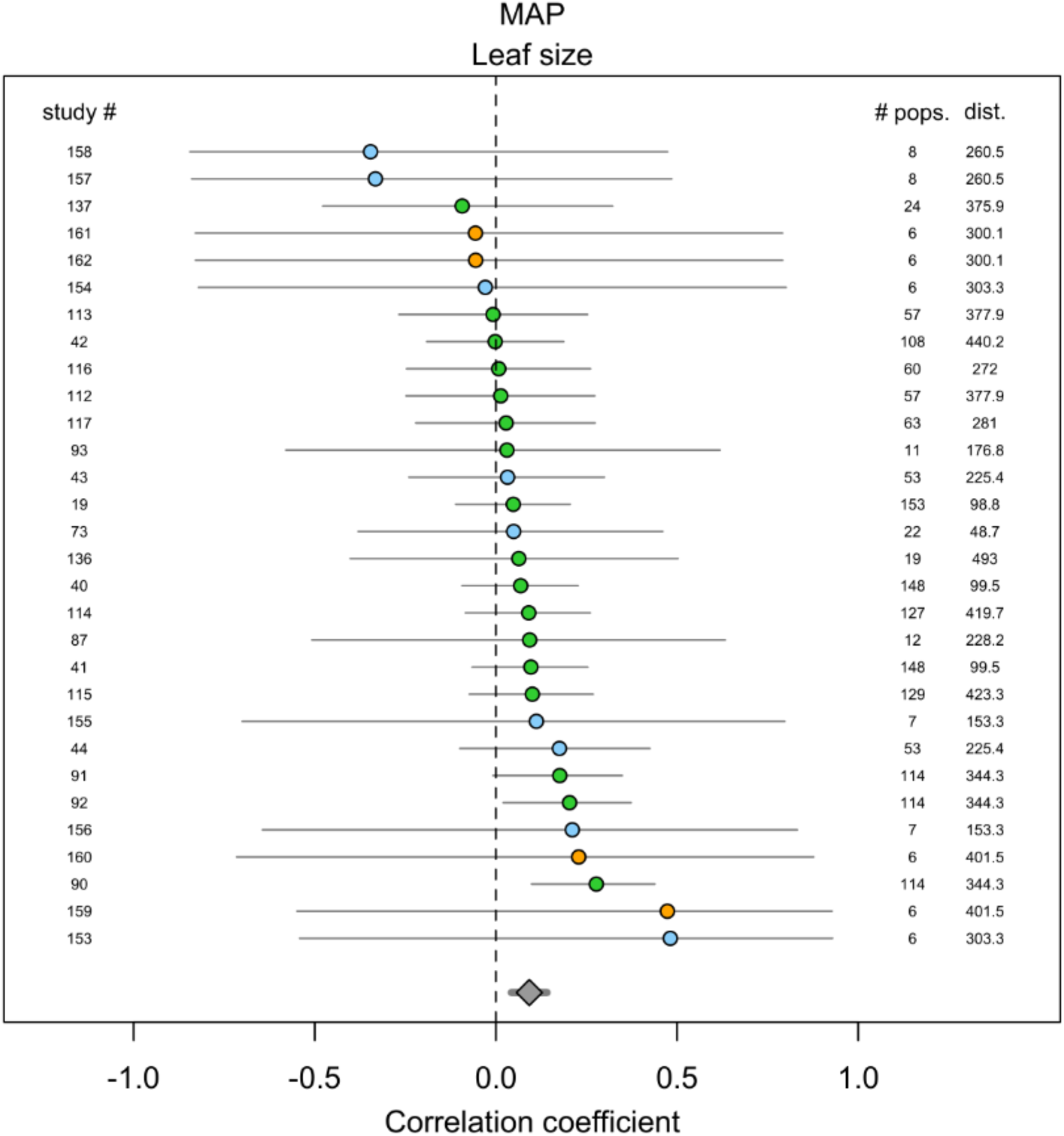
For each trait/environment correlation (16 combinations), the results of correlation coefficients (with 95% confidence intervals) for the pairwise comparisons, for each population in each experiment, between the difference in phenotypic trait and environmental characteristic at the collection location, while controlling for geographic distance among populations (see methods). MAT = Mean Annual Temperature, MAP = Mean Annual Precipitation. Study number identifies the particular experiment in the quantitative comparison dataset in the electronic supplementary material, “# pops.” is the number of populations included in each study, and “dist.” is the average pairwise distance between populations (in km). Grasses are shown in green, shrubs in blue, and forbs in orange. The overall effect size and confidence intervals, across all studies, is shown in gray at the bottom of each figure. See main text, Table 1, for descriptions of phenotypic traits.

