## appendix 2 for "Strong Patterns of Intraspecific Variation and Local Adaptation in Great Basin Plants Revealed Through a Review of 75 Years of Experiments" (from another version of this preprint: version 1, 2018-12-26)

**Appendix 2. Summary of literature and available datasets**

The data collected and generated by this study (Baughman *et al.*, 2019), as well as the list of publications that were involved in each part of this study, are provided so that additional questions may be addressed and for other applications. We encourage such additional analyses.

*Summary of literature*

Appendix 2 Table 1. Summary, by species, of the literature included in this study, including lifeform (F = forb, G = grass, S = shrub, T = tree), counts of studies, experiments, unique populations, and experiments by type (LAB = laboratory, GH = greenhouse, CG = outdoor common garden, RT = outdoor reciprocal transplant), the incidence of each signature of local adaptation (1 = differences among populations, 2 = trait/environment correlations, 3 = higher performance of local than nonlocal population in local’s environment), counts of experiments used in the quantitative comparison of trait-by-environment associations (QC), and a list of traits used in the QC. See footnotes for additional information.

[Appendix 2 Table 1 uploaded separately as .xlsx file]

*Available datasets*

Data have been uploaded to Dryad at <http://datadryad.org/review?doi=doi:10.5061/dryad.m57d386> (Baughman *et al.*, 2019). Several datasets are available. The “Summary and signature scores” dataset includes all of the studies and experiments and summarizes literature categorization as well as scores and associated information for each of the signatures of local adaptation. The “Trait scores” dataset includes basic study categorization as well as information that indicated which phenotypic traits (for signatures 1 and 2) and environmental variables (for signature 2) were involved in each of the signatures of local adaptation. The “Quantitative comparison” dataset includes all of the data used to conduct the quantitative comparison of trait-by-environment associations, and lists population-specific mean values for our priority traits for all studies for which such data were available, the latitude and longitude of each population origin, and extensive climate information for each origin generated with the ClimateNA v5.10 software package based on methodology described by Wang et al. (2016). The “Location data” dataset lists all outdoor gardens and population origin coordinates and elevations for which authors gave this information, as well as those for which we could confidently generate it. For descriptions of each column in each of these datasets, refer to the “Data Dictionary” file.

*Bibliography of reviewed literature*

A list of all the literature used in any of the datasets is provided below. Following each citation is a set of codes in brackets indicating which parts of our study the publication was used in. Codes S1, S2, and S3 indicate that at least one of the “experiments” in the given publication was used to generate a score for signatures 1, 2, and 3, and code QC indicates the publication (or the data summarized in it, even if not available from the publication itself) was used in analyses for the quantitative comparison of trait-by-environment associations. Note that some published studies were scored as multiple experiments for multiple species.

Angert, A. L., and D. W. Schemske. 2005. “The Evolution of Species’ Distributions: Reciprocal Transplants across the Elevation Ranges of Mimulus Cardinalis and M. Lewisii.” Evolution 59 (8):1671–84. <https://doi.org/10.1554/05-107.1>. [S1]

Atwater, Daniel Z., and Ragan M. Callaway. 2015. “Testing the Mechanisms of Diversity-Dependent Overyielding in a Grass Species.” Ecology 96 (12):3332–42. <https://doi.org/10.1890/15-0889.1>. [S1, QC]

Barker, Jr, and Cm McKell. 1986. “Differences in Big Sagebrush (Artemisia Tridentata) Plant Stature along Soil-Water Gradients: Genetic Components.” Journal of Range Management 39 (2):147–51. <http://www.jstor.org/stable/10.2307/3899288>. [S1, QC]

Barnes, Melanie G, and Diane L Marshall. 2009. “The Effect of Plant Source Location on Restoration Success: A Reciprocal Transplant Experiment with Winterfat (Krascheninnikovia Lanata).” Ecology. <https://doi.org/3390801>. [S1, S2, S3, QC]

Baughman, Owen W., Susan E. Meyer, Zachary T. Aanderud, and Elizabeth A. Leger. 2016. “Cheatgrass Die-Offs as an Opportunity for Restoration in the Great Basin, USA: Will Local or Commercial Native Plants Succeed Where Exotic Invaders Fail?” Journal of Arid Environments 124:193–204. <https://doi.org/10.1016/j.jaridenv.2015.08.011>. [S1, S3, QC]

Beckstead, Julie, Susan E. Meyer, and Phil S. Allen. n.d. “Effects of Afterripening on Cheatgrass (Bromus Tectorum) and Squirreltail (Elymus Elymoides) Germination.” In: Roundy, Bruce A.; McArthur, E. Durant; Haley, Jennifer S.; Mann, David K., Comps. 1995. Proceedings: Wildland Shrub and Arid Land Restoration Symposium; 1993 October 19-21; Las Vegas, NV. Gen. Tech. Rep. INT-GTR-315. Ogden, UT: U.S.D.A. [S1, S2]

Bergum, Karin E., Ann L. Hild, and Brian A. Mealor. 2010. “Phenotypes of Two Generations of Sporobolus Airoides Seedlings Derived from Acroptilon Repens-Invaded and Non-Invaded Grass Populations.” Restoration Ecology 20 (2):227–33. <https://doi.org/10.1111/j.1526-100X.2010.00754.x>. [S1, S2]

Bhattarai, Kishor, B. Shaun Bushman, Douglas A. Johnson, and John G. Carman. 2010. “Phenotypic and Genetic Characterization of Western Prairie Clover Collections from the Western United States.” Rangeland Ecology and Management 63 (6):696–706. <https://doi.org/10.2111/REM-D-10-00008.1>. [S1, S2, QC]

Bhattarai, Kishor, Douglas A. Johnson, Thomas A. Jones, Kevin J. Connors, and Dale R. Gardner. 2008. “Physiological and Morphological Characterization of Basalt Milkvetch (Astragalus Filipes): Basis for Plant Improvement.” Rangeland Ecology and Management 61 (4):444–55. <https://doi.org/10.2111/08-011.1>. [S1, S2, QC]

Bhattarai, Kishor, B. Shaun Bushman, Douglas A. Johnson, and John G. Carman. 2011. “Searls Prairie Clover (Dalea Searlsiae) for Rangeland Revegetation: Phenotypic and Genetic Evaluations.” Crop Science 51 (2):716–27. <https://doi.org/10.2135/cropsci2010.07.0387>. [S1, S2, QC]

Bleak, A.T., and Neil C. Frischknecht. 1965. “Problems in Artificial and Natural Revegetation of the Arid Shadscale Vegetation Zone of Utah and Nevada.” Journal of Range Management, 59–65. [S1]

Bohmont, B.L., and Robert Lang. 1957. “Some Variations in Morphological Characteristics and Palatability among Geographic Strains of Indian Ricegrass.” Journal of Range Management, 127–31.[S1]

Booth, D. Terrance. 1992. “Seedbed Ecology of Winterfat: Imbibition Temperature Affects Post-Germination Growth.” Journal of Range Management 45 (2):159–64. [S1]

Booth, D. Terrance, Charles G. Howard, and Charles E. Mowry. 1980. “‘Nezpar’ Indian Ricegrass: Description, Justification for Release, and Recommendations for Use.” Rangelands Archives 2 (2):53–54. [S1]

Brabec, Martha M., Matthew J. Germino, Douglas J. Shinneman, David S. Pilliod, Susan K. McIlroy, and Robert S. Arkle. 2015. “Challenges of Establishing Big Sagebrush (Artemisia Tridentata) in Rangeland Restoration: Effects of Herbicide, Mowing, Whole-Community Seeding, and Sagebrush Seed Sources.” Rangeland Ecology and Management 68 (5):432–35. <https://doi.org/10.1016/j.rama.2015.07.001>. [S1, S3, QC]

Brouillette, Larry C., Chase M. Mason, Rebecca Y. Shirk, and Lisa A. Donovan. 2014. “Adaptive Differentiation of Traits Related to Resource Use in a Desert Annual along a Resource Gradient.” New Phytologist 201 (4):1316–27. <https://doi.org/10.1111/nph.12628>. [S1, S2]

Butterfield, Bradley J., and Troy E. Wood. 2015. “Local Climate and Cultivation, but Not Ploidy, Predict Functional Trait Variation in Bouteloua Gracilis (Poaceae).” Plant Ecology 216 (10):1341–49. <https://doi.org/10.1007/s11258-015-0510-8>. [S1, S2]

Chabot, Brian F., and W. D. Billings. 1972. “Origins and Ecology of the Sierran Alpine Flora and Vegetation.” Ecological Monographs 42 (2):163–99. <https://doi.org/10.2307/1942262>. [S2]

Chaney, Lindsay, Bryce A. Richardson, and Matthew J. Germino. 2017. “Climate Drives Adaptive Genetic Responses Associated with Survival in Big Sagebrush (Artemisia Tridentata).” Evolutionary Applications 10 (4):313–22. <https://doi.org/10.1111/eva.12440>. [S1, S2, QC]

Chapin, F. S. III, and C. M. Chapin. 1981. “Ecotypic Differentiation of Growth Processes in Carex Aquatilis along a Latitudinal and Local Gradients.” Ecology 62 (4):1000–1009. <https://doi.org/10.2307/1936999>. [S1, S2, S3, QC]

Clark, Lesley D., and Neil E. West. 1971. “Further Studies of Eurotia Lanata Germination in Relation to Salinity.” The Southwestern Naturalist, 371–75. [S1]

Clary, Warren P. 1975. “Ecotypic Adaptation in Sitanion Hystrix.” Ecology 56 (6):1407–15. [S1, S2, QC]

Clary, Warren P. 1979. “Variation in Leaf Anatomy and CO₂ Assimilation in Sitanion Hystrix Ecotypes.” The Great Basin Naturalist, 427–32. [S1, S2]

Clauss, M. J., and D. L. Venable. 2000. “Seed Germination in Desert Annuals: An Empirical Test of Adaptive Bet Hedging.” The American Naturalist 155 (2):168–86. <https://doi.org/10.1086/303314>. [S1, S2, QC]

Cook, Stanton A, and Michael P Johnson. 1968. “Adaptation to Heterogeneous Environments. I. Variation in Heterophylly in Ranunculus Flammula L.” Evolution 22 (3):496–516. <https://doi.org/10.1111/j.1558-5646.1968.tb03988.x>. [S1, S2]

Dewey, Douglas R. 1960. “Salt Tolerance of Twenty-Five Strains of Agropyron.” Agronomy Journal 52 (11):631–35. <https://doi.org/10.2134/agronj1960.00021962005200110006x>. [S1, QC]

Doede, David L. 2005. “Genetic Variation in Broadleaf Lupine (Lupinus Latifolius) on the Mt Hood National Forest and Implications for Seed Collection and Deployment.” Native Plants Journal 6 (1):36–48. <https://doi.org/10.1353/npj.2005.0018>. [S1, S2]

Doescher, P.S. 1983. “Phyto-Edaphic Relationships and Ecotypic Development of Festuca Idahoensis in Eastern Oregon Habitat Types of Artemisia Tridentata.” Ph.D. Dissertation, Oregon State University, Corvallis. [S1, S2, QC]

Emery, R. J. N., C. C. Chinnappa, and J. G. Chmielewski. 1994. “Specialization, Plant Strategies, and Phenotypic Plasticity in Populations of Stellaria Longipes Along an Elevational Gradient.” International Journal of Plant Sciences 155 (2):203–19. <https://doi.org/10.2307/2995565>. [S1, QC]

Erickson, Vicky J, Nancy L Mandel, and Frank C Sorenson. 2004. “Landscape Patterns of Phenotypic Variation and Population Structuring in a Selfing Grass, Elymus Glaucus (Blue Wildrye).” Canadian Journal of Botany 82:1776–89. <https://doi.org/10.1139/B04-141>. [S1, S2, QC]

Evans, Raymond A, and James A Young. 1990. “Survival and Growth of Big Sagebrush (Artemisia Tridentata) Plants in Reciprocal Gardens.” Weed Science 38 (3):215–19. <https://doi.org/10.2307/4045014>. [S1, S3, QC]

Ferguson, Scot D., Elizabeth A. Leger, Jun Li, and Robert S. Nowak. 2015. “Natural Selection Favors Root Investment in Native Grasses during Restoration of Invaded Fields.” Journal of Arid Environments 116:11–17. <https://doi.org/10.1016/j.jaridenv.2015.01.009>. [S1]

Ferrero-Serrano, Ángel, Ann L. Hild, and Brian A. Mealor. 2011. “Can Invasive Species Enhance Competitive Ability and Restoration Potential in Native Grass Populations?” Restoration Ecology 19 (4):545–51. <https://doi.org/10.1111/j.1526-100X.2009.00611.x>. [S1, S2]

Fisk, Matthew R. 2016. “Dynamics of Cold Hardiness Accumulation and Loss in the Great Basin Native Species Eriogonum Umbellatum.” Ph.D. Dissertation, University of Idaho, Boise. [S1, S2]

Fitzsimmons, Kevin, Cynthia Lovely, and Edward Glenn. 1998. “Growth Differences among Widely Separated Geographic Accessions of Fourwing Saltbush (Atriplex Canescens) in the Great Basin Desert, New Mexico, USA.” Arid Soil Research and Rehabilitation 12 (2):87–94. <https://doi.org/10.1080/15324989809381501>. [S1, S2, QC]

Fonseca, Carolina, Erin Espeland, and James W. Baxter. 2014. “Patterns of Population Differentiation in Early Traits of Development in Elymus Glaucus: Implications for Restoration.” Ecological Restoration 32 (4):388–95. <https://doi.org/10.3368/er.32.4.388>. [S1]

Galen, Candace, Joel S Shore, and Hudson Deyoe. 1991. “Ecotypic Divergence in Alpine Polemonium Viscosum: Genetic Structure, Quantitative Variation, and Local Adaptation.” Evolution 45 (455):1218–28. <https://doi.org/10.2307/2409729>. [S1, S2, S3, QC]

Goergen, Erin M., Elizabeth A. Leger, and Erin K. Espeland. 2011. “Native Perennial Grasses Show Evolutionary Response to Bromus Tectorum (Cheatgrass) Invasion.” PLoS ONE 6 (3). <https://doi.org/10.1371/journal.pone.0018145>. [S1, S2]

Hall, J.W., D.G. Stout, and B. Brooke. 1990. “Effect of Seed Source on Growth of Giant Wildrye (Elymus Cinereus) at Two Elevations in Interior British Columbia.” Canadian Journal of Plant Science 70 (2):551–54. [S1, S3, QC]

Hardegree, Stuart P., Thomas A. Jones, Frederick B. Pierson, Patrick E. Clark, and Gerald N. Flerchinger. 2008. “Dynamic Variability in Thermal-Germination Response of Squirreltail (Elymus Elymoides and Elymus Multisetus).” Environmental and Experimental Botany 62 (2):120–28. <https://doi.org/10.1016/j.envexpbot.2007.07.010>. [S2]

Harmon, Dan, and Charlie D. Clements. 2016. “Characteristics That Determine a Successful Squirreltail (Elymus Elymoides).” In Poster Session Presented at the Society for Range Management, Corpus Christi, TX. [S1]

Hergert, Holden J., Brian A. Mealor, and Andrew R. Kniss. 2015. “Inter-and Intraspecific Variation in Native Restoration Plants for Herbicide Tolerance.” Ecological Restoration 33 (1):74–81. <https://doi.org/10.3368/er.33.1.74>. [S1]

Hild, A L, J M Muscha, and N L Shaw. 2007. “Emergence and Growth of Four Winterfat Accessions in the Presence of the Exotic Annual Cheatgrass.” Proceedings: Shrubland Dynamics-Fire and Water; 2004 August 10-12; Lubbock, TX., no. 47:0–147. [S1]

Hintz, Lisa, M.M. Eshelman, A. Foxx, T.E. Wood, and A. Kramer. 2016. “Population Differentiation in Early Life History Traits of Cleome Lutea Var. Lutea in the Intermountain West.” Western North American Naturalist 76 (1):6–17. [S1, S2]

Horning, Matthew E., Theresa R. McGovern, Dale C. Darris, Nancy L. Mandel, and Randy Johnson. 2010. “Genecology of Holodiscus Discolor (Rosaceae) in the Pacific Northwest, U.S.A.” Restoration Ecology 18 (2):235–43. <https://doi.org/10.1111/j.1526-100X.2008.00441.x>. [S1, S2]

Humphrey, L. David, and Eugene W. Schupp. 2002. “Seedling Survival from Locally and Commercially Obtained Seeds on Two Semiarid Sites.” Restoration Ecology 10 (1):88–95. <https://doi.org/10.1046/j.1526-100X.2002.10109.x>. [S1, S3, QC]

Jaindl, Raymond G., Paul Doescher, Richard F. Miller, and Lee E. Eddleman. 1994. “Persistence of Idaho Fescue on Degraded Rangelands: Adaptation to Defoliation or Tolerance.” Journal of Range Management 47 (1):54. <https://doi.org/10.2307/4002841>. [S1, S2, QC]

Johnson, R C, V J Erickson, N L Mandel, J Bradley St Clair, and K W Vance-Borland. 2010. “Mapping Genetic Variation and Seed Zones for Bromus Carinatus in the Blue Mountains of Eastern Oregon, USA.” Botany 88 (8):725–36. <https://doi.org/10.1139/B10-047>. [S1, S2, QC]

Johnson, R C, B C Hellier, and K W Vance-Borland. 2013. “Genecology and Seed Zones for Tapertip Onion in the US Great Basin.” Botany-Botanique 91 (10):686–94. https://doi.org/DOI 10.1139/cjb-2013-0046. [S1, S2, QC]

Johnson, R. C., M. J. Cashman, and K. Vance-Borland. 2012. “Genecology and Seed Zones for Indian Ricegrass Collected in the Southwestern United States.” Rangeland Ecology and Management 65 (5):523–32. <https://doi.org/10.2111/REM-D-11-00165.1>. [S1, S2, QC]

Johnson, R. C., E. A. Leger, and Ken Vance-Borland. 2017. “Genecology of Thurber’s Needlegrass (Achnatherum Thurberianum [Piper] Barkworth) in the Western United States.” Rangeland Ecology and Management 70 (4):509–17. <https://doi.org/10.1016/j.rama.2017.01.004>. [S1, S2, QC]

Johnson, R. C., and Ken Vance-Borland. 2016. “Linking Genetic Variation in Adaptive Plant Traits to Climate in Tetraploid and Octoploid Basin Wildrye [Leymus Cinereus (Scribn. & Merr.) A. Love] in the Western U.S.” PLoS ONE 11 (2). <https://doi.org/10.1371/journal.pone.0148982>. [S1, S2, QC]

Johnson, Richard C., Matthew E. Horning, Erin K. Espeland, and Ken Vance-Borland. 2015. “Relating Adaptive Genetic Traits to Climate for Sandberg Bluegrass from the Intermountain Western United States.” Evolutionary Applications 8 (2):172–84. <https://doi.org/10.1111/eva.12240>. [S1, S2, QC]

Jones, T A, D C Nielson, J T Arredondo, and M G Redinbaugh. 2003. “Characterization of Diversity among 3 Squirreltail Taxa.” Journal of Range Management 56 (5):474–82. https://doi.org/Doi 10.2307/4003839. [S1, QC]

Jones, T.A. 2004. “Registration of Ribstone Indian Ricegrass Germplasm.” Crop Science 44 (3):1031–33. [S1]

Jones, T.A., D.C. Nielson, S.K. Caicco, G.A. Fenchel, and S. A. Young. 2005. “Registration of Star Lake Indian Ricegrass Germplasm.” Crop Science 45 (4):1666–67. [S1]

Jones, Thomas A., S.R. Winslow, S.D. Parr, and K.L. Memmott. 2010. “Notice of Release of White River Germplasm Indian Ricegrass.” Native Plants Journal 11 (2):133–36. [S1]

Kardol, P., J. R. De Long, and D. A. Wardle. 2014. “Local Plant Adaptation across a Subarctic Elevational Gradient.” Royal Society Open Science 1 (3):140141–140141. <https://doi.org/10.1098/rsos.140141>. [S1]

Kim, Eunsuk, and Kathleen Donohue. 2013. “Local Adaptation and Plasticity of Erysimum Capitatum to Altitude: Its Implications for Responses to Climate Change.” Journal of Ecology 101 (3):796–805. <https://doi.org/10.1111/1365-2745.12077>. [S1, S3, QC]

Kitchen, Stanley G. n.d. “Return of the Native: A Look at Select Accessions of North American Lewis Flax.” In: Roundy, Bruce A.; McArthur, E. Durant; Haley, Jennifer S.; Mann, David K., Comps. 1995. Proceedings: Wildland Shrub and Arid Land Restoration Symposium; 1993 October 19-21; Las Vegas, NV. Gen. Tech. Rep. INT-GTR-315. Ogden, UT: U.S.D.A. [S1]

Kitchen, Stanley G., and Loren St. John. 1996. “Release Documentation for Maple Grove Lewis Flax.” USDA NRCS Aberdeen Plant Materials Center, Aberdeen, Idaho. <https://www.nrcs.usda.gov/Internet/FSE_PLANTMATERIALS/publications/idpmcrn5639.pdf>. [S1, S2, QC]

Kramer, Andrea T. 2009. “Ecological Genetics of Penstemon in the Great Basin, USA.” Ph.D. Dissertation, University of Illinois at Chicago. [S1, S2]

Kramer, Andrea T., Daniel J. Larkin, and Jeremie B. Fant. 2015. “Assessing Potential Seed Transfer Zones for Five Forb Species from the Great Basin Floristic Region, USA.” Natural Areas Journal 35 (1):174–88. [S1, S2, QC]

Larsen, Eugene C. 1947. “Photoperiodic Responses of Geographical Strains of Andropogon Scoparius.” Botanical Gazette 109 (2):132–49. [S1, S2, QC]

Leger, Elizabeth A. 2008. “The Adaptive Value of Remnant Native Plants in Invaded Communities: An Example from the Great Basin.” Ecological Applications 18 (5):1226–35. [S1, S2]

Li, Dapeng, Ian T. Baldwin, and Emmanuel Gaquerel. 2015. “Navigating Natural Variation in Herbivory-Induced Secondary Metabolism in Coyote Tobacco Populations Using MS/MS Structural Analysis.” Proceedings of the National Academy of Sciences 112 (30):E4147–55. [S1, S2]

Liao, Huixuan, Priscila C.S. Gurgel, Robert W. Pal, David Hooper, and Ragan M. Callaway. 2016. “Solidago Gigantea Plants from Nonnative Ranges Compensate More in Response to Damage than Plants from the Native Range.” Ecology 97 (9):2355–63. <https://doi.org/10.1002/ecy.1481>. [S1, S2]

Link, Steven O., Jeffrey L. Smith, Jonathan J. Halvorson, and Harvey Bolton. 2003. “A Reciprocal Transplant Experiment within a Climatic Gradient in a Semiarid Shrub-Steppe Ecosystem: Effects on Bunchgrass Growth and Reproduction, Soil Carbon, and Soil Nitrogen.” Global Change Biology 9 (7):1097–1105. <https://doi.org/10.1046/j.1365-2486.2003.00647.x>. [S1, S3, QC]

Love, Stephen L, Robert R Tripepi, and Thomas Salaiz. 2014. “Influence of Harvest Timing and Storage Interval on Rabbitbrush Seed Germination, Emergence, and Viability.” Native Plants Journal (University of Wisconsin Press) 15 (2):98–108. <https://doi.org/10.1353/npj.2014.0017>. [S1]

Mann, Rebecca K. 2016. “Intraspecific Variation in the Response of Elymus Elymoides to Competition from Bromus Tectorum.” Ph.D. Dissertation, Utah State University, Logan. [S1, S2]

McArthur, E. Durant, Susan E. Meyer, and Darrel J. Weber. 1987. “Germination Rate at Low Temperature: Rubber Rabbitbrush Population Differences.” Journal of Range Management, 530–33. [S1, S2]

McArthur, E. Durant, Richard Stevens, and A. Clyde Blauer. 1983. “Growth Performance Comparisons among 18 Accessions of Fourwing Saltbush [Atriplex Canescens] at Two Sites in Central Utah.” Journal of Range Management, 78–81. [S1, S3, QC]

McArthur, E. Durant, and Bruce L Welch. 1982. “Growth Rate Differences among Big Sagebrush [Artemisia Tridentata ] Accessions and Subspecies.” Journal of Range Management 35 (3):396–401. <https://doi.org/10.2307/3898327>. [S1, QC]

McIntyre, Patrick J., and Sharon Y. Strauss. 2014. “Phenotypic and Transgenerational Plasticity Promote Local Adaptation to Sun and Shade Environments.” Evolutionary Ecology 28 (2):229–46. <https://doi.org/10.1007/s10682-013-9670-y>. [S1, S3, QC]

McMillan, Calvin. 1957. “Nature of the Plant Community. III. Flowering Behavior within Two Grassland Communities under Reciprocal Transplanting.” American Journal of Botany, 144–53. [S1, QC]

McMillan, Calvin. 1959a. “Nature of the Plant Community. V. Variation within the True Prairie Community-Type.” American Journal of Botany, 418–24. [S1, QC]

McMillan, Calvin. 1959b. “The Role of Ecotypic Variation in the Distribution of the Central Grassland of North America.” Ecological Monographs 29 (4):286–308. https://doi.org/Genetic Considerations in Ecological Restoration. [S1, S2, QC]

McNaughton, S.J. 1966. “Thermal Inactivation Properties of Enzymes from Typha Latifolia L. Ecotypes.” Plant Physiology 41:1736–38. <https://doi.org/10.2307/4260909>. [S1]

Mealor, Brian A., and Ann L. Hild. 2007. “Post-Invasion Evolution of Native Plant Populations: A Test of Biological Resilience.” Oikos 116 (9):1493–1500. <https://doi.org/10.1111/j.2007.0030-1299.15781.x>. [S1, S2]

Messina, Frank J., Susan L. Durham, James H. Richards, and E. Durant McArthur. 2002. “Trade-off between Plant Growth and Defense? A Comparison of Sagebrush Populations.” Oecologia 131 (1):43–51. <https://doi.org/10.1007/s00442-001-0859-3>. [S1, QC]

Meyer, S. E., and S. G. Kitchen. 1994. “Life History Variation in Blue Flax (Linum Perenne: Linaceae): Seed Germination Phenology.” American Journal of Botany 81 (5):528–35. <https://doi.org/10.2307/2445726>. [S1, S2, QC]

Meyer, S. E., S. G. Kitchen, and S. L. Carlson. 1995. “Seed Germination Timing Patterns in Intermountain Penstemon (Scrophulariaceae).” American Journal of Botany. <https://doi.org/10.2307/2445584>. [S1, S2]

Meyer, S. E., E. D. McArthur, and G. L. Jorgensen. 1989. “Variation in Germination Response to Temperature in Rubber Rabbitbrush (Chrysothamnus Nauseosus: Asteraceae) and Its Ecological Implications.” American Journal of Botany. <https://doi.org/10.2307/2444519>. [S1]

Meyer, S. E., and S. B. Monsen. 1992. “Big Sagebrush Germination Patterns: Subspecies and Population Differences.” Journal of Range Management 45 (1):87–93. <https://doi.org/10.2307/4002533>. [S1, S2, QC]

Meyer, SE, J Beckstead, PS Allen, and H Pullman. 1995. “Germination Ecophysiology of Leymus Cinereus (Poaceae).” International Journal of Plant Sciences 156 (2):206–15. <https://doi.org/10.1086/297242>. [S1, S2, QC]

Meyer, Susan E. 1992. “Habitat Correlated Variation in Firecracker Penstemon (Penstemon Eatonii Gray: Scrophulariaceae) Seed Germination Response.” Bulletin of the Torrey Botanical Club 119 (3):268–79. <https://doi.org/10.2307/2996758>. [S1, S2]

Meyer, Susan E. 1997. “Ecological Correlates of Achene Mass Variation in Chrysothamnus Nauseosus (Asteraceae).” American Journal of Botany 84 (4):471–77. <https://doi.org/10.2307/2446023>. [S1, S2, QC]

Meyer, Susan E., and Stephanie L. Carlson. n.d. “Seed Germination Biology of Intermountain Populations of Fourwing Saltbush (Atriplex Canescens: Chenopodiaceae).” In: Sosebee, Ronald E.; Wester, David B.; Britton, Carlton M.; McArthur, E. Durant; Kitchen, Stanley G., Comps. 2007. Proceedings: Shrubland Dynamics—fire and Water; 2004 August 10-12; Lubbock, TX. Proceedings RMRS-P-47. Fort Collins, CO: U.S.D.A. [S1, S2]

Meyer, Susan E., Stephanie L. Carlson, and Susan C. Garvin. 1998. “Seed Germination Regulation and Field Seed Bank Carryover in Shadscale (Atriplex Confertifolia: Chenopodiaceae).” Journal of Arid Environments 38 (2):255–67. <https://doi.org/10.1006/jare.1997.0321>. [S1, S2, QC]

Meyer, Susan E., and Stephen B. Monsen. 1991. “Habitat-Correlated Variation in Mountain Big Sagebrush (Artemisia Tridentata Ssp. Vaseyana) Seed Germination Patterns.” Ecology 72 (2):739–42. [S1, S2]

Meyer, Susan E., Stephen B. Monsen, and E. Durant McArthur. 1990. “Germination Response of Artemisia Tridentata (Asteraceae) to Light and Chill: Patterns of Between-Population Variation.” Botanical Gazette. <https://doi.org/10.1086/337817>. [S1, S2]

Meyer, Susan E, and Stanley G Kitchen. 1994. “Habitat-Correlated Variation in Seed Germination Response to Chilling in Penstemon Section Glabri (Scrophulariaceae).” American Midland Naturalist 132 (2):349–65. <https://doi.org/10.2307/2426591>. [S1]

Miller, Roy V. 1967. “Ecotypic Variation in Andropogon Scoparius and Bouteloua Gracilis.” Ph.D. Dissertation, Colorado State University, Fort Collins. [S1, S2]

Miller, Stephanie A., Amy Bartow, Melanie Gisler, Kimiora Ward, Amy S. Young, and Thomas N. Kaye. 2011. “Can an Ecoregion Serve as a Seed Transfer Zone? Evidence from a Common Garden Study with Five Native Species.” Restoration Ecology 19 (201):268–76. <https://doi.org/10.1111/j.1526-100X.2010.00702.x>. [S1, S2, QC]

Monaco, T. A., S. B. Monsen, B. N. Smith, and L. D. Hansen. 2005. “Temperature-Dependent Physiology of Poa Secunda, a Cool Season Grass Native to the Great Basin, United States.” Russian Journal of Plant Physiology 52 (5):653–58. <https://doi.org/10.1007/s11183-005-0096-4>. [S1]

Monson, R K, S D Smith, J L Gehring, W D Bowman, S R Szarek, British Ecological Society, and Functional Ecology. 1992. “Physiological Differentiation within an Encelia Farinosa Population along a Short Topographic Gradient in the Sonoran Desert.” Functional Ecology 6 (6):751–59. [S1, S2]

Moyer, J.L., and R. L. Lang. 1976. “Variable Germination Response to Temperature for Different Sources of Winterfat Seed.” Journal of Range Management 29:320–21. [S1]

Mummey, Daniel L., M.E. Herget, K.M. Hufford, and L. Shreading. 2016. “Germination Timing and Seedling Growth of Poa Secunda and the Invasive Grass, Bromus Tectorum, in Response to Temperature: Evaluating Biotypes for Seedling Traits That Improve Establishment.” Ecological Restoration 34 (3):200–208. [S1, S2]

Munda, B.D., S.M. Lambert, and J.C. Garrison. 1990. “Registration of ‘Santa Rita’ Fourwing Saltbush.” Crop Science 30 (6). [S1]

Nasri, Mohamed, and Paul S. Doescher. 1995. “Effect of Temperature on Growth of Cheatgrass and Idaho Fescue.” Journal of Range Management, 406–9. [S1]

Orodho, A B, R L Cuany, and M J Trlica. 1998. “Previous Grazing or Clipping Affects Seed of Indian Ricegrass.” Journal of Range Management 51 (1):37–41. <https://doi.org/none>. [S1, S2]

Orodho, Apollo B., and M J Trlica. 1990. “Clipping and Long-Term Grazing Effects on Biomass and Carbohydrate Reserves of Indian Ricegrass.” Journal of Range Management, 52–57. [S1, S2]

Par, Steve, and Marti Walsh. 2008. Notice of Release of Long Ridge Germplasm Utah Serviceberry. Upper Colorado Environmental Plant Center, USDA NRCS, Colorado State Agricultural Experiment Station. https://www.nrcs.usda.gov/Internet/FSE_PLANTMATERIALS/publications/copmcrn8043.pdf [S1]

Parsons, Matthew C., Thomas A. Jones, Steven R. Larson, Ivan W. Mott, and Thomas A. Monaco. 2011. “Ecotypic Variation in Elymus Elymoides Subsp. Brevifolius in the Northern Intermountain West.” Rangeland Ecology and Management 64 (6):649–58. <https://doi.org/10.2111/REM-D-09-00143.1>. [S1]

Parsons, Matthew C., Thomas A. Jones, and Thomas A. Monaco. 2011. “Genetic Variation for Adaptive Traits in Bottlebrush Squirreltail in the Northern Intermountain West, United States.” Restoration Ecology 19 (4):460–69. <https://doi.org/10.1111/j.1526-100X.2010.00705.x>. [S1, S2]

Pearcy, R W, and R T Ward. 1972. “Phenology and Growth of Rocky Mountain Populations of Deschampsia Caespitosa at Three Elevations in Colorado.” Ecology 53:1171–78. <https://doi.org/10.2307/1935431>. [S1, S2, S3, QC]

Pendleton, B. K., and S. E. Meyer. 2004. “Habitat-Correlated Variation in Blackbrush (Coleogyne Ramosissima: Rosaceae) Seed Germination Response.” Journal of Arid Environments 59 (2):229–43. <https://doi.org/10.1016/j.jaridenv.2003.12.009>. [S1, S2]

Petersen, J L, D N Ueckert, R L Potter, and J E Huston. 1987. “Ecotypic Variation in Selected Fourwing Saltbush Populations in Western Texas USA.” Journal of Range Management 40 (4):361–66. https://doi.org/Genetic Considerations in Ecological Restoration. [S1, S3, QC]

Phillips, Nathan C., Daniel T. Drost, William A. Varga, Leila M. Shultz, and Susan E. Meyer. 2010. “Germination Characteristics along Altitudinal Gradients in Three Intermountain Allium spp.(Amaryllidaceae).” Seed Technology 32 (1):15–25. <http://www.jstor.org/stable/23433619>. [S1, S2]

Polley, H. W., and J. K. Detling. 1988. “Herbivory Tolerance of Agropyron Smithii Populations with Different Grazing Histories.” Oecologia 77 (2):261–67. <https://doi.org/10.1007/BF00379196>. [S1, S2]

Potter, R.L., D.N. Ueckert, and J L Petersen. 1986. “Germination of Fourwing Saltbush Seeds: Interaction of Temperature, Osmotic Potential, and pH.” Journal of Range Management, 43–46. [S1]

Quinn, James A., and Richard T Ward. 1969. “Ecological Differentiation in Sand Dropseed (Sporobolus Cryptandrus).” Ecological Monographs 39 (1):61–78. [S1, S2, QC]

Quinn, James A, and Jeffrey D Wetherington. 2002. “Genetic Variability and Phenotypic Plasticity in Flowering Phenology in Populations of Two Grasses.” Journal of the Torrey Botanical Society 129 (2):96–106. <https://doi.org/10.2307/3088723>. [S1, S2]

Ray, Peter M., and William E. Alexander. 1966. “Photoperiodic Adaptation to Latitude in Xanthium Strumarium.” American Journal of Botany, 806–16. [S1, S2, QC]

Rice, Kevin J., and Eric E. Knapp. 2008. “Effects of Competition and Life History Stage on the Expression of Local Adaptation in Two Native Bunchgrasses.” Restoration Ecology 16 (1):12–23. <https://doi.org/10.1111/j.1526-100X.2007.00257.x>. [S1, S3, QC]

Richardson, Bryce A., Stanley G. Kitchen, Rosemary L. Pendleton, Burton K. Pendleton, Matthew J. Germino, Gerald E. Rehfeldt, and Susan E. Meyer. 2014. “Adaptive Responses Reveal Contemporary and Future Ecotypes in a Desert Shrub.” Ecological Applications 24 (2):413–27. <https://doi.org/10.1890/13-0587.1>. [S1, S2]

Richardson, Bryce A., Hector G. Ortiz, Stephanie L. Carlson, Deidre M. Jaeger, Nancy L. Shaw, and D. P.C. Peters. 2015. “Genetic and Environmental Effects on Seed Weight in Subspecies of Big Sagebrush: Applications for Restoration.” Ecosphere 6 (10). <https://doi.org/10.1890/ES15-00249.1>. [S1, QC]

Robertson, Phillip A. 1976. “Photosynthetic and Respiratory Responses of Natural Populations of Koeleria Cristata Grown in Three Environmental Regimes.” Botanical Gazette 137 (1):94–98. [S1]

Robertson, Phillip A., and Richard T Ward. 1970. “Ecotypic Differentiation in Koeleria Cristata (L.) Pers. from Colorado and Related Area.” Ecology 51 (6):1083–87. [S1, S2, QC]

Rogler, George A. 1960. “Relation of Seed Dormancy of Indian Ricegrass (Oryzopsis Hymenoides (Roem. & Schult) Ricker.) to Age and Treatment 1.” Agronomy Journal 52 (8):470–73. [S1]

Rowe, Courtney L J, and Elizabeth A. Leger. 2011. “Competitive Seedlings and Inherited Traits: A Test of Rapid Evolution of Elymus Multisetus (Big Squirreltail) in Response to Cheatgrass Invasion.” Evolutionary Applications 4 (3):485–98. <https://doi.org/10.1111/j.1752-4571.2010.00162.x>. [S1, S2]

Rowe, Courtney L. J., and Elizabeth A. Leger. 2012. “Seed Source Affects Establishment of Elymus Multisetus in Postfire Revegetation in the Great Basin.” Western North American Naturalist 72 (4):543–53. <https://doi.org/10.3398/064.072.0410>. [S1, S3]

Rumbaugh, M.D., and B.M. Pendery. 1993. “Registration of ARS-2892 Munroe Globemallow Germplasm.” Crop Science 33 (5). [S1]

Rumbaugh, M.D., B.M. Pendery, H.F. Mayland, and G.E. Shewmaker. 1993. “Registration of ARS-2936 Scarlet Globemallow Germplasm.” Crop Science 33:1106–8. [S1]

Sanderson, S. C., H. C. Stutz, and E. D. McArthur. 1990. “Geographic Differentiation in Atriplex Confertifolia.” American Journal of Botany. [S1, S2]

Schellenberg, M.P. 2003. “Germination Temperature Response of Two Ecotypes of Winterfat [Kraschenninikovia Lanata (Pursh) Guldenstaedt].” Canadian Journal of Plant Science 83 (1):65–68. [S1]

Shaw, Nancy L., Marshall R. Haferkamp, and Emerenciana G. Hurd. 1994. “Germination and Seedling Establishment of Spiny Hopsage in Response to Planting Date and Seedbed Environment.” Journal of Range Management 47 (2):165–74. <https://doi.org/10.2307/4002827>. [S1, QC]

Shock, Clinton C., Erik B. Feibert, A. Rivera, Lamont D. Saunders, Nancy Shaw, and Francis F. Kilkenny. 2016. “Irrigation Requirements for Seed Production of Five Lomatium Species in a Semiarid Environment.” HortScience 51 (10):1270–77. [S1]

Slauson, William L., and Richard T Ward. 1986. “Ecogenetic Patterns of Four Shrub Species in Semi-Arid Communities of Northwest Colorado.” The Southwestern Naturalist, 319–29. [S1, S2]

Smith, David Solance, Matthew K. Lau, Ryan Jacobs, Jenna A. Monroy, Stephen M. Shuster, and Thomas G. Whitham. 2015. “Rapid Plant Evolution in the Presence of an Introduced Species Alters Community Composition.” Oecologia 179 (2):563–72. <https://doi.org/10.1007/s00442-015-3362-y>. [S1, S2]

Smith, David Solance, Jennifer A. Schweitzer, Philip Turk, Joseph K. Bailey, Stephen C. Hart, Stephen M. Shuster, and Thomas G. Whitham. 2012. “Soil-Mediated Local Adaptation Alters Seedling Survival and Performance.” Plant and Soil 352 (1–2):243–51. <https://doi.org/10.1007/s11104-011-0992-7>. [S1]

Springfield, H.W. 1968. “Germination of Winterfat Seeds under Different Moisture Stresses and Temperatures.” Journal of Range Management, 314–16. [S1]

Springfield, H.W. 1966. “Germination of Fourwing Saltbush Seeds at Different Levels of Moisture Stress.” Agronomy Journal 58 (2):149–50. [S1]

Springfield, H.W. 1964. “Some Factors Affecting Germination of Fourwing Saltbush.” Rocky Mountain Forest and Range Experiment Station, Forest Service, US Department of Agriculture 25. [S1, S2]

St. Clair, John Bradley, Francis F. Kilkenny, Richard C. Johnson, Nancy L. Shaw, and George Weaver. 2013. “Genetic Variation in Adaptive Traits and Seed Transfer Zones for Pseudoroegneria Spicata (Bluebunch Wheatgrass) in the Northwestern United States.” Evolutionary Applications 6 (6):933–48. <https://doi.org/10.1111/eva.12077>. [S1, S2, QC]

Staub, Jack E, Matthew D Robbins, Yingmei Ma, and Paul G Johnson. 2014. “Phenotypic and Genotypic Analysis of a US Native Fine-Leaved Festuca Population Reveals Its Potential Use for Low-Input Urban Landscapes.” Journal of the American Society for Horticultural Science 139 (6):706–15. [S1, QC]

Stettler, Jason M., Douglas A. Johnson, B. Shaun Bushman, Kevin J. Connors, Thomas A. Jones, Jennifer W. Macadam, and David J. Hole. 2017. “Utah Lotus: North American Legume for Rangeland Revegetation in the Southern Great Basin and Colorado Plateau.” Rangeland Ecology and Management 70 (6):691–99. <https://doi.org/10.1016/j.rama.2017.06.002>. [S1, S2]

Stevens, Allan R, Val Jo Anderson, and Rachel Fugal. 2014. “Competition of Squirreltail with Cheatgrass at Three Nitrogen Levels *.” American Jourmal of Plant Sciences 5:990–96. [S1, QC]

Stinson, Kristina A. 2004. “Natural Selection Favors Rapid Reproductive Phenology in Potentilla Pulcherrima (Rosaceae) at Opposite Ends of a Subalpine Snowmelt Gradient.” American Journal of Botany 91 (4):531–39. <https://doi.org/10.3732/ajb.91.4.531>. [S1, S2]

Tilley, Derek J. 2015. “Notice of Release of Amethyst Germplasm Hoary Tansyaster: Selected Class of Natural Germplasm.” Native Plants Journal 16 (1):54–60. [S1, QC]

Tilley, Derek J. 2015. Douglas’ Dustymaiden Initial Evaluation Planting Final Study Report. USDA NRCS Aberdeen Plant Materials Center. https://www.nrcs.usda.gov/Internet/FSE_PLANTMATERIALS/publications/idpmcsr13140.pdf. [S1, S2, QC]

Tisdale, E. W. 1961. “Intraspecific Variation in Festuca.” Carnegie Institution of Washington Yearbook 60:388–91. [S1]

Toole, Vivian K. 1941. “Factors Affecting the Germination of Various Dropseed Grasses (Sporobolus Spp.).” Journal of Agricultural Research 62:691–715. [S1, S2]

USDA NRCS Bridger Plant Materials Center. 1996. Notice of Release of ‘Rimrock’ Indian Ricegrass. USDA NRCS Bridger Plant Materials Center, Montana Agricultural Experiment Station, Wyoming Agricultural Experiment Station, USDA Agricultural Research Station. [S1]

USDA NRCS Los Lunas Plant Materials Center. 1973. Supporting Data For the Release of Arriba Western Wheatgrass. New Mexico State University’s Los Lunas Agricultural Science Center, Colorado State University, New Mexico Department of Transportation, and the USDA Natural Resources Conservation Service Los Lunas Plant Materials Center. [S1, QC]

Waldron, B L, K B Jensen, A J Palazzo, T J Cary, J G Robins, M D Peel, D G Ogle, and L St John. 2011. “‘Recovery’, a New Western Wheatgrass Cultivar with Improved Seedling Establishment on Rangelands.” Journal of Plant Registrations 5 (3):367–73. <https://doi.org/Doi>10.3198/Jpr2010.09.0527crc. [S1]

Wan, Changgui, Ronald E. Sosebee, and Bobby L. McMichael. 1998. “Water Relations and Root Growth of Two Populations of Gutierrezia Sarothrae.” Environmental and Experimental Botany 39 (1):11–20. <https://doi.org/10.1016/S0098-8472(97)00021-X>. [S1]

Wan, Changgui, Ronald E. Sosebee, and Bobby L. McMichael. 1995. “Water Acquisition and Rooting Characteristics in Northern and Southern Populations of Gutierrezia Sarothrae.” Environmental and Experimental Botany 35 (1):1–7. <https://doi.org/10.1016/0098-8472(94)00038-7>. [S1, S2]

Wan, Changgui, Ronald E. Sosebee, and Bobby L. McMichael. 1996. “Lateral Root Development and Hydraulic Conductance in Four Populations of Gutierrezia Sarothrae.” Environmental and Experimental Botany 36 (2):157–65. <https://doi.org/10.1016/0098-8472(96)01008-8>. [S1]

Ward, Richard T. 1969. “Ecotypic Variation in Deschampsia Caespitosa (L.) Beauv. from Colorado.” Ecology 50 (3):519. <https://doi.org/10.2307/1933914>. [S1, S2, QC]

Waser, Nickolas M., and Mary V. Price. 1985. “Reciprocal Transplant Experiments with Delphinium Nelsonii (Ranunculaceae): Evidence for Local Adaptation.” American Journal of Botany, 1726–32. [S1, S2, S3]

Wood, M. Karl, Robert W. Knight, and James A. Young. 1976. “Spiny Hopsage Germination.” Journal of Range Management, 53–56. [S1]

Workman, John P, and Neil E West. 1967. “Germination of Eurotia Lanata in Relation to Temperature and Salinity.” Ecology 48 (4):659–61. [S1]

Workman, John P, and Neil E West. 1969. “Ecotypic Variation of Eurotia Lanata Populations in Utah.” Botanical Gazette 130 (1):26–35. <https://doi.org/10.2307/2473599>. [S1, S2]

Young, J A, C D Clements, and T Jones. 2003. “Germination of Seeds of Big and Bottlebrush Squirreltail.” Journal of Range Management 56 (3):277–81. <https://doi.org/10.2307/4003819>. [S1]

Young, JA, RA Evans, and DE Palmquist. 1989. “Big Sagebrush (Artemisia Tridentata) Seed Production.” Weed Science 37 (1):47–53. http://www.jstor.org/stable/10.2307/4044754. [S1, S3, QC]

Young, James A., and Raymond A. Evans. 1989. “Reciprocal Common Garden Studies of the Germination of Seeds of Big Sagebrush (Artemisia Tridentata).” Weed Science 37 (3):319–325. <https://doi.org/none>. [S1, QC]

Young, James A, Raymond A Evans, and B L Kay. 1984. “Persistence and Colonizing Ability of Rabbitbrush Collections in a Common Garden.” Journal of Range Management 37 (4):373–77. <https://doi.org/10.2307/3898715>. [S1, QC]

Zhang, Huarong, Laura E. DeWald, Thomas E. Kolb, and Dan E. Koepke. 2011. “Genetic Variation in Ecophysiological and Survival Responses to Drought in Two Grasses: Koeleria Macrantha and Elymus Elymoides.” Western North American Naturalist 71 (1):25–32. <https://doi.org/10.3398/064.071.0104>. [S1, S2]
