## appendix 3 figures for "Strong Patterns of Intraspecific Variation and Local Adaptation in Great Basin Plants Revealed Through a Review of 75 Years of Experiments" (from another version of this preprint: version 1, 2018-12-26)

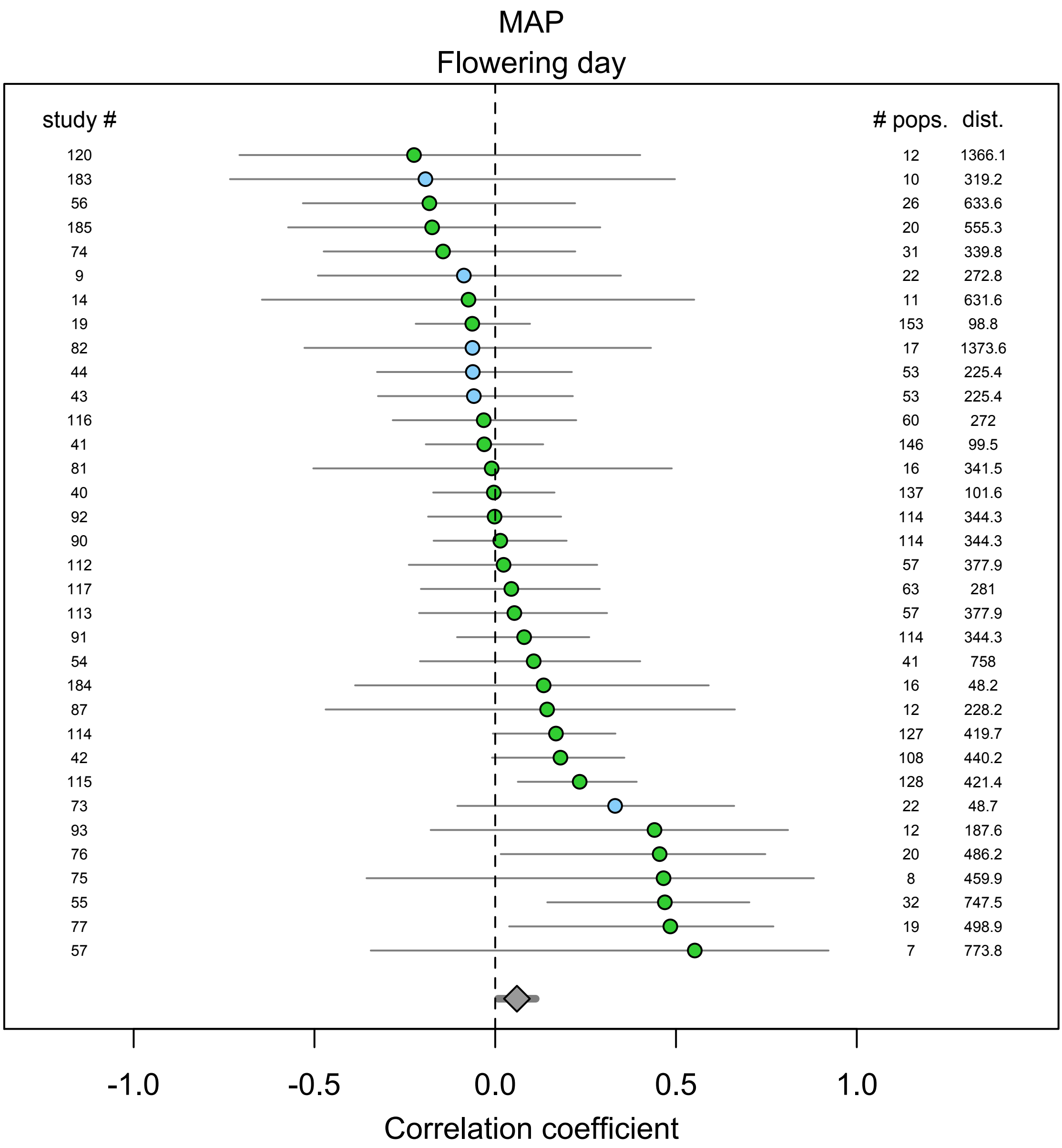
MAP
Flowering day
study #
### pops.
dist.
120 12 1366.1
183 10 319.2
56 26 633.6
185 20 555.3
74 31 339.8
9 22 272.8
14 11 631.6
19 153 98.8
82 17 1373.6
44 53 225.4
43 53 225.4
116 60 272
41 146 99.5
81 16 341.5
40 137 101.6
92 114 344.3
90 114 344.3
112 57 377.9
117 63 281
113 57 377.9
91 114 344.3
54 41 758
184 16 48.2
87 12 228.2
114 127 419.7
42 108 440.2
115 128 421.4
73 22 48.7
93 12 187.6
76 20 486.2
75 8 459.9
55 32 747.5
77 19 498.9
57 7 773.8
-1.0
-0.5
0.0
0.5
1.0
Correlation coefficient

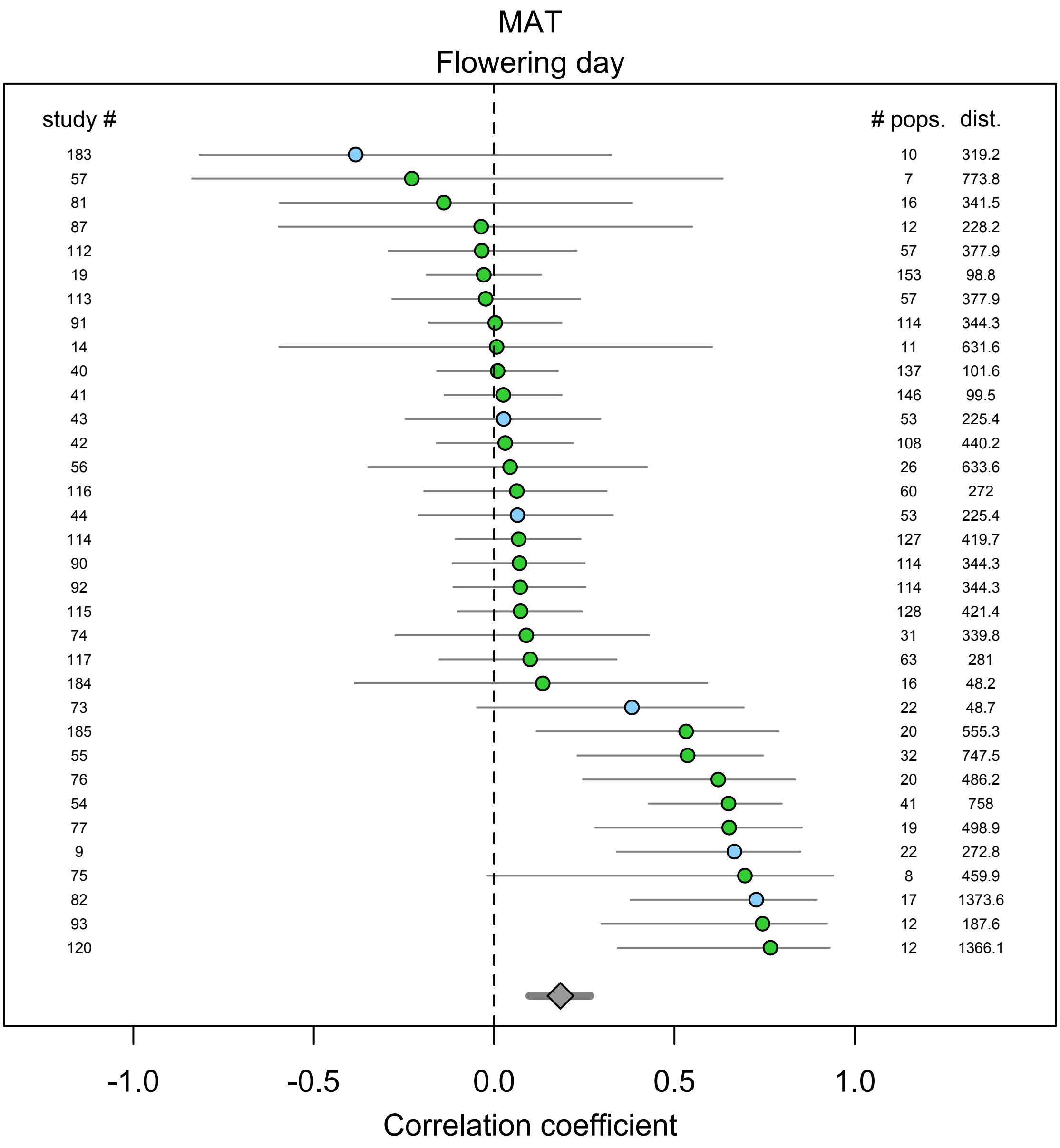
MAT
Flowering day
study #
### pops.
dist.
183 10 319.2
57 7 773.8
81 16 341.5
87 12 228.2
112 57 377.9
19 153 98.8
113 57 377.9
91 114 344.3
14 11 631.6
40 137 101.6
41 146 99.5
43 53 225.4
42 108 440.2
56 26 633.6
116 60 272
44 53 225.4
114 127 419.7
90 114 344.3
92 114 344.3
115 128 421.4
74 31 339.8
117 63 281
184 16 48.2
73 22 48.7
185 20 555.3
55 32 747.5
76 20 486.2
54 41 758
77 19 498.9
9 22 272.8
75 8 459.9
82 17 1373.6
93 12 187.6
120 12 1366.1
-1.0
-0.5
0.0
0.5
1.0
Correlation coefficient

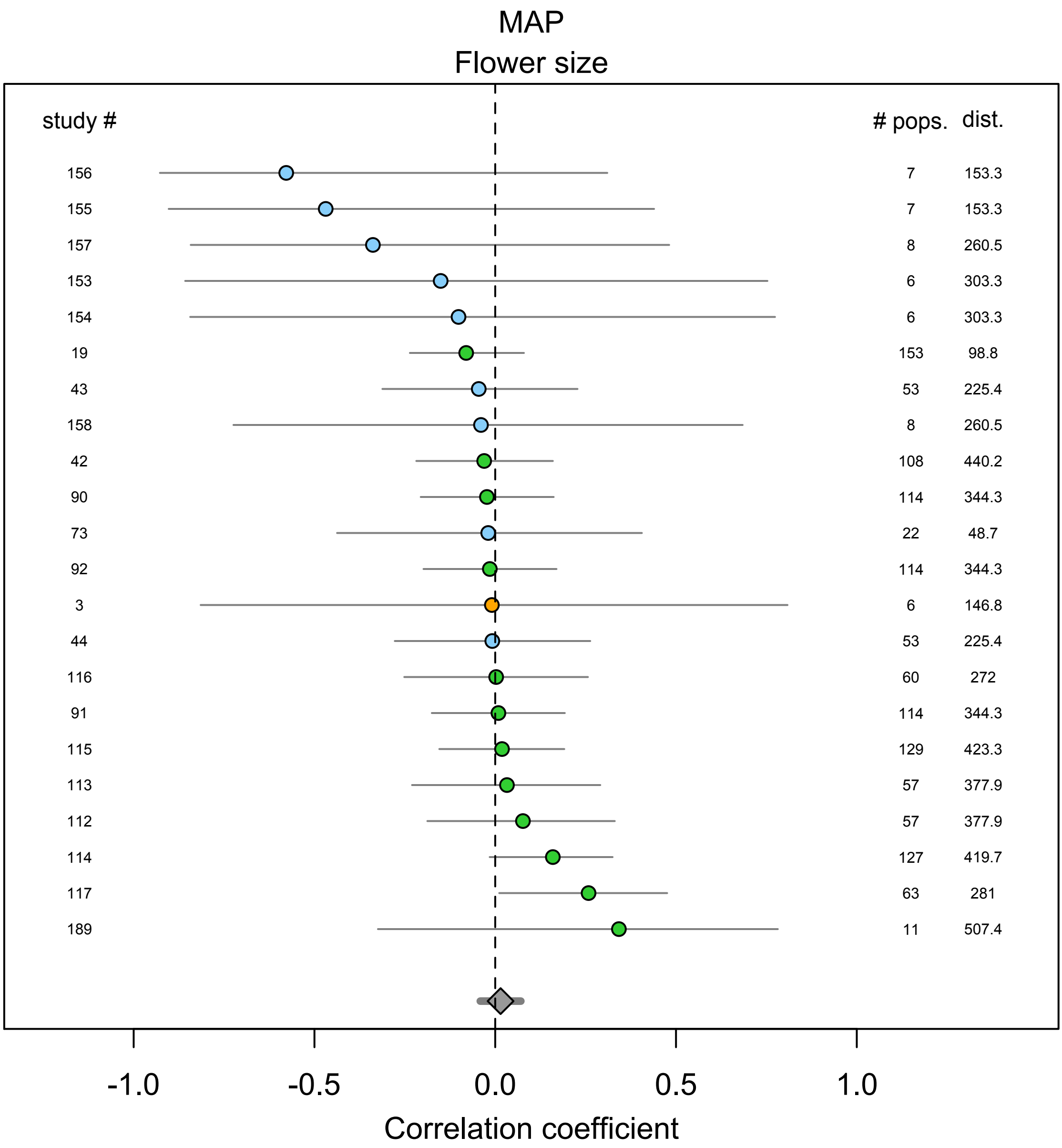
MAP
Flower size
study #
### pops.
dist.
156 7 153.3
155 7 153.3
157 8 260.5
153 6 303.3
154 6 303.3
19 153 98.8
43 53 225.4
158 8 260.5
42 108 440.2
90 114 344.3
73 22 48.7
92 114 344.3
3 6 146.8
44 53 225.4
116 60 272
91 114 344.3
115 129 423.3
113 57 377.9
112 57 377.9
114 127 419.7
117 63 281
189 11 507.4
-1.0
-0.5
0.0
0.5
1.0
Correlation coefficient

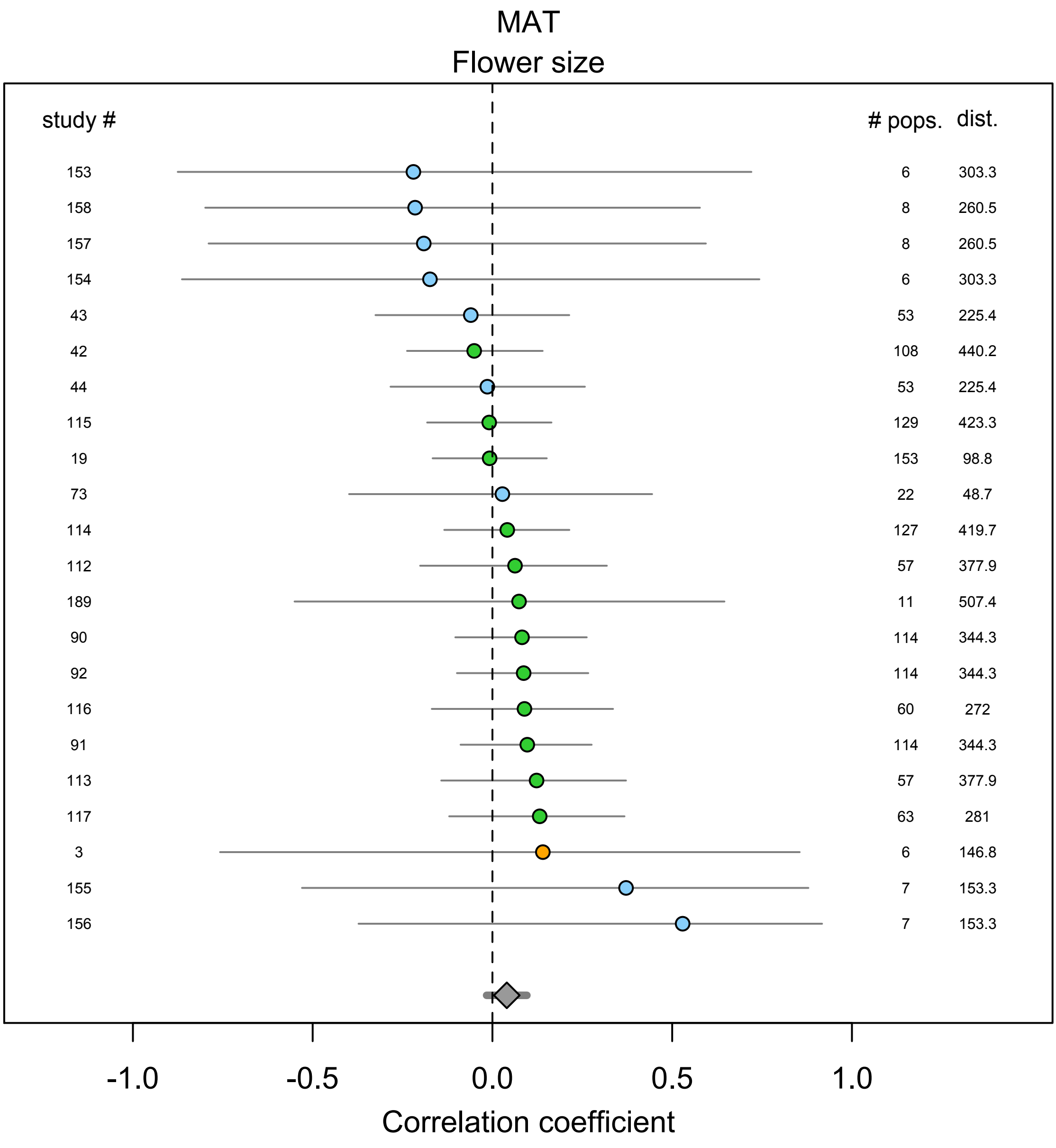
MAT
Flower size
study #
### pops.
dist.
153 6 303.3
158 8 260.5
157 8 260.5
154 6 303.3
43 53 225.4
42 108 440.2
44 53 225.4
115 129 423.3
19 153 98.8
73 22 48.7
114 127 419.7
112 57 377.9
189 11 507.4
90 114 344.3
92 114 344.3
116 60 272
91 114 344.3
113 57 377.9
117 63 281
3 6 146.8
155 7 153.3
156 7 153.3
-1.0
-0.5
0.0
0.5
1.0
Correlation coefficient

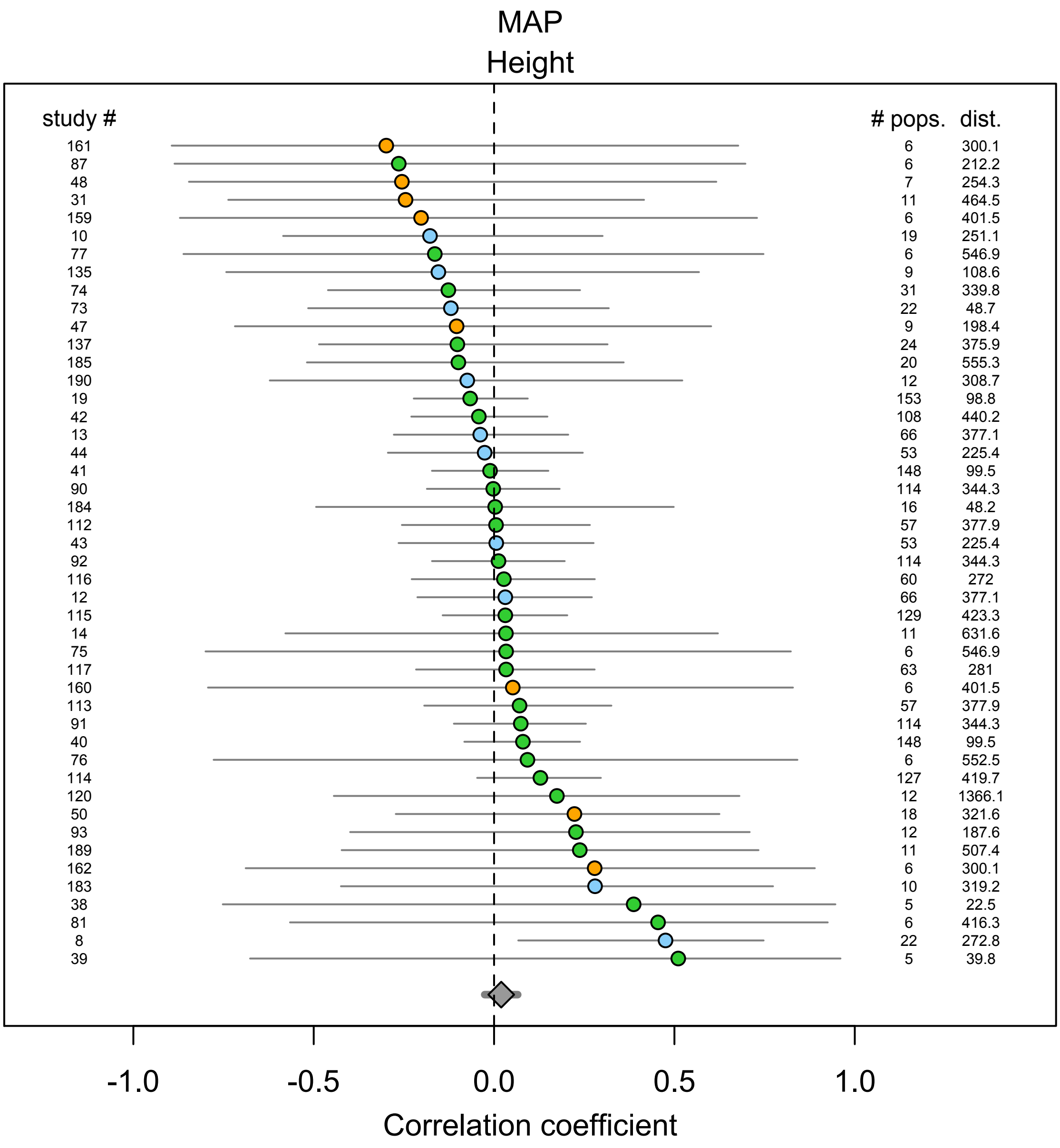
MAP
Height
study #
### pops.
dist.
161 6 300.1
87 6 212.2
48 7 254.3
31 11 464.5
159 6 401.5
10 19 251.1
77 6 546.9
135 9 108.6
74 31 339.8
73 22 48.7
47 9 198.4
137 24 375.9
185 20 555.3
190 12 308.7
19 153 98.8
42 108 440.2
13 66 377.1
44 53 225.4
41 148 99.5
90 114 344.3
184 16 48.2
112 57 377.9
43 53 225.4
92 114 344.3
116 60 272
12 66 377.1
115 129 423.3
14 11 631.6
75 6 546.9
117 63 281
160 6 401.5
113 57 377.9
91 114 344.3
40 148 99.5
76 6 552.5
114 127 419.7
120 12 1366.1
50 18 321.6
93 12 187.6
189 11 507.4
162 6 300.1
183 10 319.2
38 5 22.5
81 6 416.3
8 22 272.8
39 5 39.8
-1.0
-0.5
0.0
0.5
1.0
Correlation coefficient

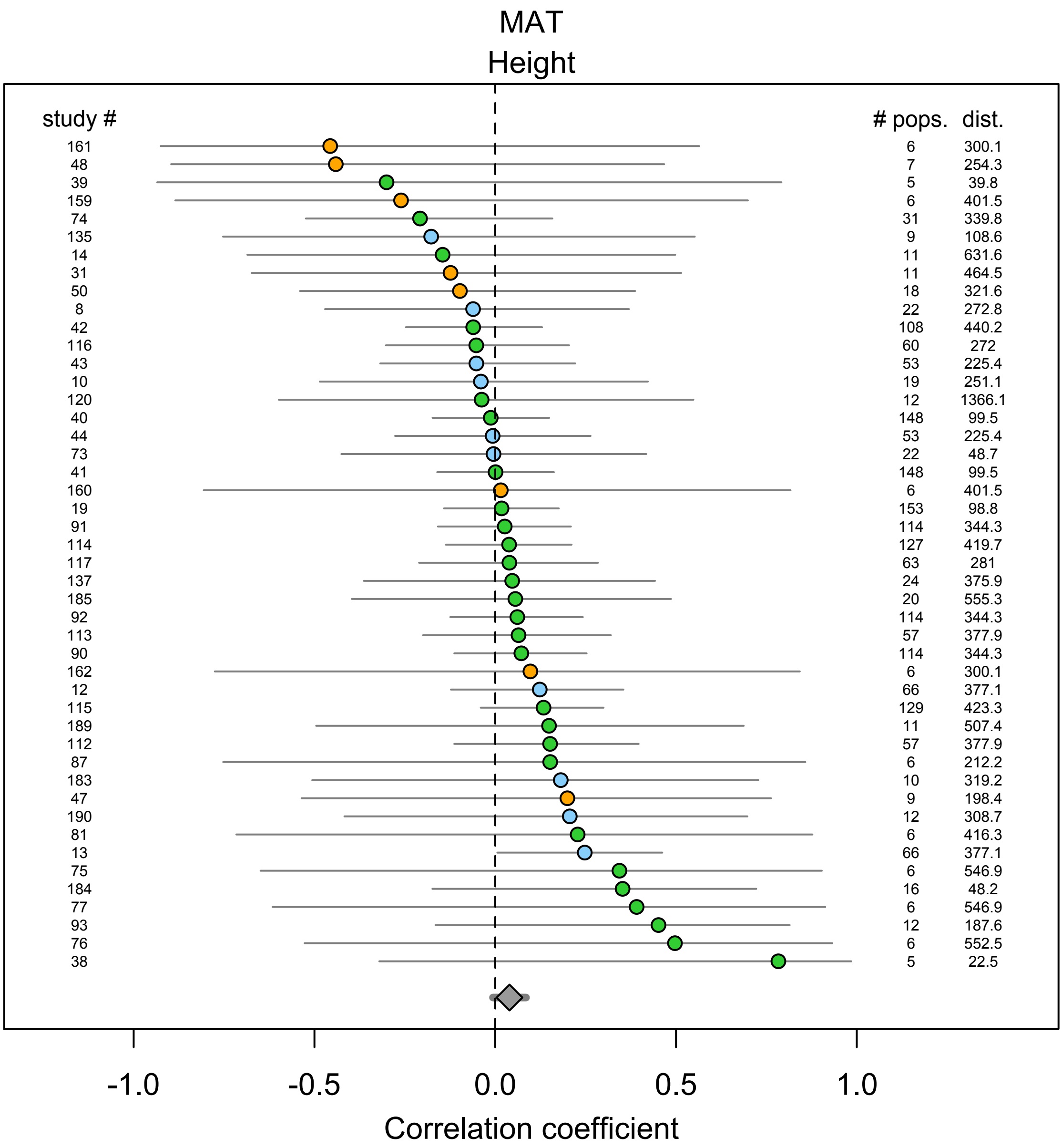
MAT
Height
study #
### pops.
dist.
161 6 300.1
48 7 254.3
39 5 39.8
159 6 401.5
74 31 339.8
135 9 108.6
14 11 631.6
31 11 464.5
50 18 321.6
8 22 272.8
42 108 440.2
116 60 272
43 53 225.4
10 19 251.1
120 12 1366.1
40 148 99.5
44 53 225.4
73 22 48.7
41 148 99.5
160 6 401.5
19 153 98.8
91 114 344.3
114 127 419.7
117 63 281
137 24 375.9
185 20 555.3
92 114 344.3
113 57 377.9
90 114 344.3
162 6 300.1
12 66 377.1
115 129 423.3
189 11 507.4
112 57 377.9
87 6 212.2
183 10 319.2
47 9 198.4
190 12 308.7
81 6 416.3
13 66 377.1
75 6 546.9
184 16 48.2
77 6 546.9
93 12 187.6
76 6 552.5
38 5 22.5
-1.0
-0.5
0.0
0.5
1.0
Correlation coefficient

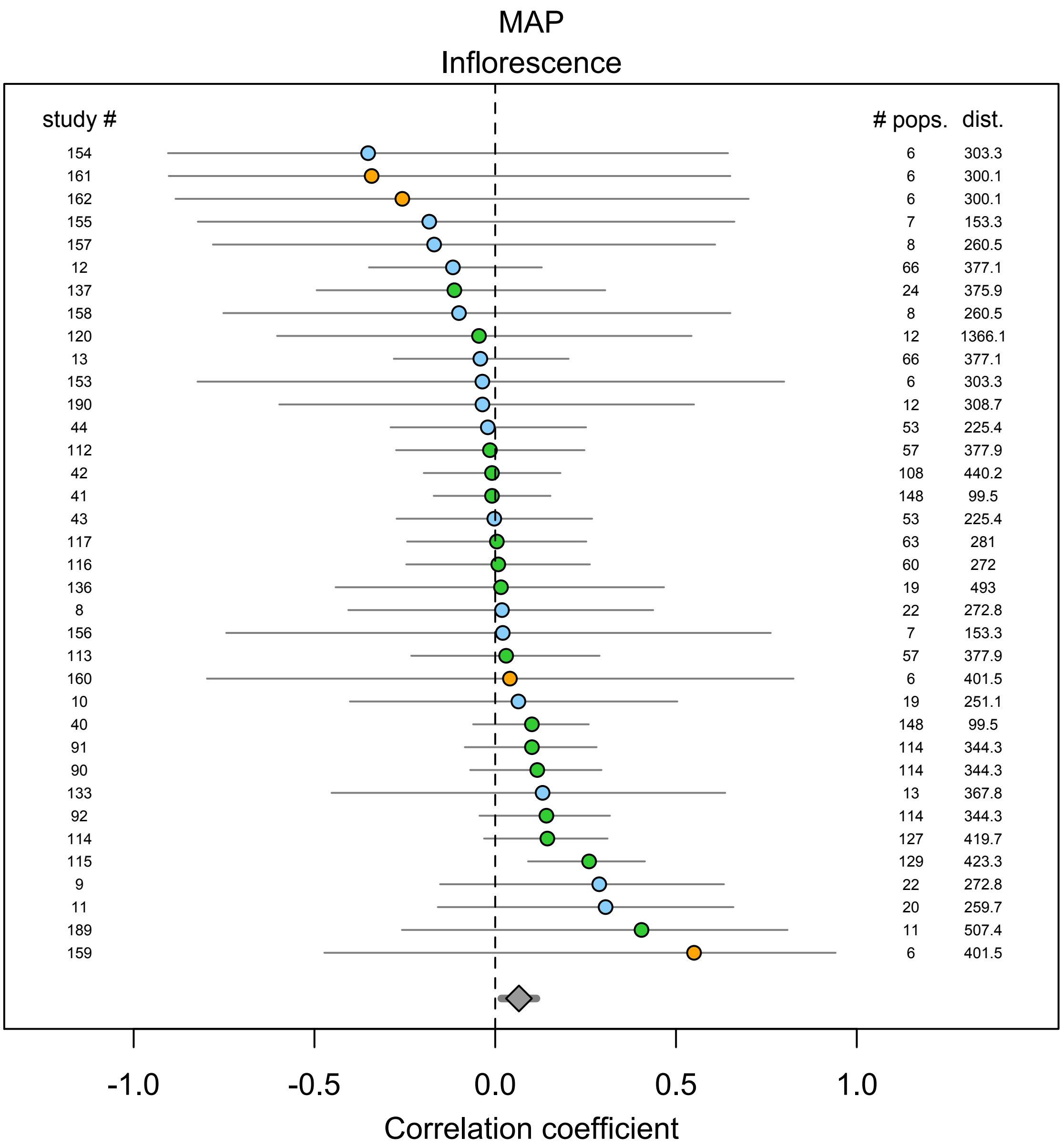
MAP
Inflorescence
study #
### pops.
dist.
154
161
162
155
157
12
137
158
120
13
153
190
44
112
42
41
43
117
116
136
8
156
113
160
10
40
91
90
133
92
114
115
9
11
189
159
6 303.3
6 300.1
6 300.1
7 153.3
8 260.5
66 377.1
24 375.9
8 260.5
12 1366.1
66 377.1
6 303.3
12 308.7
53 225.4
57 377.9
108 440.2
148 99.5
53 225.4
63 281
60 272
19 493
22 272.8
7 153.3
57 377.9
6 401.5
19 251.1
148 99.5
114 344.3
114 344.3
13 367.8
114 344.3
127 419.7
129 423.3
22 272.8
20 259.7
11 507.4
6 401.5
-1.0
-0.5
0.0
0.5
1.0
Correlation coefficient

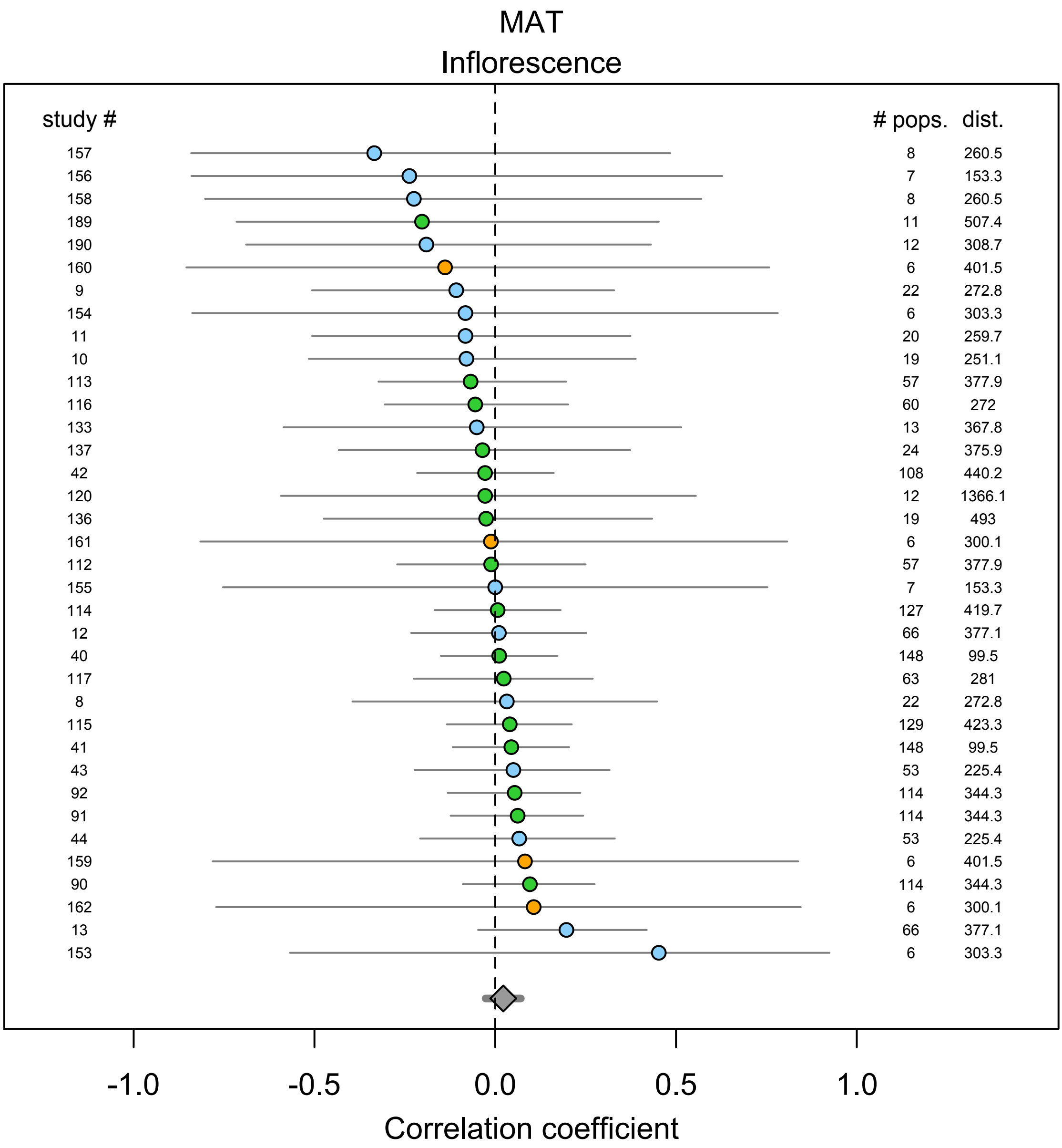

MAT
Inflorescence
study #
### pops.
dist.
157 8 260.5
156 7 153.3
158 8 260.5
189 11 507.4
190 12 308.7
160 6 401.5
9 22 272.8
154 6 303.3
11 20 259.7
10 19 251.1
113 57 377.9
116 60 272
133 13 367.8
137 24 375.9
42 108 440.2
120 12 1366.1
136 19 493
161 6 300.1
112 57 377.9
155 7 153.3
114 127 419.7
12 66 377.1
40 148 99.5
117 63 281
8 22 272.8
115 129 423.3
41 148 99.5
43 53 225.4
92 114 344.3
91 114 344.3
44 53 225.4
159 6 401.5
90 114 344.3
162 6 300.1
13 66 377.1
153 6 303.3
-1.0
-0.5
0.0
0.5
1.0
Correlation coefficient

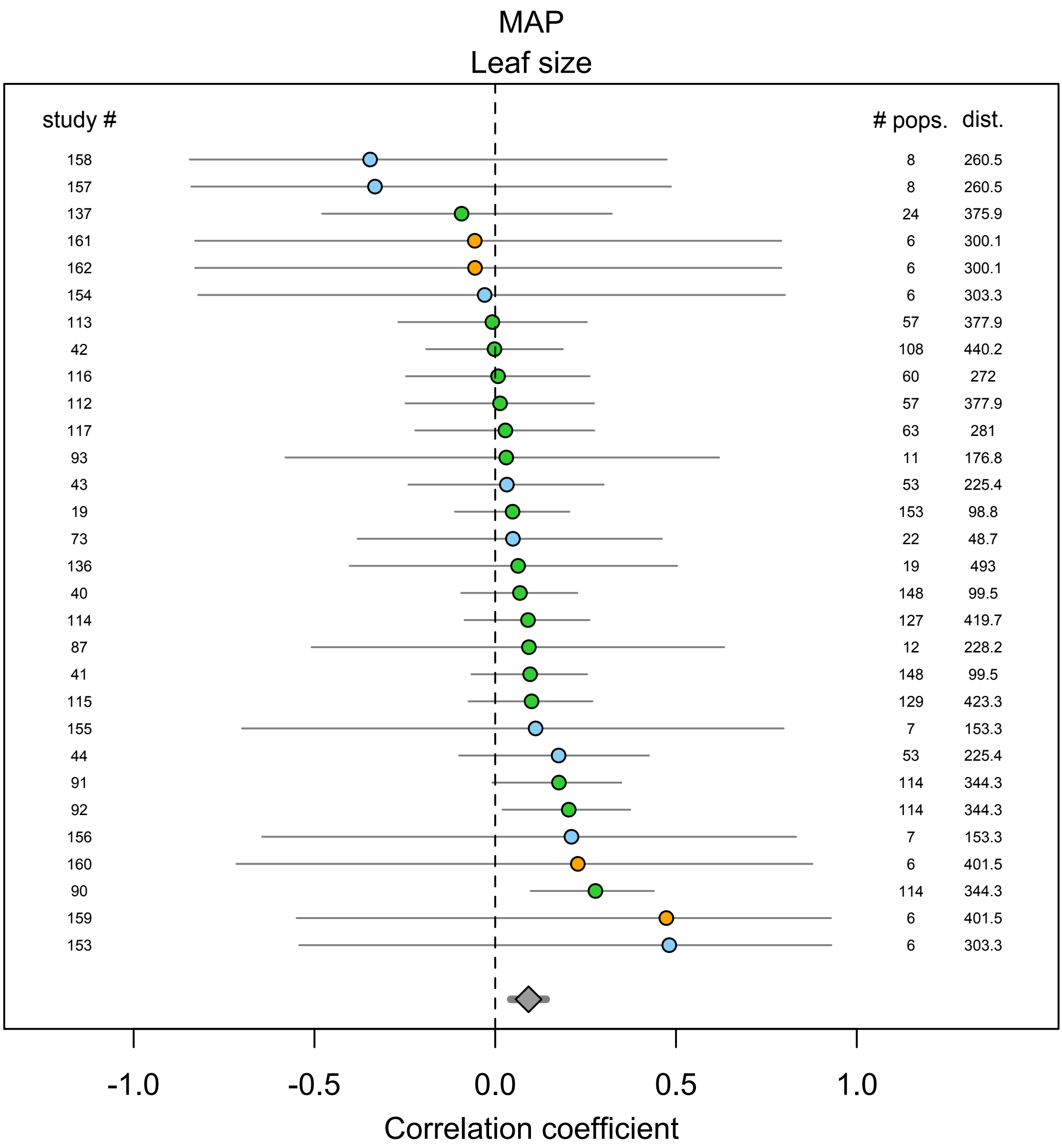
MAP
Leaf size
study #
### pops.
dist.
158 8 260.5
157 8 260.5
137 24 375.9
161 6 300.1
162 6 300.1
154 6 303.3
113 57 377.9
42 108 440.2
116 60 272
112 57 377.9
117 63 281
93 11 176.8
43 53 225.4
19 153 98.8
73 22 48.7
136 19 493
40 148 99.5
114 127 419.7
87 12 228.2
41 148 99.5
115 129 423.3
155 7 153.3
44 53 225.4
91 114 344.3
92 114 344.3
156 7 153.3
160 6 401.5
90 114 344.3
159 6 401.5
153 6 303.3
-1.0
-0.5
0.0
0.5
1.0
Correlation coefficient

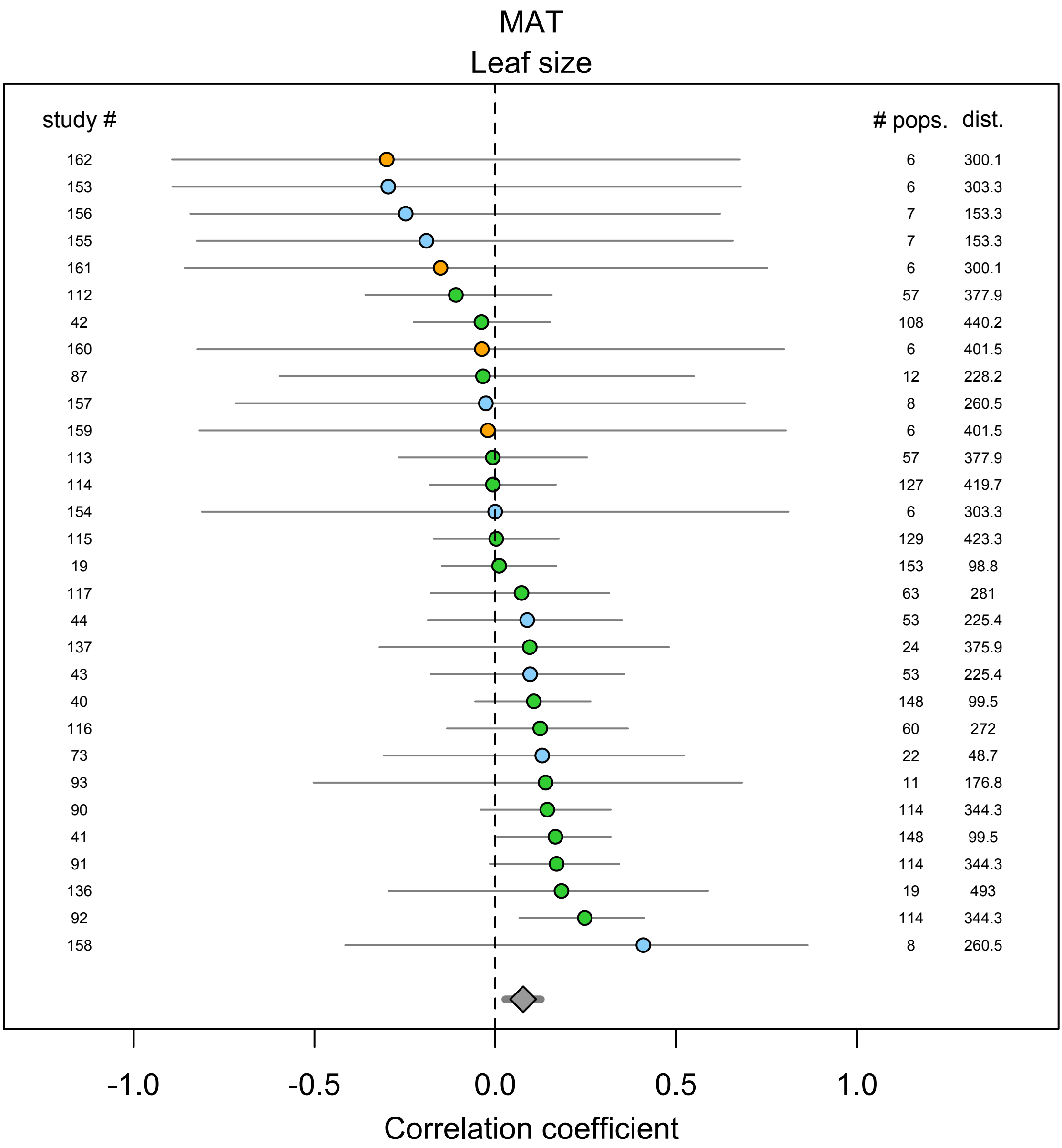
MAT
Leaf size
study #
### pops.
dist.
162 6 300.1
153 6 303.3
156 7 153.3
155 7 153.3
161 6 300.1
112 57 377.9
42 108 440.2
160 6 401.5
87 12 228.2
157 8 260.5
159 6 401.5
113 57 377.9
114 127 419.7
154 6 303.3
115 129 423.3
19 153 98.8
117 63 281
44 53 225.4
137 24 375.9
43 53 225.4
40 148 99.5
116 60 272
73 22 48.7
93 11 176.8
90 114 344.3
41 148 99.5
91 114 344.3
136 19 493
92 114 344.3
158 8 260.5
-1.0
-0.5
0.0
0.5
1.0
Correlation coefficient

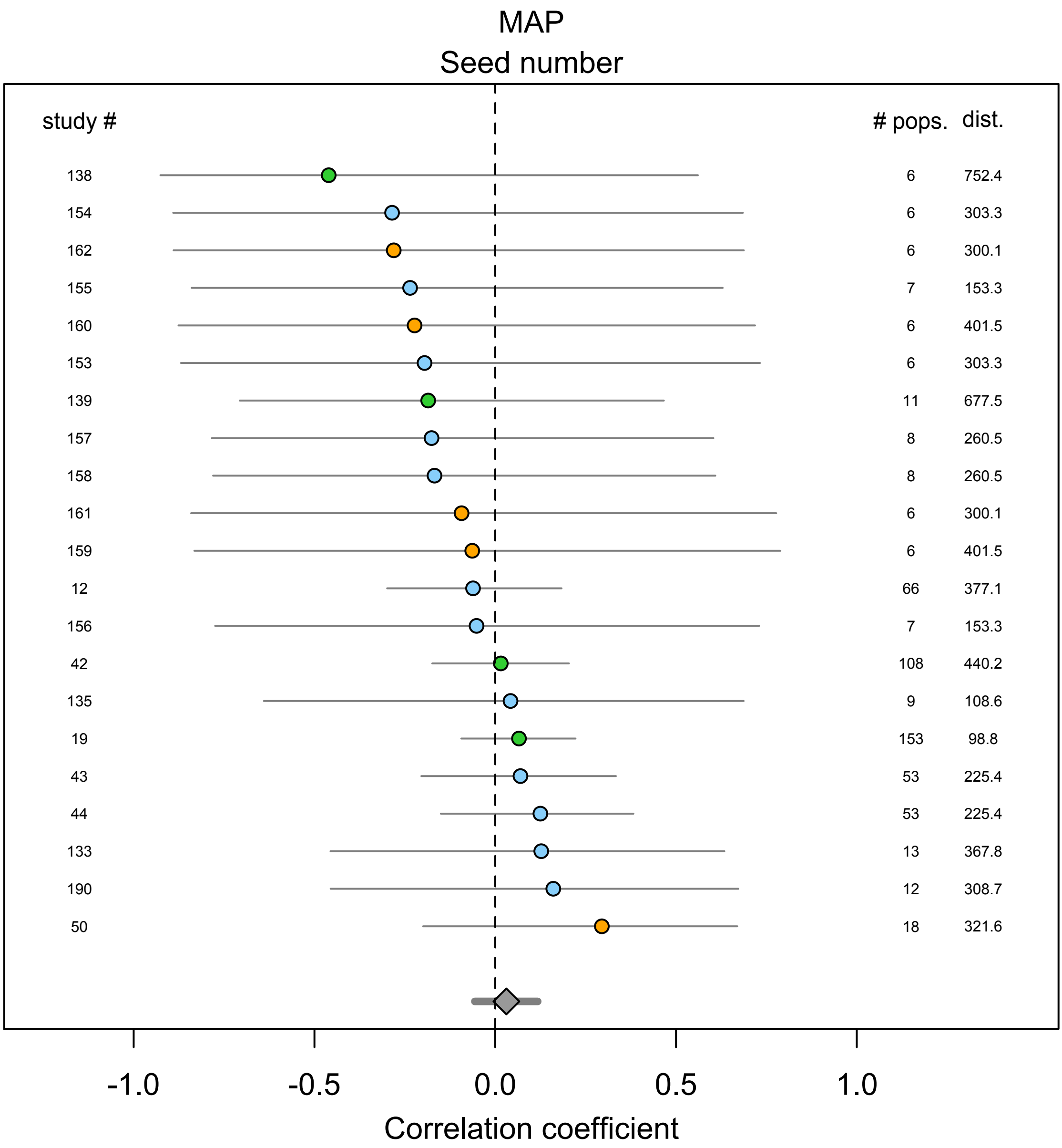
MAP
Seed number
study #
### pops.
dist.
138 6 752.4
154 6 303.3
162 6 300.1
155 7 153.3
160 6 401.5
153 6 303.3
139 11 677.5
157 8 260.5
158 8 260.5
161 6 300.1
159 6 401.5
12 66 377.1
156 7 153.3
42 108 440.2
135 9 108.6
19 153 98.8
43 53 225.4
44 53 225.4
133 13 367.8
190 12 308.7
50 18 321.6
-1.0
-0.5
0.0
0.5
1.0
Correlation coefficient

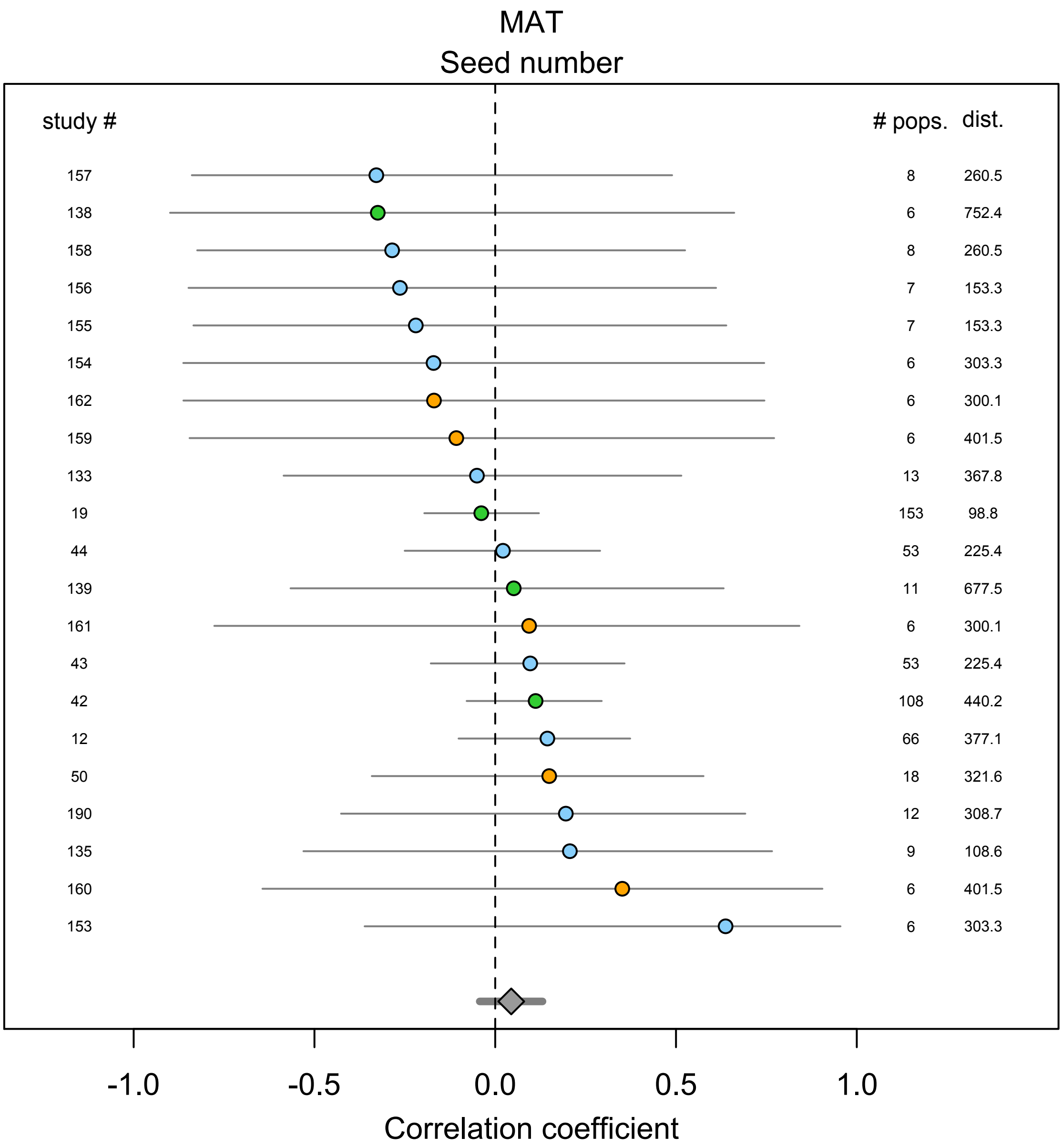
MAT
Seed number
study #
### pops.
dist.
157 8 260.5
138 6 752.4
158 8 260.5
156 7 153.3
155 7 153.3
154 6 303.3
162 6 300.1
159 6 401.5
133 13 367.8
19 153 98.8
44 53 225.4
139 11 677.5
161 6 300.1
43 53 225.4
42 108 440.2
12 66 377.1
50 18 321.6
190 12 308.7
135 9 108.6
160 6 401.5
153 6 303.3
-1.0
-0.5
0.0
0.5
1.0
Correlation coefficient

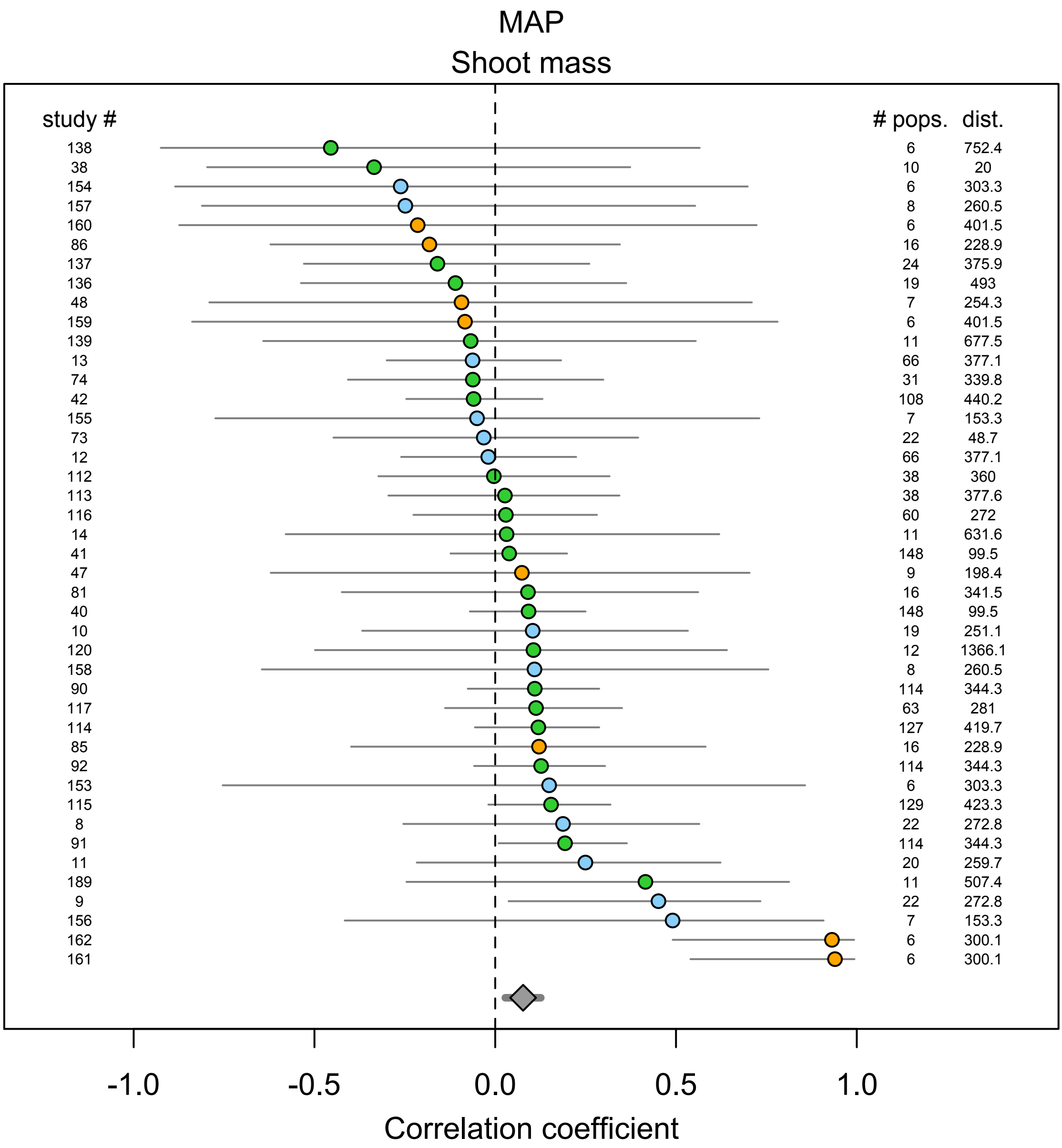
MAP
Shoot mass
study #
### pops.
dist.
138 6 752.4
38 10 20
154 6 303.3
157 8 260.5
160 6 401.5
86 16 228.9
137 24 375.9
136 19 493
48 7 254.3
159 6 401.5
139 11 677.5
13 66 377.1
74 31 339.8
42 108 440.2
155 7 153.3
73 22 48.7
12 66 377.1
112 38 360
113 38 377.6
116 60 272
14 11 631.6
41 148 99.5
47 9 198.4
81 16 341.5
40 148 99.5
10 19 251.1
120 12 1366.1
158 8 260.5
90 114 344.3
117 63 281
114 127 419.7
85 16 228.9
92 114 344.3
153 6 303.3
115 129 423.3
8 22 272.8
91 114 344.3
11 20 259.7
189 11 507.4
9 22 272.8
156 7 153.3
162 6 300.1
161 6 300.1
-1.0
-0.5
0.0
0.5
1.0
Correlation coefficient

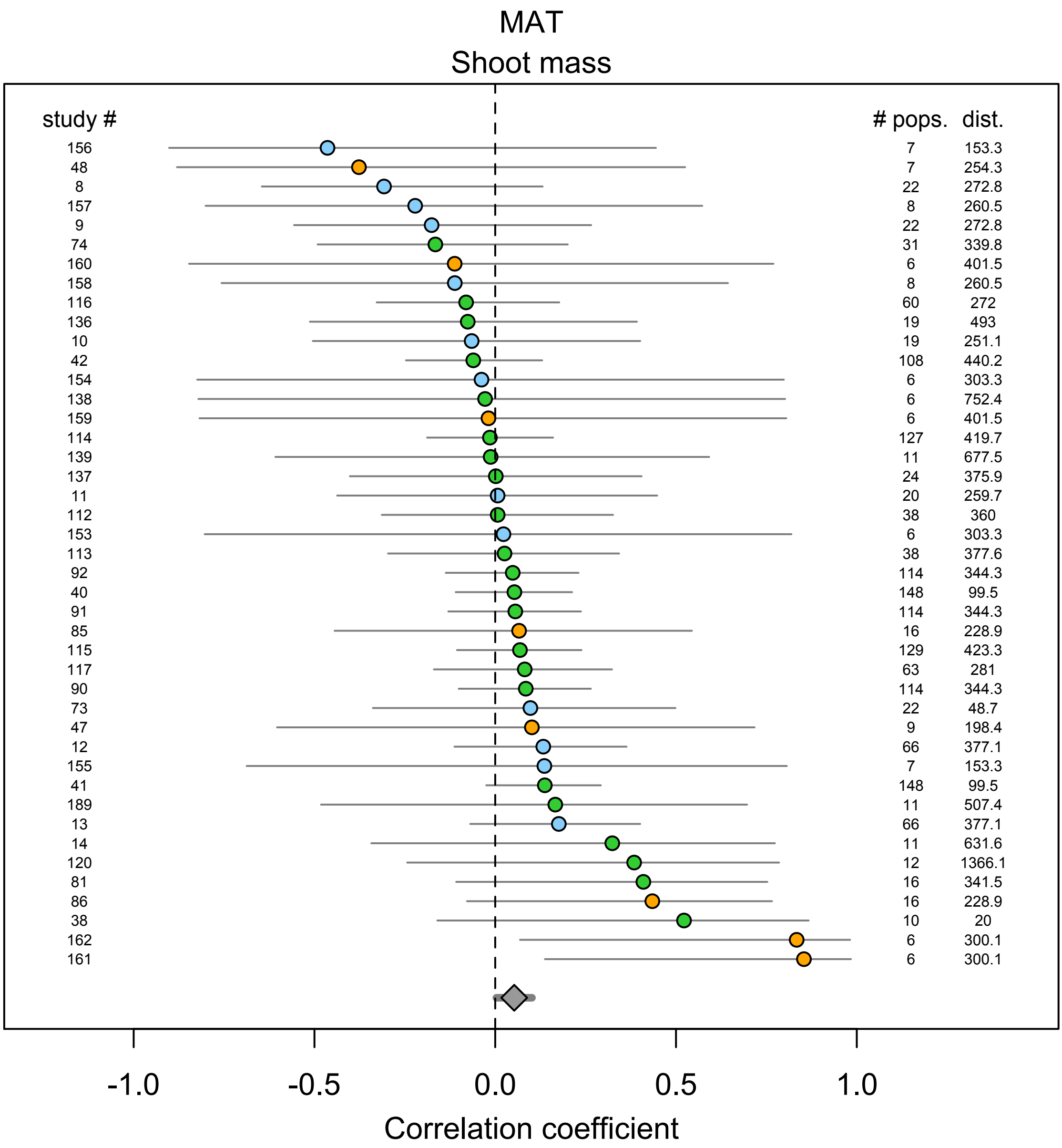

MAT
Shoot mass
study #
### pops.
dist.
156 7 153.3
48 7 254.3
8 22 272.8
157 8 260.5
9 22 272.8
74 31 339.8
160 6 401.5
158 8 260.5
116 60 272
136 19 493
10 19 251.1
42 108 440.2
154 6 303.3
138 6 752.4
159 6 401.5
114 127 419.7
139 11 677.5
137 24 375.9
11 20 259.7
112 38 360
153 6 303.3
113 38 377.6
92 114 344.3
40 148 99.5
91 114 344.3
85 16 228.9
115 129 423.3
117 63 281
90 114 344.3
73 22 48.7
47 9 198.4
12 66 377.1
155 7 153.3
41 148 99.5
189 11 507.4
13 66 377.1
14 11 631.6
120 12 1366.1
81 16 341.5
86 16 228.9
38 10 20
162 6 300.1
161 6 300.1
-1.0
-0.5
0.0
0.5
1.0
Correlation coefficient

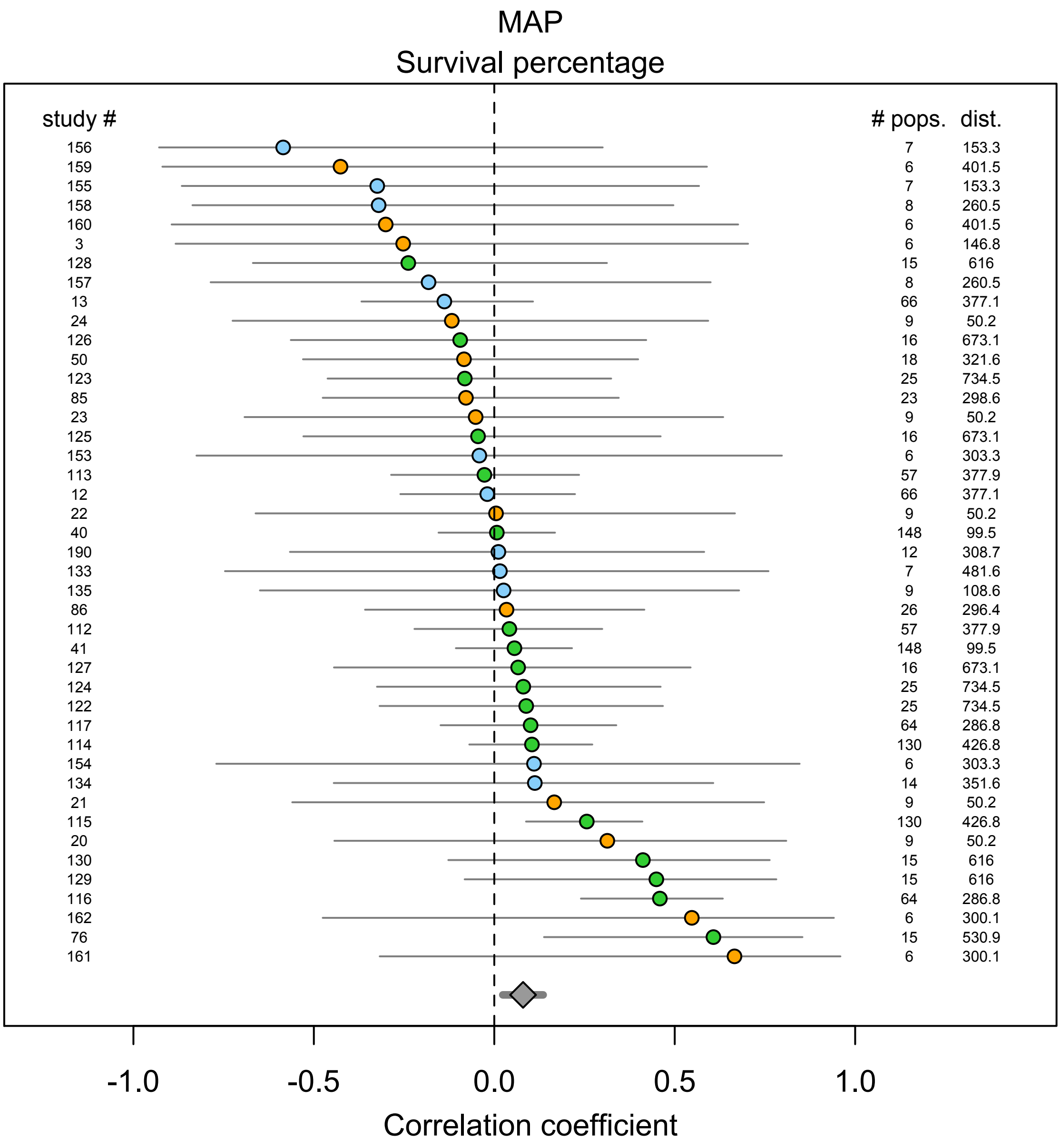
MAP
Survival percentage
study #
### pops.
dist.
156 7 153.3
159 6 401.5
155 7 153.3
158 8 260.5
160 6 401.5
3 6 146.8
128 15 616
157 8 260.5
13 66 377.1
24 9 50.2
126 16 673.1
50 18 321.6
123 25 734.5
85 23 298.6
23 9 50.2
125 16 673.1
153 6 303.3
113 57 377.9
12 66 377.1
22 9 50.2
40 148 99.5
190 12 308.7
133 7 481.6
135 9 108.6
86 26 296.4
112 57 377.9
41 148 99.5
127 16 673.1
124 25 734.5
122 25 734.5
117 64 286.8
114 130 426.8
154 6 303.3
134 14 351.6
21 9 50.2
115 130 426.8
20 9 50.2
130 15 616
129 15 616
116 64 286.8
162 6 300.1
76 15 530.9
161 6 300.1
-1.0
-0.5
0.0
0.5
1.0
Correlation coefficient

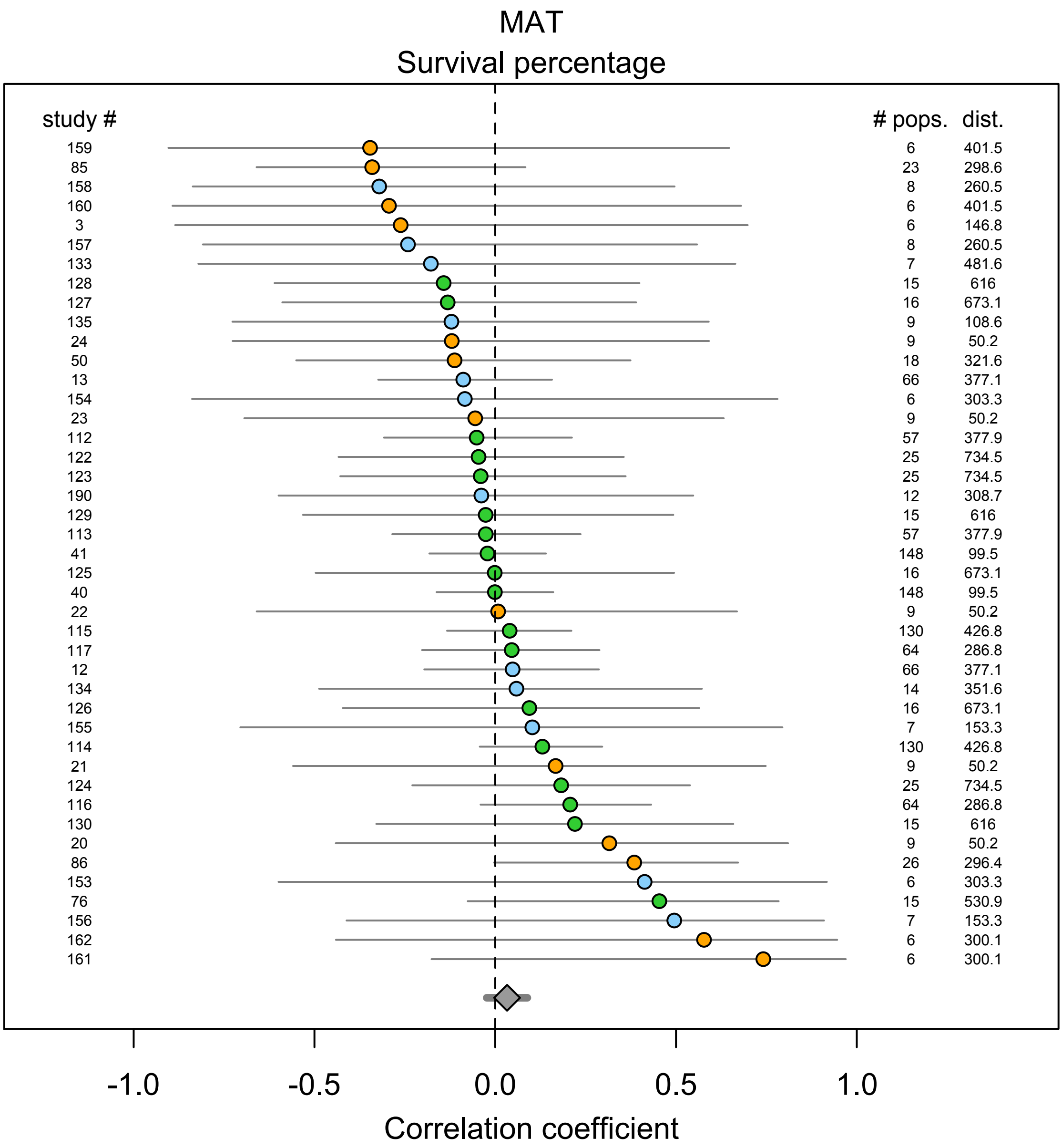
MAT
Survival percentage
study #
### pops.
dist.
159 6 401.5
85 23 298.6
158 8 260.5
160 6 401.5
3 6 146.8
157 8 260.5
133 7 481.6
128 15 616
127 16 673.1
135 9 108.6
24 9 50.2
50 18 321.6
13 66 377.1
154 6 303.3
23 9 50.2
112 57 377.9
122 25 734.5
123 25 734.5
190 12 308.7
129 15 616
113 57 377.9
41 148 99.5
125 16 673.1
40 148 99.5
22 9 50.2
115 130 426.8
117 64 286.8
12 66 377.1
134 14 351.6
126 16 673.1
155 7 153.3
114 130 426.8
21 9 50.2
124 25 734.5
116 64 286.8
130 15 616
20 9 50.2
86 26 296.4
153 6 303.3
76 15 530.9
156 7 153.3
162 6 300.1
161 6 300.1
-1.0
-0.5
0.0
0.5
1.0
Correlation coefficient
